# Comprehensive characterization of human color discrimination thresholds

**DOI:** 10.1101/2025.07.16.665219

**Authors:** Fangfang Hong, Ruby Bouhassira, Jason Chow, Craig Sanders, Michael Shvartsman, Phillip Guan, Alex H. Williams, David H. Brainard

**Affiliations:** Department of Psychology, University of Pennsylvania; Reality Lab Research, Meta; FAIR, Meta; Center for Neural Science, New York University; Center for Computational Neuroscience, Flatiron Institute

## Abstract

Color discrimination thresholds—the smallest detectable color differences—provide a benchmark for models of color vision, enable quantitative evaluation of eye diseases, and inform the design of display technologies. Despite their importance, a comprehensive characterization of these thresholds has long been considered intractable due to the psychophysical curse of dimensionality. Here, we address this challenge using a novel semi-parametric Wishart Process Psychophysical Model (WPPM), which leverages the feature that the internal noise limiting color discrimination varies smoothly across stimulus space. The model was fit to data collected with a non-parametric adaptive trial-placement procedure, enabling efficient stimulus selection. Together, through the combination of adaptive trial placement and *post hoc* WPPM fitting, we achieved a comprehensive characterization of color discrimination in the isoluminant plane with only ∼6,000 trials per participant (N = 8). Once fit, the WPPM allows readouts of discrimination performance for any stimulus pair. We validated these readouts against 25 probe psychometric functions, measured with an additional 6,000 trials per participant held out from model fitting. In conclusion, our study provides a foundational dataset for color vision, and our approach generalizes beyond color to any domain in which the internal noise limiting performance varies smoothly across stimulus space, offering a powerful and efficient method for comprehensively characterizing various perceptual discrimination thresholds.

## Introduction

Measurements of discrimination threshold—the smallest detectable stimulus change—are foundational for understanding biological vision. Threshold measurements support inferences about the neural mechanisms mediating performance (***Hecht et al., 1942; Campbell and Robson, 1968***), guide the design of displays and specification of perceptual tolerances (***MacAdam, 1942; de Lange Dzn, 1958***), allow quantitative evaluation of eye diseases (***Aspinall et al., 1983; Johnson et al., 2011; Niwa et al., 2014; Vemala et al., 2017***), inform models of supra-threshold perceptual representations (***Fechner, 1860; Hillis and Brainard, 2007; Zhou et al., 2024***), and allow perceptual effects to be incorporated into the study of cognitive processes (***Palmer et al., 1993; Najemnik and Geisler, 2005; Olkkonen et al., 2014***). Modern psychophysical methods (***Knoblauch and Maloney, 2012; Prins et al., 2016***) provide rigorous quantification of thresholds, and the theory of signal detection (***Green et al., 1966; Ashby and Soto, 2015; Hautus et al., 2021***) provides a mature framework for relating thresholds to the precision of the underlying representation.

Despite the central role of perceptual thresholds, characterization of thresholds has largely been limited to single stimulus dimensions. For example, pedestal functions characterize contrast discrimination thresholds across varying baseline contrasts (***Foley and Legge, 1981***). To generalize threshold characterization beyond a single dimension, we introduce the concept of the *psychometric field*: a multidimensional function that specifies the probability of a particular perceptual response as a joint function of both a reference and a comparison stimulus. In contrast to the psychometric function, which describes response probability as a function of variation around a fixed reference, the psychometric field captures how discrimination performance varies across all combinations of reference and comparison stimuli in a stimulus space. As the dimensionality of the psychometric field increases, the number of trials needed to tile the field grows exponentially—a psychophysical curse of dimensionality.

In this study, we focus on human color discrimination thresholds. Despite their significance and applications described above, fully characterizing human color discrimination—even on a single planar slice—has long been considered impractical (***Schrödinger, 1920***). This is because, although the stimulus space itself is two-dimensional, the underlying psychometric field is four-dimensional, as both the reference and comparison stimuli vary along two color dimensions. Mapping this field requires estimating discrimination performance across a densely sampled set of reference stimuli, with multiple comparison stimuli tested at each. The number of required trials quickly becomes intractable using conventional methods such as the method of constant stimuli (MOCS). While adaptive trial-placement procedures can greatly improve sampling efficiency (***Lesmes et al., 2010; Watson, 2017***), they typically rely on certain parametric forms. In many cases—including ours— such forms are not known in advance.

Here, we show that it is possible to obtain a comprehensive characterization of the color discrimination psychometric field in the isoluminant plane. We achieved this by efficiently sampling reference-comparison stimulus pairs obtained using a non-parametric adaptive trial-placement procedure (***Owen et al., 2021; Letham et al., 2022***), and then fitting the data *post hoc* with a semiparametric model that leverages the feature that the internal noise limiting color discrimination varies smoothly across stimulus space. We collected full datasets from 8 individual participants, and for each participant, we validated the accuracy of the model readouts against independent threshold measurements from held-out validation trials. Importantly, from the model fit, we can read out the psychometric function along any chromatic direction around any reference stimulus in the plane and thus determine the discrimination threshold in that direction. Our study provides a foundational dataset that can be used to test computational and neural models of color discrimination, benchmark color metrics, and develop models that can predict supra-threshold color discrimination performance.

## Results

### Overview

The Results section is organized as follows. We begin with a brief overview of the experimental stimuli and task (Task and stimuli), followed by a summary of how our model characterizes the full psychometric field (The Wishart Process Psychophysical Model (WPPM)) and a description of the non-parametric adaptive trial-placement procedure used to collect the data (Adaptively sampled trials). Having described these essential methods, we then present our core results (WPPM threshold estimates) and evaluate the validity of our model (Validation of the WPPM). Finally, we compare our findings with previous measurements from the color discrimination literature (Comparison with previous measurements). Additional technical details are provided in Methods and Materials and Appendix 1 - Appendix 12.

### Task and stimuli

Participants (N = 8) performed a 3AFC oddity task. On each trial, three blobby stimuli were shown in a triangular spatial arrangement—two identical reference stimuli and one comparison stimulus with a different surface color (***Figure 1***A). The comparison stimulus was pseudo-randomly assigned to one of the three positions. Participants were asked to identify the odd one out. Stimuli were rendered using the Unity graphics engine, and color was controlled by varying the specified surface reflectance using RGB (red, green, blue) coordinates, with other scene aspects held constant. We used naturalistic stimuli to increase the relevance of our results for understanding color vision in the real world. Hedjar and colleagues (***Hedjar et al., 2025***) provide a comparison of color discrimination using stimuli similar to ours versus traditional flat spatially uniform patches.

**Figure 1.**
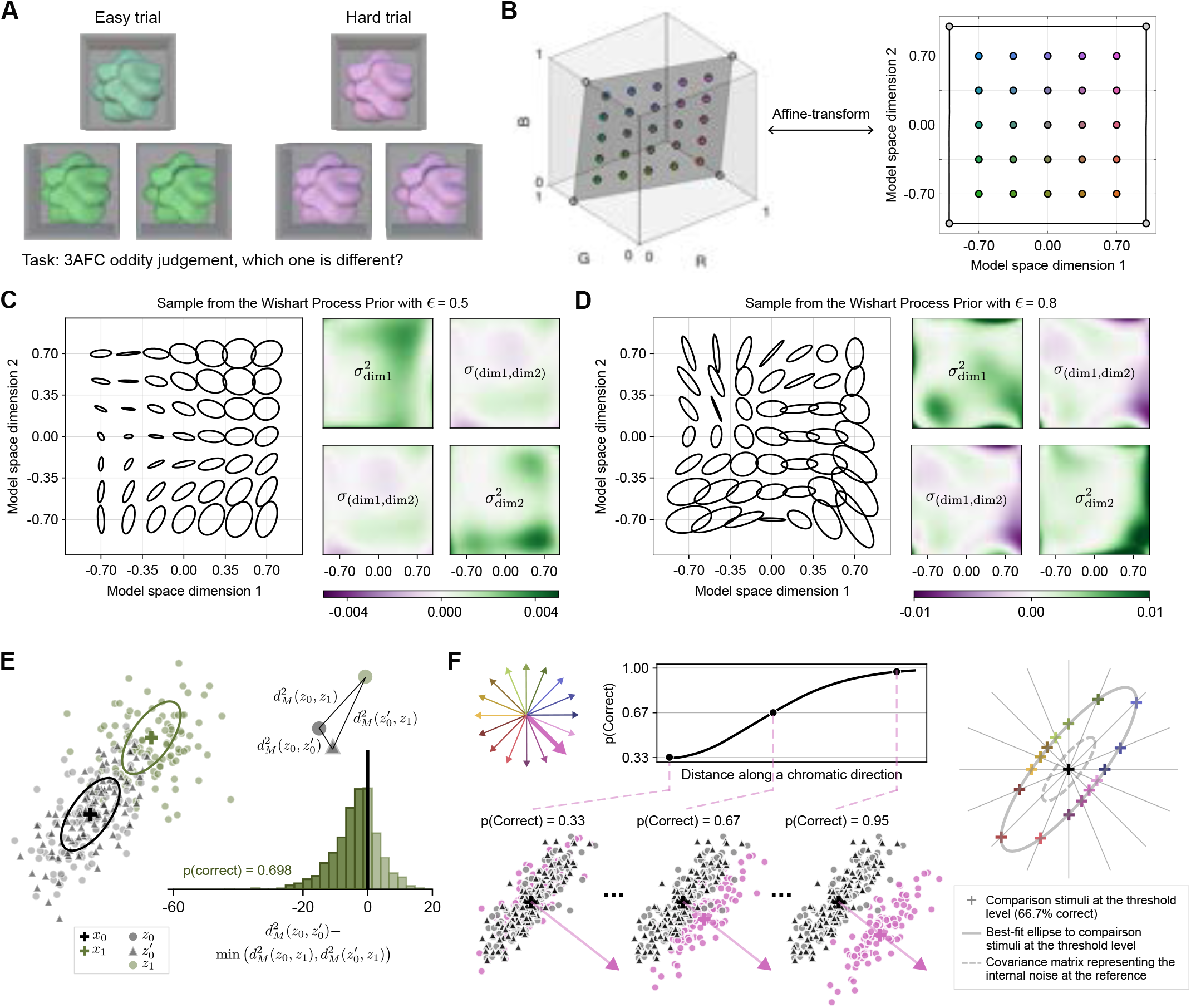
Task, stimuli and the WPPM. (A) 3AFC oddity task. On each trial, participants viewed a triplet of stimuli—two identical references and one different comparison—and identified the odd one out. (B) Stimuli are constrained to lie in the isoluminant plane that passes through the display’s gray point. Data are represented and fit in a transformation of this plane, which we refer to as *model space*. The grid of dots illustrates the transformation between the plane in the RGB and model space. (C) Example of a smoothly varying covariance matrix field produced by the WPPM. The field is generated by sampling from a finite-basis Wishart random process with a smooth prior (*ϵ* = 0.5 and *γ* = 0.0003; Prior over the weight matrix). Although the field is illustrated on a 7 × 7 grid, it specifies a covariance matrix 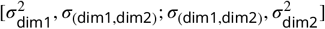 for every stimulus in the plane, as shown in the heat maps. (D) Example of a less smoothly varying covariance matrix field. This field is obtained by drawing from a finite-basis Wishart random process with a less smooth prior (*ϵ* = 0.8 and *γ* = 0.0003). (E) Observer model. For each stimulus triplet 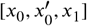, internal representations 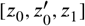 are drawn from multivariate Gaussian distributions centered at the reference stimulus with noise characterized by the corresponding covariance matrices. The model determines whether the observer correctly identifies the odd stimulus by comparing the squared Mahalanobis distances 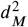 between the representation pairs. (F) Derivation of elliptical threshold contour. One-dimensional psychometric functions are approximated using Monte Carlo simulations (10,000 samples per stimulus pair shown for illustration; 2,000 used during model fitting). For each selected chromatic direction, we derive the threshold distance corresponding to 66.7% correct. An ellipse is then fit to the threshold distances to describe the discrimination threshold contour.

We made spectral calibration measurements (***Brainard et al., 2002***) to establish the relationship between RGB and the light emitted from the display. These measurements allowed us to represent the stimuli in terms of the excitations of the human L, M, and S cones elicited by the stimuli, and more generally in any standard color space (***Brainard, 1996, 2003; Brainard and Stockman, 2010***). For this study, stimuli were constrained to lie in the isoluminant plane passing through the monitor’s gray point and bounded by its gamut. This plane was then affine-transformed into a square ranging between -1 and 1 along each axis (***Figure 1***B; Appendix 1). We refer to the space in which the transformed plane is as the *model space* because it is directly related to the way we formulated our semi-parametric model, and also served as a convenient representation for the non-parametric adaptive trial-placement procedure we used.

### The Wishart Process Psychophysical Model (WPPM)

As an overview of our modeling approach, we fit the color discrimination responses (coded as ‘correct’ or ‘incorrect’) with a novel model, the WPPM—a Bayesian probabilistic model that combines an observer model (specified through a likelihood function) with an expectation of smoothness in the internal noise limiting color discrimination (specified through a prior distribution). Once fit to the data, the WPPM yields a continuously varying field of covariance matrices that characterize the internal noise in the perceptual representation of color stimuli (***Figure 1***C-D). These covariance matrices, in turn, determine the entire psychometric field.

More specifically, we designed the observer model within the WPPM to formalize the intuition that the stimulus perceived as the most distant from the other two is identified as the “odd one out”. The internal representation of each stimulus is assumed to be noisy and modeled as a multivariate Gaussian with the same dimensionality as the stimulus space. We assume the mean of each distribution is given by the corresponding stimulus’ location in the model space. In contrast, we allow the covariance matrices to vary across the model space to account for differences in the encoding precision of the stimuli. Because discrimination thresholds depend on the relative size of signal change and internal noise, an alternative formulation could instead attribute threshold variation across stimulus location to nonlinearities in signal encoding while assuming constant internal noise, or to a mixture of nonlinearities and stimulus-varying noise (***Zhou et al., 2024***). Our formulation should be understood as characterizing the signal-to-noise properties that limit discrimination and not a commitment to a particular interpretation of how these properties arise.

On each trial, the observer model has access to one sample from the distribution of each of the three stimuli—two identical reference stimuli and one comparison. The observer model computes the pairwise squared Mahalanobis distance between each pair of noisy samples, using the weighted average of the covariance matrices of the reference and comparison (***Figure 1***E). By using Mahalanobis distance to make decisions (instead of, for example, Euclidean distance), the observer accounts for the expected noise structure. The two stimuli whose pairwise distance is smallest are identified as the references, and the remaining stimulus as the comparison (the “odd one out”). Because there is no simple closed-form solution for this decision rule (***Mullen and Ennis, 1991***), we used Monte Carlo simulation to approximate the percent-correct performance (Observer model).

We expect the internal noise that limits color discrimination to vary smoothly across the model space—that is, small changes in the reference stimulus should produce only small changes in the corresponding internal noise. The WPPM reflects this expectation by placing a finite-basis Wishart process prior over the continuous field of covariance matrices (***Wilson and Ghahramani, 2011***). Intuitively, the Wishart process prior introduces a regularization term to the model—it penalizes rapid variation in the covariance matrix field. The strength of smoothness is controlled by two hyperparameters of the model, *ϵ* and *γ* (***Figure 1***C-D; Prior over the weight matrix).

To fit the model to each participant’s data, we found the *maximum a posteriori* (MAP) estimates of the WPPM’s parameters, using gradient-based numerical optimization of the log posterior density, that is, the sum of the log prior density and log likelihood function (Model fitting).

The best-fit model parameters, together with the observer model, allow us to read out percent-correct performance for any pair of reference and comparison stimuli. In particular, to read out a one-dimensional psychometric function, we select a reference stimulus and use the observer model to approximate performance as the comparison stimulus varies along a line (***Figure 1***F, left panels). The threshold distance along the line is defined as the distance that yields 66.7% correct. By repeating this process across many directions, we derive a set of threshold distances around the reference (***Figure 1***F, right panel). Given our assumption that internal noise follows a multivariate Gaussian distribution, these threshold distances form approximately elliptical contours, which we fit with ellipses for visualization. This approach is consistent with prior work showing that ellipses provide a good approximation of color discrimination thresholds (***MacAdam, 1942; Brown and MacAdam, 1949; Noorlander et al., 1981, 1983; Poirson and Wandell, 1990; Krauskopf and Gegenfurtner, 1992; Knoblauch and Maloney, 1996; Danilova and Mollon, 2025***), despite some reported deviations (***Newton and Eskew, 2003; Shepard et al., 2016, 2017***). Notably, while we show threshold contours corresponding to 66.7% correct for visualization, once fit, the WPPM allows us to read out the full psychometric function for any reference and chromatic direction—effectively mapping the entire psychometric field. Given that the psychometric field is derived from the underlying field of covariance matrices that characterize internal noise, the smoothness constraint imposed on the covariance matrices naturally propagates to the threshold contours and the field itself.

### Adaptively sampled trials

Reference and comparison stimuli for each trial were selected using AEPsych (***Owen et al., 2021***), an open-source package for adaptive psychophysics. For the adaptive sampling model, we used a probit-Bernoulli Gaussian Process (GP) model (***Williams and Rasmussen, 2006***) with a radial basis function (RBF) kernel. As with the WPPM, the GP assumes smooth variation in performance across the model space due to the RBF kernel, but unlike the WPPM, it does not impose any specific parametric form on the internal noise (or thresholds). The semi-parametric constraint— multivariate Gaussian-shaped internal noise—was introduced only when fitting the WPPM. For this reason, we describe the adaptive trial-placement procedure as non-parametric (relative to the WPPM)—acknowledging that while it incorporates some parametric assumptions, they are less restrictive than those of the WPPM. This non-parametric approach ensures that our data collection was not biased by assuming the correctness of the WPPM prior to validation.

Each participant completed 6,000 AEPsych-driven trials: the first 900 were generated using quasi-random Sobol’ sampling (***Sobol, 1967***) to provide an adequate initialization for the GP; for the remaining 5,100 trials, the GP was updated continuously based on participants’ responses, and each trial was adaptively selected to be most informative for estimating the thresholds targeted at 66.7% correct (***Letham et al., 2022***) (***Figure 2***A, S1, S2).

**Figure 2.**
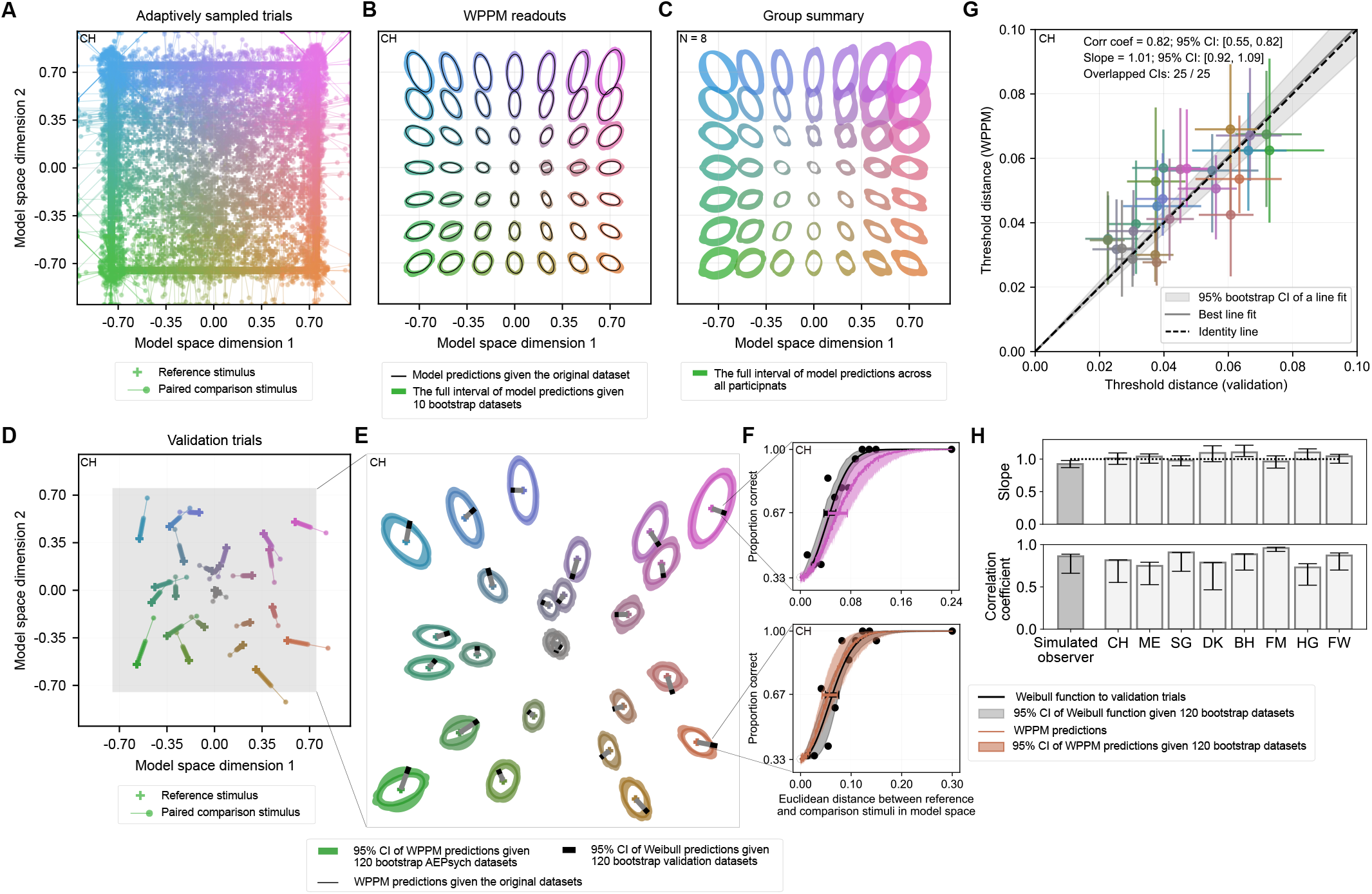
Threshold results and validation. (A) Adaptively sampled trials. AEPsych-driven stimulus pairs are adaptively sampled for estimating thresholds across the psychometric field. Of the 6,000 trials, the first 900 are Sobol’-sampled; the remaining 5,100 (shown here) are adaptively selected using the EAVC acquisition function, based on a non-parametric GP model that is updated every 20 trials. (B) Discrimination threshold contours (66.7% correct) read out from the WPPM on a grid of reference stimuli for a representative participant, based on fits to the 6,000 AEPsych trials. (C) Group summary of WPPM readouts (N = 8), evaluated on the same grid of reference stimuli. (D) Validation trials for the same participant. The validation conditions (reference stimuli and chromatic directions along which the comparison stimulus varies) are randomly generated for each participant (see Appendix 4 for validation conditions used for the remaining participants). (E) Comparison of thresholds. Ellipses represent discrimination threshold contours read out from the WPPM fit (same fit as in panel B), evaluated at the 25 reference stimuli used in the validation trials. Gray lines: the validation directions; black bars: the 95% bootstrapped confidence intervals for the corresponding validation thresholds. (F) Comparison of psychometric functions. Only two validation conditions are shown for illustration (see Appendix 4.1 for all 25 conditions for each participant). (G) Linear regression of thresholds predicted by the WPPM against validation thresholds for the same participant. Horizontal and vertical error bars represent 95% confidence intervals for the validation thresholds and the WPPM predictions, respectively. (H) Summary of regression slopes and correlation coefficients for all participants. Error bars: 95% confidence intervals. As a benchmark, the same analysis is performed on a dataset simulated using a ground truth WPPM instance that approximates CIELab Δ*E*94 (Appendix 5).

Adaptive sampling with AEPsych requires solving two optimization problems: one for updating the GP model (***Williams and Rasmussen, 2006***) and another for selecting the next trial using the Expected Absolute Volume Change (EAVC) acquisition function (***Letham et al., 2022***). To reduce computational time, we updated the GP model only every 20 trials. Sometimes, however, either the fitting or the trial selection process did not complete in time for the upcoming stimulus presentation. To avoid perturbing the participants’ rhythm, in these cases we slotted in pre-generated fallback trials (Appendix 3). The number of fallback trials varied across participants, ranging from 0 to 466 ***Figure S1***). These trials were included along with the 6000 AEPsych-driven trials when fitting the WPPM.

### WPPM threshold estimates

For each participant, we fit the WPPM to the 6,000 AEPsych-driven trials, along with any additional fallback trials. To visualize the fits, we read out the elliptical threshold contours around a grid of reference stimuli (***Figure 2***B for a representative participant). The threshold contours revealed three key regularities: (1) thresholds were lowest for references near the achromatic point defined by the background behind the blobby stimuli, (2) thresholds increased with the distance of the reference from the achromatic point, and (3) the major axes of the elliptical threshold contours tend to be radially oriented with respect to the achromatic point. These regularities are consistent with previous results in the color discrimination literature, as explained further in Comparison with previous measurements.

The data were broadly consistent across participants, in the sense that the three regularities noted above are observed in the individual participant data (Appendix 2.1). In the model space representation, individual variability was lowest near the achromatic point where sensitivity was highest and increased with the distance between the reference and the achromatic point (***Figure 2***C). Specifically, this variation was quite large in the upper right quadrant of the model space, where ellipse orientations varied considerably. This variation in orientation was also apparent when examining the data in other colorimetric representations (Appendix 7–Appendix 8). Increases in interparticipant variability with increasing thresholds have been observed in other perceptual discrimination tasks (***Girshick et al., 2011; Aguilar et al., 2017; Hong et al., 2021***).

### Validation of the WPPM

To validate the WPPM estimates, we interleaved 6,000 validation trials throughout the experiment. These trials were held out from WPPM model fitting. For each participant, we used Sobol’ sampling to select 25 reference stimuli and associated chromatic directions, with a unique draw per participant. Along each sampled chromatic direction, we used MOCS to sample 12 comparison levels: 11 were evenly spaced, and one was selected to provide easily discriminable catch trials (***Figure 2***D). The comparison levels were selected based on a pilot dataset to account for variability in thresholds across different reference stimuli and chromatic directions (see Design for details). Notably, we intentionally avoided densely sampling around a small number of references to minimize differential perceptual learning between the trials used for fitting the WPPM and those reserved for validation (***Horiuchi and Nagai, 2024***).

For each of the 25 validation references, we fit a Weibull psychometric function to the 240 MOCS trials collected along the sampled chromatic direction and identified the comparison stimulus corresponding to 66.7% correct (see two examples in ***Figure 2***F). We then used the WPPM fit (constrained by non-overlapping trials) to extract elliptical contours corresponding to the 66.7% threshold level for each validation reference (***Figure 2***E). To compare the WPPM and validation thresholds, we read out the WPPM threshold along the MOCS chromatic direction for each reference. The 95% bootstrapped confidence intervals for the WPPM estimates overlapped with those from the Weibull fits in all 25 conditions for participant CH (***Figure 2***E) and in 22 to 25 conditions across other participants (Appendix 4.1). These results demonstrate a high degree of agreement between thresholds derived from the WPPM psychometric field and the 25 discrete MOCS validation thresholds. This agreement indicates that the Wishart process prior we imposed did not lead to substantial over-smoothing, as the validation thresholds were estimated independently without a smoothness constraint. Also notable is that the size of the 95% bootstrapped confidence intervals for the WPPM and validation thresholds was similar (***Figure 2***F, G; Appendix 4.1).

To quantify the agreement, we performed a linear (slope only) regression between the thresholds read out from the WPPM fit and those obtained using the validation trials (***Figure 2***G). The results further support agreement between the two sets of estimates (mean correlation coefficient = 0.84, range = 0.73–0.96). For 7 out of 8 participants, the regression slope was not significantly different from 1 (mean slope = 1.04, range = 0.96–1.10) (***Figure 2***H; see Appendix 4.1 for comparisons for each participant). To assess whether there were more subtle sources of bias not captured by the regression slope, we analyzed the residuals—the differences between the WPPM and validation thresholds. While we found no evidence that residuals depended on the orientation or shape of threshold contours read out from the WPPM fit, we did observe one small but statistically significant relationship: the model slightly overestimated thresholds when validation thresholds were low and underestimated them when validation thresholds were high (slope = −0.176, *t*(198) = −5.727, *p* < 0.001, *R*^2^ = 0.142). However, the magnitude of this bias was small (Appendix 4.2).

As an additional benchmark, we simulated trials and responses from a ground-truth WPPM instance chosen to approximate the CIELab Δ*E*94 metric, and fitted the model to the simulated data (Appendix 5). This allowed us to assess the ability of the WPPM to recover simulated ground truth, which is not possible with human data. The readout threshold ellipses based on the WPPM fit are in good agreement with the ground truth (***Figure S16***C). We then conducted the same validation analyses on the simulated data as described above. The thresholds read out from the WPPM fit agreed with 23 of the 25 validation thresholds, based on overlapping confidence intervals. A linear regression yielded a correlation coefficient of 0.86 and a slope of 0.92—well within the confidence intervals observed in participants’ data (***Figure 2***H, left bar). Residual analysis revealed a negative correlation between residuals and the magnitude of the ground-truth validation thresholds (Appendix 5.6; ***Figure S14***), consistent with trends observed in human participants. With the simulated data, however, we can interpret the magnitude of this bias in the context of the agreement with ground truth and conclude that it is small (***Figure S16***C). Access to ground truth also provides us with additional ways to visualize patterns in the bias (Appendix 5.7).

Taken together, these results validate the accuracy of the WPPM and highlight the remarkable efficiency of our approach. With 6,000 trials, conventional psychophysical methods only allowed us to estimate percent-correct performance along one chromatic direction across 25 references. In contrast, our new approach—combining a non-parametric, adaptive trial placement with *post hoc* fitting of the semi-parametric WPPM—allowed us to map the entire psychometric field, providing the percent-correct performance for any reference-comparison stimulus pair in the isoluminant plane, using the same number of trials.

### Comparison with previous measurements

The WPPM is equivariant under affine transformations of color space (Appendix 1.4), allowing threshold contours derived in our model space to be transformed into other colorimetric representations. This flexibility enables direct comparisons with color discrimination thresholds reported in the literature. At the outset, we emphasize that the size and shape of threshold contours depend on ancillary experimental factors, including task design, stimulus spatial and temporal properties, and participants’ state of adaptation. Given these differences, we do not expect quantitative agreement across studies. Nonetheless, such comparisons help set our findings in the context of the literature. To illustrate, we present several such comparisons below, in the colorimetric representations used in the original studies.

We first compared the overall pattern of threshold variation in the isoluminant plane with measurements made by MacAdam using the method of adjustment (***MacAdam, 1942***) (Appendix 6). In his seminal work, the ellipses do not represent discrimination thresholds *per se*, but rather the standard deviation of color matches for each reference stimulus. Nevertheless, we consider his measurements to be comparable to ours, based on the linking assumption that discrimination thresholds are proportional to the internal noise that governs the variability of the appearance-based matches (***Crozier and Holway, 1937***). We observed a similar global structure in how the orientation and scale of the ellipses vary with reference stimulus. As expected, the absolute sizes of the threshold contours differ between studies (***Figure 3***A; MacAdam ellipses magnified by 10× while ours magnified by 2×). In addition to differences in task and stimulus spatial and temporal structure, it is worth noting that in MacAdam’s experiment, participants controlled the stimulus duration themselves (***Wandell, 1985***) and their state of adaptation differed considerably across reference stimuli (***Krauskopf and Gegenfurtner, 1992***). Despite these differences, the general correspondence between the datasets is apparent. It is also noteworthy that MacAdam’s results are based on 25,000 adjustments at a limited number of reference locations, whereas our ∼6,000 forced-choice responses enabled us to characterize discrimination performance across all in-gamut reference–comparison pairs in the isoluminant plane.

**Figure 3.**
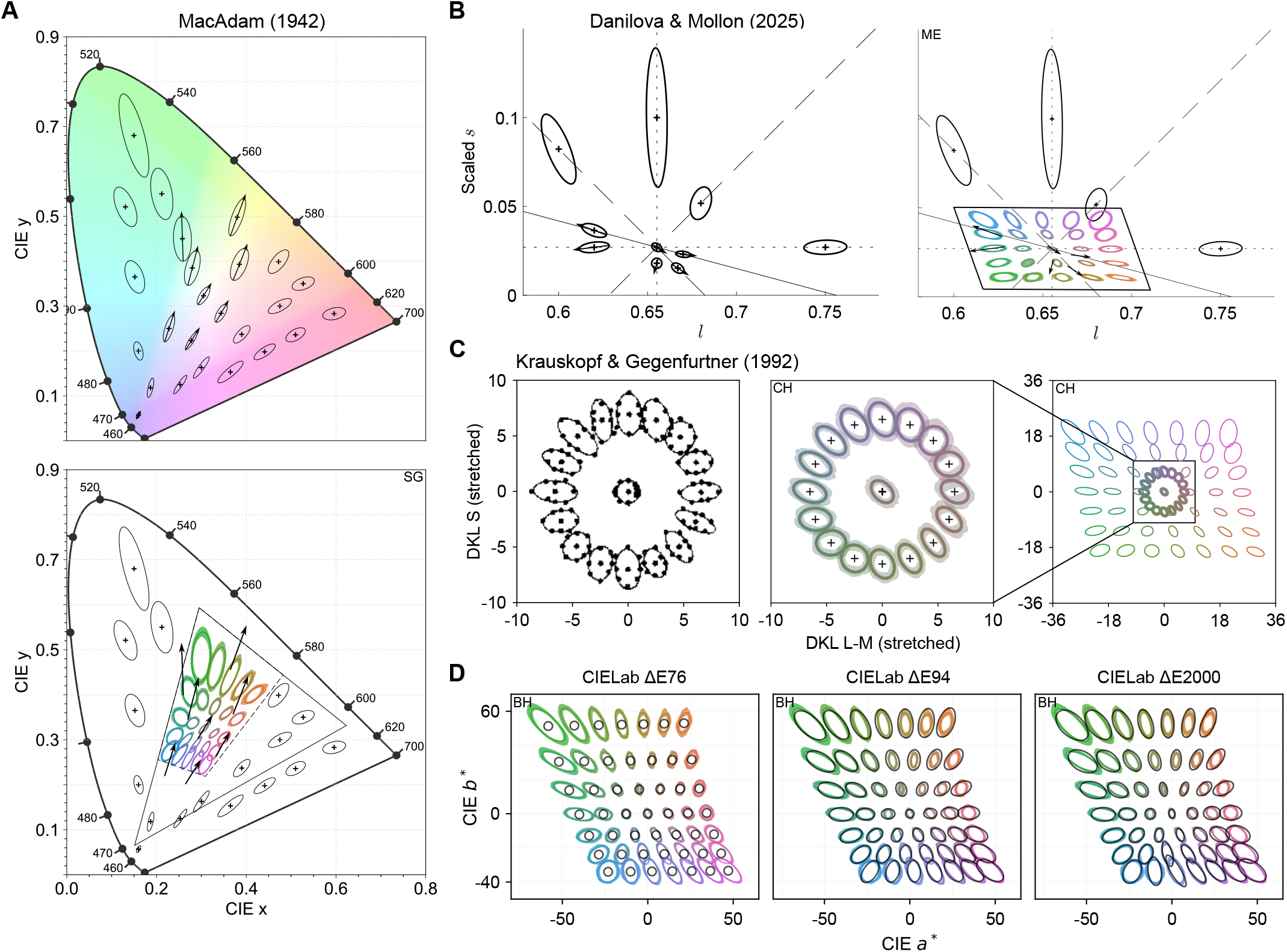
Comparison of color discrimination thresholds with previous measurements. Across all panels, black contours represent thresholds from prior studies, whereas colored contours represent the 66.7% discrimination thresholds estimated in our study. Colored shaded regions indicate 95% confidence intervals computed from 120 bootstrapped datasets. (A) ***MacAdam 1942***.Top panel: MacAdam’s original threshold ellipses, magnified 10× for visualization. Bottom panel: Threshold contours measured from one participant in our study and transformed from the model space into the CIE 1931 chromaticity diagram. Reference stimuli are sampled from a 5 × 5 grid spanning [−0.7, 0.7] along each dimension of the model space. To reduce visual clutter, MacAdam ellipses falling within the gamut of our isoluminant plane (parallelogram) are shown only by arrows indicating their major axes. For visual comparability, our ellipses are magnified 2× to approximately match the scale of MacAdam’s data. The triangle indicates our monitor’s gamut. (B) ***Danilova and Mollon 2025***. Left panel: Original threshold contours (79.4% correct) from their study, magnified by 4×. Right panel: Threshold contours from one participant in our study, transformed from the model space into the scaled MacLeod–Boynton space used in their study. Reference points are sampled on a 5 × 5 grid ranging from –0.7 to 0.7. As in panel A, to reduce visual clutter, their ellipses that fall within the gamut of our isoluminant plane (parallelogram) are shown as black arrows indicating only their major axes. For visual comparability, our ellipses are magnified 1.5×. (C) ***Krauskopf and Gegenfurtner 1992***. Left panel: 79.4% threshold contours (Fig. 14 from their study, reproduced under Creative Commons CC BY-NC-ND 4.0). Right panel: 66.7% threshold contours from one participant in our study, transformed into DKL space with the axes scaled for each participant to equate thresholds along the L-M and S axes at the adapting chromaticity, as was done in their study. All contours are shown at their original sizes in this scaled representation. (D) CIELab Δ*E*76, Δ*E*94, and Δ*E*00. Threshold is defined as Δ*E* = 2.5, chosen to approximately match the scale of our measured thresholds, which are shown at their original sizes. See Appendix 6 - Appendix 9 for additional details.

In a more recent study, ***Danilova and Mollon 2025*** measured threshold contours across a relatively broad region of the isoluminant plane, with sparse sampling of reference stimuli. The experimental paradigm in their study closely resembled ours: both used a fixed adapting point—D65 in their case and the monitor gray point in ours—and employed an oddity task to estimate discrimination thresholds. They used a 4AFC design combined with an adaptive staircase procedure, whereas we used a 3AFC version. To compare our data with theirs, we transformed our discrimination threshold contours read out from the WPPM fit into the same scaled MacLeod–Boynton space (***MacLeod and Boynton, 1979***) used in their study (Appendix 7). Despite methodological and stimulus differences, our results replicated the overall pattern of variation in ellipse orientation and size across the color space. In particular, thresholds were smallest near the adapting point, increased with distance from it, and the ellipses generally pointed toward the adapting point. We observe closer agreement in the absolute sizes of the threshold contours than when comparing with MacAdam’s data (***Figure 3***B; their ellipses were magnified by 4×, ours by 1.5×). An interesting commonality between our data and those of ***Danilova and Mollon 2025*** is the rotation of the ellipses at the adapting point relative to the axes of the MacLeod-Boynton space, a rotation seen in all of our participants (***Figure S21***). See their discussion of this rotation for possible mechanistic interpretations.

In the next comparison, we turned to the study by ***Krauskopf and Gegenfurtner 1992***, whose measurements were concentrated within a small region near the achromatic point. Their experiment used a fixed adapting point and a 4AFC oddity task, with individual thresholds estimated using a three-down-one-up staircase procedure. To enable direct comparison, we read out threshold contours from our model at the same set of reference stimuli they used. Their results revealed two key features: (1) the threshold contour was smallest at the adapting point, and (2) as the reference moved away from it, the contours generally became increasingly elongated along the axis pointing toward the adapting point. Both features were observed in our data albeit with some interparticipant variability (***Figure 3***C; Appendix 8). Our measurements differ from theirs, however, in the orientation of the ellipse at the adapting chromaticity: in our data, the ellipses are rotated with respect to the DKL axes for all our participants.

Lastly, we compared our results with iso-distance contours obtained with different versions of the CIELab Δ*E* color difference metrics (***CIE, 2004***). Although Δ*E* metrics were developed to describe supra-threshold perceptual color differences for a stimulus configuration that differs from ours, comparisons with threshold-level measurements are of interest—particularly because of the widespread use of Δ*E* to equate perceptual differences in studies of cognitive processes (***Winawer and Witthoft, 2023; Garside et al., 2025***). To derive a threshold contour for any given reference stimulus, we identified the comparison stimuli corresponding to Δ*E* = 2.5 across multiple chromatic directions and fit an ellipse. While the choice of Δ*E* = 2.5 is arbitrary, it primarily affects the overall size of the contour rather than its shape. The comparison reveals that the iso-distance contours of the original CIELab Δ*E*76, which remains widely used, bear little resemblance to our threshold contours (***Figure 3***D, left panel; Appendix 9). The large deviations we observed between Δ*E*76 and our data provide further caution against the practice of using Δ*E*76 to predict perceptual color difference. In contrast, the more recent Δ*E*94 and Δ*E*00 metrics provided a much closer match (***Figure 3***D, center and right), with only modest deviations from our measurements. These deviations may arise from differences between threshold and supra-threshold perceptual judgments, as well as from discrepancies in experimental conditions between our study and those used to constrain the parameters of the CIELab Δ*E* metrics. An important feature of our data is that it enables such comparison with any perceptual metric across the isoluminant plane.

## Discussion

### A data-efficient approach for characterizing color discrimination thresholds

In this study, we demonstrated a data-efficient approach for achieving a comprehensive characterization of human color discrimination thresholds. Participants performed a 3AFC oddity task and completed 6,000 trials that were specifically targeted near threshold via a non-parametric adaptive trial-placement procedure (***Owen et al., 2021; Letham et al., 2022***). We then developed and fit a novel WPPM to these adaptively sampled trials (along with a small number of fallback trials). The WPPM defines a continuous mapping from each reference stimulus to its associated internal noise, characterized by a covariance matrix. This mapping, in turn, enables predictions of discrimination performance for any pair of reference and comparison stimuli—effectively mapping out the full four-dimensional psychometric field. To evaluate model validity, we interleaved 6,000 additional validation trials to estimate 25 probe psychometric functions. The results revealed that thresholds read out from the WPPM closely matched those derived from the validation trials, supporting the model’s accuracy. Thus, by combining the non-parametric adaptive trial-placement procedure with *post hoc* fitting of the semi-parametric WPPM, we achieved an unprecedentedly comprehensive characterization of color discrimination in the isoluminant plane.

Our measurements align qualitatively with previous studies that used either sparse sampling or targeted at a small region of a color space (***MacAdam, 1942; Krauskopf and Gegenfurtner, 1992; Danilova and Mollon, 2025***). Moreover, our measurements provide a more comprehensive characterization, in that the WPPM allows direct readout of a threshold contour at any reference stimulus without the need for additional measurement. Additionally, for studies examining how thresholds vary with factors such as stimulus size, presentation duration, or adaptation state, our approach offers a scalable and data-efficient approach for measuring how these factors affect the psychometric field. Finally, we have performed simulations and collected preliminary data that indicate it will be feasible to fully characterize the color discrimination psychometric field across the three-dimensional gamut of our display (***Hong et al., 2026***), a goal that has been previously described as “hopelessly difficult” (***Schrödinger, 1920***).

### Prior specification

A key assumption of the WPPM is that internal noise varies smoothly across the stimulus space. This smoothness assumption is implemented through a prior on the variance of the weights applied to the model’s basis functions (Prior over the weight matrix). The smoothness prior plays a nontrivial role in the final WPPM estimates and therefore the choice of prior required careful selection.

Our cross-validation analyses indicate that the smoothness hyperparameters that characterize the prior we used in the main analyses fall within a regime that balances over-smoothing against excessive uncertainty in the estimates (Appendix 10.1). When the smoothness embodied in the prior is too strong, the model produces overly uniform threshold estimates that fail to capture structure in the data. When the prior smoothness is too weak, the estimates become more variable. Consistent with this bias–variance tradeoff, agreement between WPPM and validation thresholds starts off low with strong smoothness, increases as the smoothness constraint is relaxed, and declines when smoothness becomes too weak (Appendix 10.3). These two analyses narrow the range of sensible hyperparameter values and provide support for the hyperparameter values (*ϵ* = 0.4; *γ* = 0.0003) adopted in the main analyses.

As a general matter, determining appropriate prior hyperparameter values can be challenging for interpreting data using Bayesian models. This problem is at the heart of empirical Bayesian approaches to data analyses, in which prior hyperparameters are estimated from the data, and then the data are analyzed with these estimates (***Efron, 2012***). A full empirical Bayesian approach of this sort currently exceeds what we can compute under reasonable time constraints. In our experience, evaluating model performance across a range of the hyperparameters using cross-validation helps identify the region that balances over-smoothing against excessive uncertainty in the estimates, while the inclusion of validation trials helps identify the regime that maximizes agreement between WPPM predictions and validation thresholds.

The number of basis functions included in the model constitutes an additional modeling choice. To evaluate this choice, we examined the fitted weights as a function of the order of the Chebyshev polynomials and found that they decay to near zero at the highest polynomial order used in the model. This indicates that including additional basis functions would not materially affect the inferred psychometric field, given our hyperparameter values (***Figure S4***).

### Implications for the mechanisms of color perception

Consistent with a well-established body of evidence, we found that thresholds were smallest near the achromatic reference, reflecting heightened sensitivity at the adapting point (***Craik, 1938; Brown, 1952; Hurvich and Hurvich-Jameson, 1961; Pointer, 1974; Loomis and Berger, 1979; Krauskopf and Gegenfurtner, 1992***). In addition, threshold contours were oriented toward the achromatic center, in agreement with previous findings (***Krauskopf and Gegenfurtner, 1992; Gegenfurtner, 2025; Danilova and Mollon, 2025***). Although the WPPM characterizes the data in terms of stimulus-dependent noise, the fitted psychometric field can be used to evaluate mechanistic models that posit specific transformations between stimuli and their internal representations.

The observation that the size and orientation of the elliptical threshold contours vary with the reference stimulus rules out mechanistic models that posit a linear transformation of cone excitations into three post-receptoral channels followed by fixed additive noise. Such models predict identical ellipses across the stimulus space. Moreover, the observation that the orientation of the elliptical threshold contours changes across reference stimuli also rules out mechanistic models in which a linear transformation of cone excitations is followed by limiting noise applied independently to each of the three channels. These models allow variation in the lengths of the major and minor ellipse axes, but predict that the orientation of these axes will be the same for all reference stimuli.

Cone-opponent models that posit noise and nonlinearities at multiple stages of processing, possibly with an over-complete cone-opponent representation to capture parallel channels along the visual pathways, may be able to account for the observed data, as may models that invoke higher-order mechanisms (e.g. mechanisms narrowly tuned for hue). For more on relevant ideas see ***Wyszecki 1982; Wandell 1995; Chen et al. 2000; Eskew Jr 2009; Stockman et al. 2010; Hansen and Gegenfurtner 2013; Shevell and Martin 2017***. Notably, mechanistic models are often tested using additional manipulations such as adaptation and noise-masking; our approach can be extended to incorporate manipulation of such factors (***Zhang et al., 2026***), as well as of stimulus spatial and temporal structure and retinal location.

We studied a relatively young cohort of eight participants and found broadly consistent patterns across individuals, while also observing individual differences. Individual differences have provided valuable insights into the mechanisms of color vision (***Bosten, 2022***) and are also of interest for understanding how much any given individual is likely to differ from an average characterization. A successful mechanistic model should allow investigation of whether our observed individual differences can be attributed to individual variation in biological factors known to influence color vision, for example pre-retinal absorption, photopigment spectral sensitivity, and the ratio of L to M cones in the mosaic (***Neitz and Jacobs, 1986; Brainard et al., 2000; Kremers et al., 2000; Carroll et al., 2002; Hofer et al., 2005; Bosten, 2022; Rezeanu et al., 2023***).

#### Extensions of the WPPM framework

To enable studies involving larger and more diverse populations, further improvements in the efficiency of our approach are likely achievable. In the present study, we used a non-parametric adaptive trial-placement procedure to avoid biasing data collection by assuming the correctness of the WPPM. Given the validation of the WPPM presented here, future studies could instead incorporate adaptive trial-placement strategies tailored to the model, thereby improving efficiency. Alternatively, one could leverage the current dataset to develop stronger priors that capture the regularities we observe and use these priors to guide more efficient trial placement. Stronger priors could also increase the quality of the estimates available from a fixed set of trials, although care should be taken to ensure that the prior does not overly constrain the estimates. Another approach is to develop mechanistic models with relatively few parameters, which could be estimated efficiently using parametric adaptive sampling (***Watson, 2017***). Finally, a complementary strategy is to increase the rate at which participants provide information about thresholds through more efficient psychophysical experimental paradigms (***Agosti et al., 2026; Barnett et al., 2025; Burge and Cormack, 2024***).

Another aspect of the WPPM framework that can be adapted for different applications is the mapping between stimulus space and model space, as well as the choice of basis functions. In the present work, we defined the model space to be bounded within [−1, 1] for mathematical convenience, as this is the domain on which our chosen basis functions—the 2D Chebyshev polynomials— are defined. These basis functions could in principle be replaced by alternatives such as Zernike (***Zernike, 1934; Thibos et al., 2000***) or sinusoidal (Fourier) basis functions (***Stein and Shakarchi, 2011***), which may be better suited for stimulus domains with disc-shaped geometries or without clear boundaries. The number of basis functions can also be adjusted based on prior knowledge about the expected smoothness of the psychometric field for a given stimulus domain. More generally, other approaches to leveraging the smoothness of psychophysical performance and physiological response have been developed (***Gravesen, 2015; Waz et al., 2025; Rad and Paninski, 2010; Savin and Tkacik, 2016***), including recent work on color discrimination (***Koenderink et al., 2026***).

### Toward a metric of supra-threshold color difference

A longstanding and fundamental question in vision science is whether it is possible to develop a perceptual metric that accurately predicts both threshold-level and supra-threshold judgments of color difference. For example, considerable effort has gone into attempts to find color representations where the perceptual color difference between two color stimuli is predicted by the Euclidean distance of their coordinates in the representation (e.g., the original 1976 CIELab and CIELuv Δ*E* metrics; see ***Brainard 2003; Robertson et al. 1977***). Our measurements directly establish a locally Euclidean metric for threshold-level differences. While threshold behavior is well described as locally Euclidean, supra-threshold judgments have been shown to violate the assumptions of a globally Euclidean geometry (***Wuerger et al., 1995; Ennis and Zaidi, 2019***). In particular, perceptual similarity judgments at larger distances often fail to satisfy key Euclidean properties such as the expectation that variability in judgments should increase with Euclidean distance (***Wuerger et al., 1995***), and that a stimulus equidistant from two endpoints should be perceived as equally similar to both (***Ennis and Zaidi, 2019***).

An alternative framework, originally proposed by Fechner (***Fechner, 1860***) and explored subsequently (***Schrödinger, 1920; Macadam, 1979; Wyszecki, 1982; Zaidi, 2001; Koenderink, 2010; Bujack et al., 2022; Roberti, 2024; Stark et al., 2025***), suggests that supra-threshold differences may be understood as the accumulation of small threshold-level differences along a path between stimuli. In this framework, color space is taken to be a Riemannian manifold—a space that is locally Euclidean but may be globally curved. The perceptual distance between two colors is hypothesized to correspond to the geodesic—the shortest path between them in terms of accumulated thresholds. This distance is computed by integrating local thresholds along all possible paths between the two points and selecting the path with the smallest total. In our observer model, this integration is effectively equivalent (up to a constant) to summing internal noise along the path.

Testing this *geodesic hypothesis* requires knowledge of how internal noise (or thresholds) varies across color space, as this determines the geodesics. Our measurements provide the necessary knowledge for the isoluminant plane, enabling direct empirical tests of the geodesic hypothesis within this slice of color space, as well as elaborations of this hypothesis (***Bujack et al., 2022; Stark et al., 2025***). The results of such tests may depend on the particular experimental paradigms used to assess supra-threshold perceptual differences.

Because there is no guarantee that the geodesics between two stimuli in the isoluminant plane are themselves confined to this plane within the full three-dimensional color space, testing the geodesic hypothesis in this plane based on our current data would be considered provisional. Nonetheless, such tests would provide valuable exploration of the perceptual geometry revealed by our measurements. As noted above, our approach makes it feasible to extend the measurements to the full three-dimensional color space (***Hong et al., 2026***), which, when completed, will allow subsequent investigations to overcome this limitation.

It is possible that the geodesic hypothesis—and more generally the idea that threshold-level judgments can predict supra-threshold judgments—will fail. Nonetheless, we view understanding whether, when, and how such failures occur as central to guiding development of a successful account of supra-threshold color difference perception.

### Beyond color discrimination

Our approach is generalizable to a wide range of perceptual tasks. A key insight that makes comprehensive characterization of human color discrimination thresholds feasible is the assumption— shared by both our model and the models implemented in AEPsych—that internal noise, and thus thresholds, vary smoothly across stimulus space. This smoothness assumption is not unique to color perception; it applies broadly to other domains. Indeed, smoothly varying elliptical or ellipsoidal thresholds have been reported in studies of motion perception (***Reisbeck and Gegenfurtner, 1999; Champion and Freeman, 2010***), auditory speed discrimination (***Freeman et al., 2014; Carlile and Leung, 2016; Bertonati et al., 2021***), motion-in-depth (***Wardle and Alais, 2013***), and numerosity perception (***Cicchini et al., 2016, 2019, 2023***). These parallels highlight the broader relevance of our framework and suggest that combining the non-parametric adaptive trial-placement procedure with the WPPM could be a powerful strategy for characterizing perceptual limits across diverse domains.

## Methods and Materials

### Preregistration

This study was preregistered at a public repository. As described in the preregistration document, exploratory analyses were conducted on data from one participant (CH) prior to preregistering an initial hyperparameter choice of *ϵ* = 0.5 for the main analysis. After data collection completed, we performed the hyperparameter sweeps (Appendix 10). These led to our final choice of *ϵ* = 0.4 and *γ* = 0.0003.

### Participants

Eight participants (six female, aged 22–30 years; seven right-handed) were recruited for the study. Six were paid volunteers who were naive to the purpose of the study. The remaining two were experimenters and participated without additional compensation. All participants had normal or corrected-to-normal vision (20/40 or better in each eye, assessed using a Snellen eye chart) and normal color vision (assessed using Ishihara plates). The study was approved by the Institutional Review Board at University of Pennsylvania, and written informed consent was obtained from all participants prior to the experiment.

### Apparatus

Stimuli were presented using an Alienware computer (Aurora R11) running Windows 10 Enterprise, equipped with Intel Core− i7-10700K processor and NVIDIA GeForce RTX 3080 GPU. The display was a DELL U2723QE monitor (59.8cm width, 33.6 cm height, 3840 × 2160 resolution, 60 Hz refresh rate). The monitor was positioned 189 cm from the chinrest, subtending a visual angle of 18.0 × 10.2 degrees of visual angle (dva). Monitor color and luminance measurements were obtained with a Klein K-10A colorimeter and a SpectraScan PR-670 radiometer. The display resolution was approximately 200 pixels/dva, above the typical human foveal resolution limit.

The Alienware computer was used solely for stimulus presentation, whereas adaptive sampling of the stimuli was performed on a separate custom-built PC with a high-performance Gigabyte motherboard (X299X aorus master), an NVIDIA GeForce RTX3070 GPU and a 12-core Intel i9-10920X processor. This computer also ran Windows Enterprise 10. The two computers communicated via a shared network disk, using a custom protocol based on text files that both computers could read and write.

A USB speaker (3 Watts output power, 20k Hz frequency response) was used for playing auditory feedback, and a gamepad controller (Logitech Gamepad F310) was used for registering trial-by-trial responses.

### Stimulus

The visual scene (***Figure S33***A) was constructed in Unity (v2022.3.24f1) and rendered using its standard shader. The scene consisted of three identical blobby 3D objects, each created in Blender (v4.0) with a matte, non-reflective surface. On each trial, the surface color of the blobby objects was varied by adjusting their RGB values in Unity. The three blobby objects (2.5 × 2.5 dva; 203,900 pixels each) were arranged in a triangular configuration (***Figure 1***A). Each blobby object was centered and floating inside its own cubic room (3.3 × 3.3 dva; *x* = 0.302, *y* = 0.322, *Y* = 66.1 *cd*/*m*^2^). Each room, along with the blobby stimulus inside it, was illuminated exclusively by an achromatic spotlight positioned in front of the object and set to maximum intensity (*R* = *G* = *B* = 1). The three rooms were presented against a spatially uniform gray background (18.0 × 10.2 dva; *x* = 0.306, *y* = 0.326, *Y* = 116.8 *cd*/*m*^2^). The centers of the blobby objects were 3.7 dva apart.

### Calibration and color depth

We used a SpectraScan PR-670 to measure the monitor’s primaries and gamma function as rendered through Unity (Appendix 11.1). These measurements directly characterized the relationship between the specified RGB values for the blobby stimuli and the light emitted from the display. The same calibration was repeated for all three blobby stimuli, confirming consistent color behavior across screen locations. Based on these results, a single gamma correction—derived from the bottom-right stimulus—was applied to all three objects during the experiment. This correction was validated by remeasuring the output with gamma correction applied, showing good alignment with the predicted identity line. To confirm stability over time, we repeated the calibration one month into data collection and observed negligible changes.

Additionally, we used a Klein K-10A colorimeter to verify that the system achieved sufficient color depth. For this check, a single blobby stimulus was presented at the center of the screen, rather than in the full triangular arrangement. Measurements confirmed that Unity and our video chain were able to produce at least 12-bit color precision via its native 8-bit output and implicit spatial dithering that occurred across the surface of the blobby object through the rendering process (Appendix 11.2).

### Design

We restricted our stimuli to lie within the isoluminant plane that passes through the monitor’s gray point (i.e., R = G = B = 0.5). To define the boundaries of this plane, we identified the limits of RGB values that remained within the monitor’s gamut. These boundary points formed a parallelogram in RGB space. We then computed an affine transformation that maps this parallelogram onto a square bounded within [−1, 1] (Appendix 1). The forward and inverse transformations enabled conversion between RGB and the model space: stimuli were rendered in RGB space, while trial placement and model fitting were performed in the model space.

We used AEPsych (v0.7) to sample a total of 6,000 reference–comparison stimulus pairs. The first 900 trials were generated using Sobol’ sampling (***Sobol, 1967***), a “space-filling” design based on a low-discrepancy quasi-random sequence. The remaining 5,100 trials were adaptively selected to efficiently estimate thresholds across the entire psychometric field. Each stimulus pair was defined in the 2D model space. As a result, the psychometric field comprised four variables: two specifying the reference stimulus, *x*_0_ ∈ ℝ^2^, and the other two specifying a difference vector, Δ ∈ ℝ^2^, which was added to the reference to define the comparison stimulus *x*_1_ = *x*_0_ + Δ. Reference values were constrained between [−0.75, 0.75] along each model dimension. Each element of Δ was constrained between [−0.25, 0.25] to ensure that all comparison stimuli remained within the [−1, 1]^2^ bounds of the model space. During the initial 900 Sobol’-sampled trials, the difference vector Δ was scaled by one of three factors (1/4, 2/4, or 3/4) before being added to the reference stimulus. This controlled the distance between the reference and comparison stimuli, effectively modulating task difficulty. These scaling factors were evenly distributed and pseudo-randomized across trials. For the remaining 5,100 trials, all four variables were adaptively selected using AEPsych’s optimization procedure. Specifically, the underlying GP model was updated every 20 trials, and new trials were selected using the Expected Absolute Volume Change (EAVC) acquisition function (***Letham et al., 2022***), targeting the 66.7% threshold level across the entire psychometric field.

In addition to the 6,000 AEPsych-driven trials, we interleaved an additional 6,000 validation trials sampled using MOCS. Each participant was tested on 25 reference stimuli: one was fixed at the achromatic point and the remaining 24 were Sobol’-sampled within the isoluminant plane bounded within [−0.6, 0.6] along each model dimension. For each reference, a chromatic direction was Sobol’-sampled between 0^°^ and 360^°^. Each validation condition consisted of 12 stimulus levels: 11 equally spaced along the sampled direction and one easily discriminable level, with each level repeated 20 times. These levels were determined based on a pilot dataset described in the preregistration documents.

The validation trials were pre-generated for each participant, pseudo-randomized so that every 300 validation trials contained all the unique trials (25 conditions × 12 levels). To minimize differential learning effects between AEPsych-driven and validation trials, we pre-generated a randomized sequence in which the two trial types were arranged in alternating pairs, with the order within each pair shuffled. However, because AEPsych occasionally required longer time to determine the next trial placement, this sequence could not always be followed in real time. For this reason, we implemented a fallback trial strategy (Appendix 3): if, for any trial, AEPsych did not have trial placement computed in time, the next validation trial was inserted to keep the experiment moving. If necessary, subsequent validation trials were queued, but this was capped to a lead of four validation trials ahead of AEPsych trials. Once the cap was reached and AEPsych was still not ready, pregenerated fallback trials were presented instead. These fallback trials were Sobol’-sampled with the difference vector Δ scaled by one of three factors (2/8, 3/8, or 4/8) to manipulate task difficulty. Validation trials resumed once AEPsych caught up. Notably, the fallback trials (range: 0–466) were included alongside the 6,000 AEPsych trials when fitting the WPPM.

### Procedure

Participants performed a 3AFC oddity task. Each trial began with a fixation cross presented at the center of the screen for 0.5 s, followed by a blank screen for 0.2 s. Then, three blobby stimuli appeared inside the cubic rooms for 1 s. After participants responded, a blank screen was shown for 0.2 s, followed by auditory and visual feedback indicating accuracy (“correct” with a beep or “incorrect” with a buzz). Each trial was separated by a 1.5 s inter-trial interval. Participants were instructed that they could move their eyes freely during the stimulus presentation, but should maintain fixation while the fixation cross was on the screen.

The majority of the participants (7 out of 8) completed a total of 12 sessions. Each session began with 40 practice trials to familiarize participants with the task. This was followed by 1,000 experimental trials, consisting of 500 AEPsych-driven trials, 500 predetermined validation trials, and a small number of fallback trials. The validation trials were randomized and the two trial types were fully intermixed. Participants took a break every 200 trials. Each session took approximately 1.5 hours to complete. In total, those seven participants completed between 12,256 and 12,466 trials, depending on the number of fallback trials inserted. Participant CH completed 12,000 trials across 10 sessions, without fallback trials implemented. As a result, the inter-trial interval was slightly longer for this participant, but we expected this to have a negligible effect on performance.

### The Wishart Process Psychometric Model

Our implementation of the WPPM relies on two core assumptions about color perception: (1) internal noise that limits color discrimination follows a multivariate Gaussian distribution, centered at the reference stimulus, with a covariance matrix that captures both the magnitude and directional structure (i.e., size and orientation) of the noise, and (2) the covariance matrix varies smoothly across the model space, without abrupt local discontinuities. In the following subsections, we describe the WPPM in five parts. First, we define the observer model, which predicts percent-correct performance for a given pair of reference and comparison stimuli by modeling both the noisy internal representations and the decision rule. Second, we describe how we use a finite-basis Wishart process to parameterize the entire field of covariance matrices across the model space, along with the factors that control its smoothness. Third, we describe the weak prior imposed on the covariance matrix field to favor smooth variation. Fourth, we explain how, given a specification of the covariance matrix field, we compute the likelihood and thereby the posterior probability of the model and its free parameters given binary (correct or incorrect) color-discrimination responses. Finally, we show how, once the model is fit, the covariance matrix for any reference-comparison stimulus pair can be read out and combined with the observer model to predict percent-correct performance, including threshold contours around any reference stimulus.

#### Observer model

On each trial, the observer is presented with two identical reference stimuli, denoted *x*_0_, and one comparison stimulus, denoted *x*_1_ = *x*_0_ + Δ where Δ represents a small offset from the reference. Our model assumes that these three stimuli are independently encoded into an internal representational space by a noisy process, which we assume to follow a multivariate Gaussian distribution. Formally,

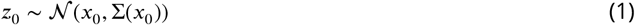

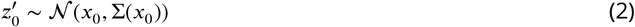

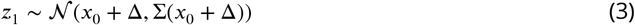

where 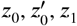 denote the internal representations derived from the two reference and the comparison stimuli, respectively. Our model posits that the observer correctly identifies *z*_1_ as representing the comparison stimulus (i.e. the “odd-one-out”) if

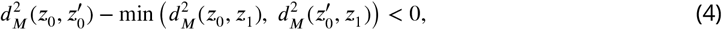

where 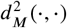 denotes the squared Mahalanobis distance for a selected pair of internal representations, formulated as

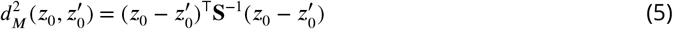

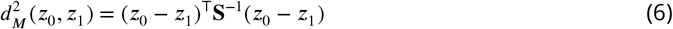

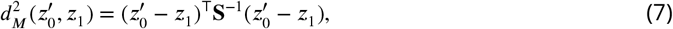

where **S** is the weighted average of the covariance across the reference and the comparison stimuli, that is,

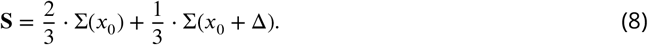

This decision rule is consistent with an observer that uses distances between internal representations to judge stimulus similarity (***Churchland, 1986***). We approximated the percent-correct performance using (N=2,000) Monte Carlo simulations (***Figure 1***E) as the closed-form analytical solution is complicated to derive (***Ennis and Mullen, 2014***). In each Monte Carlo simulation, we draw samples according to ***Equation 1*** - ***Equation 3*** and the outcome is marked as correct if the condition in ***Equation 4*** is fulfilled. The proportion of correct outcomes in the Monte Carlo simulation defines the model’s predicted percent-correct performance, which is then used to evaluate the likelihood function as explained in Model fitting.

#### Covariance matrix field

The WPPM specifies a covariance matrix at any selected reference stimulus across the entire isoluminant plane. Each matrix specifies the perceptual noise in terms of the variance along the two model dimensions 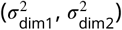 and their covariance (*σ*_dim1,dim2_) (***Figure 1***C-D).

The covariance matrix field is constructed using one-dimensional Chebyshev polynomial basis functions (***Chebyshev, 1853***). We chose Chebyshev polynomials because they allow for the expression of smoothness over a bounded interval without imposing periodic boundary conditions. Let*x* = [*x*_dim1_, *x*_dim2_] denote a location in the 2D model space. The basis functions are defined recursively for each model space dimension as given here for *x*_dim1_:

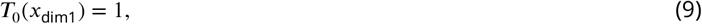

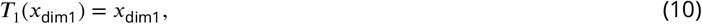

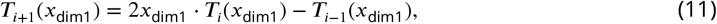

where *x*_dim1_, *T*_*i*_(*x*_dim1_) ∈ ℝ^*n*^, and *n* is the number of discretized points along that stimulus dimension, which can be chosen flexibly to achieve any desired resolution. We construct two-dimensional basis functions by taking the outer product:

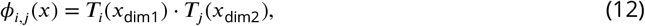

where *ϕ*_*i,j*_ ∈ ℝ^*n*×*m*^, with *n* = *m* representing the number of discretized points along each dimension of the model space. We limited the number of basis functions to five per dimension, i.e., *i, j* ∈ {0, 1, …, 4}, resulting in a total of 5 × 5 = 25 2D basis functions (***Figure 4***, first panel). The polynomial order of each 2D basis function is given by *i* + *j*, with higher-order basis functions describing more rapidly varying patterns.

**Figure 4.**
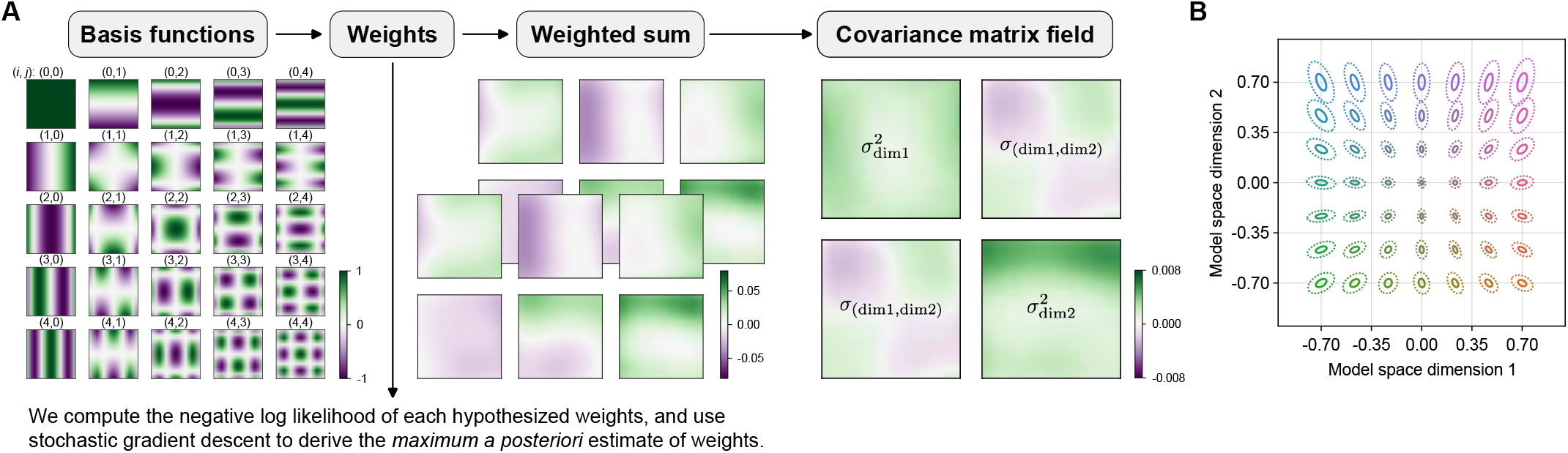
The finite-basis Wishart Process Psychophysical Model (WPPM). (A) Model overview. In our implementation, we use a set of 5 × 5 two-dimensional Chebyshev polynomial basis functions, denoted *ϕ*_*i,j*_ (*x*), where *i, j* ∈ {0, 1, …, 4}. These basis functions are summed using a learnable weight matrix **W** to produce an overcomplete representation **U**_*k,l*_(*x*), where *k* ∈ 1, 2 and *l* ∈ 1, 2, 3. The resulting representation **U**_*k,l*_ is then combined with its own transpose to produce a field of symmetric, positive semi-definite covariance matrices. Each matrix specifies the internal noise in terms of the variance along the two model dimensions 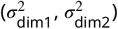 and their covariance (*σ*_dim1,dim2_). For this example covariance matrix field, the weights used to generate the field correspond to the best-fitting parameters for participant CH (see ***Figure S3*** for all participants). (B) Model readouts. Internal noise can be read out anywhere in the model space, illustrated here on a grid of 7 × 7 reference stimuli (solid lines), from which threshold contours (dashed lines) can be derived.

The basis functions were weighted by a learnable parameter matrix, **W** ∈ ℝ^5×5×2×3^, where the first two dimensions index the Chebyshev basis functions along each model space dimension (*i, j* ∈ {0, 1, …, 4}), and the last two dimensions index the output components (*k* ∈ {1, 2} and *l* = {1, 2, 3}). The weighted basis functions are expanded into an overcomplete representation **U**_*k,l*_ ∈ ℝ^*n*×*m*^ (***Figure 4***, second panel) as the following,

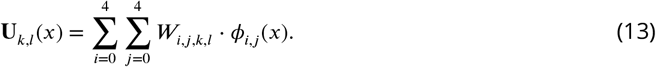

This weighted sum overcomplete representation was then combined with its own transpose to yield a positive semi-definite covariance matrix (***Figure 4***, third panel), Σ(*x*) ∈ ℝ^2×2^ for *x* at any discretized point in the model space, that is,

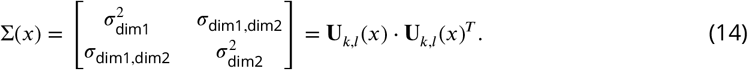

Notably, in our implementation, rather than matching the dimensionality of the intermediate representation *U* to that of Σ, we adopt an overcomplete parameterization motivated primarily by practical considerations. When we restricted the dimensionality indexed by *l* to 2, the optimization occasionally became ill-conditioned, leading to singular or unstable solutions. Expanding *l* to 3 substantially improved numerical stability and made the fitting procedure more robust. Increasing *l* beyond 3, however, would introduce additional degrees of freedom. We therefore selected *l* = 3 as a compromise between model flexibility and numerical stability. Regardless of whether the representation is square or overcomplete, the resulting matrices are symmetric and positive semidefinite.

The weight matrix serves as the free parameters of the model, controlling the smoothness of the covariance matrix field. The model is highly flexible, capable of generating a wide range of covariance matrix fields, from smooth to rapidly varying fields (***Figure 1***C-D).

#### Prior over the weight matrix

We imposed a weak prior over the weight matrix **W**. Specifically, we assumed that each weight was distributed *a priori* as a zero-mean one-dimensional Gaussian,

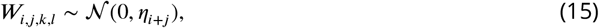

where *η* represents the variance of each weight and it decays exponentially with *i*+*j*, which denotes the polynomial order of the corresponding 2D basis function, that is,

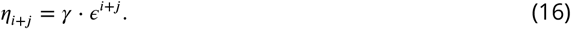

The hyperparameter *γ* controls the overall amplitude of the variance. The hyperparameter *ϵ* controls the rate at which the prior variance decays with increasing polynomial order. A higher value of *γ* or *ϵ* results in a prior that favors more rapidly varying covariance matrix fields, while a lower value favors smoother fields. By setting *γ* = 0.0003 and *ϵ* = 0.4, we adopted a prior that favors relatively smooth variation across the space (***Figure 4***, fourth panel).

#### Model fitting

We computed the negative log-likelihood of any hypothesized weight matrix given the participant’s binary responses *y*_*r*_ as follows:

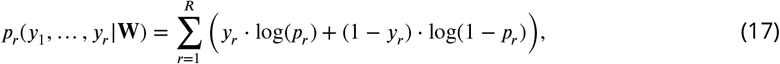

where *y*_*r*_ ∈ {0, 1} indicates whether the response on trial *r* was correct (1) or incorrect (0), *R* is the total number of trials used to fit the WPPM. The model-predicted accuracy *p*_*r*_ for each trial is given by:

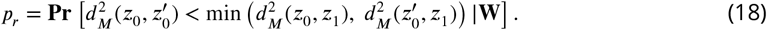

Note that on the *r*^th^ trial, *z*_0_, 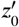, and *z*_1_ are internal representations that depend on the reference and comparison stimuli (*x*_0_ and *x*_1_) for that trial. For notational simplicity, the subscript *r* is omitted here.

Since we imposed a prior on the covariance matrix field to reflect the expectation of smooth variation, we combined the likelihood (***Equation 17***) and the prior (***Equation 16***) to calculate the posterior probability of **W**. As there is no simple closed form expression for *p*_*r*_, we resorted to a numerical approximation based on Monte Carlo simulations. The numerical approximation we built was differentiable with respect to the covariance matrix field, which enabled us to use gradient descent to maximize the posterior probability of **W** (see details in Appendix 12).

Notably, the factorization in ***Equation 14*** is not unique; multiple choices of *U* and consequently of *W* can yield the same covariance matrix. This non-uniqueness reflects the overcomplete parameterization and does not affect the uniqueness of the resulting covariance matrices or the corresponding threshold readouts of the model.

#### Psychometric field

For any given reference stimulus, the WPPM allows readouts of percent-correct performance along any chromatic direction, which in turn allows us to construct a threshold contour. We sampled comparison stimuli along 16 chromatic directions and simulated internal representations to estimate percent-correct performance, yielding a psychometric function for each direction (***Figure 1***F). The threshold distance in each direction was defined as the comparison stimulus corresponding to 66.7% correct. Collectively, these threshold distances form a contour that closely resembles an ellipse, with only minor deviations due to inhomogeneous internal noise between the reference and comparison stimuli. However, because the stimuli are locally proximal in the model space, such discrepancies are negligible. We therefore fit an ellipse to these points as a practical approximation. As a way of visualizing the psychometric field, we plot these ellipses—each corresponding to 66.7% threshold level—at a grid of reference locations. We emphasize, however, that the WPPM provides the full four-dimensional psychometric field, enabling readouts of the psychometric function along any chromatic direction for any reference stimulus within the model space.

### Data analysis

Color calibration analyses were performed using MATLAB 2023b. We computed inverse gamma lookup tables from the measured gamma functions (Appendix 11) and derived transformation matrices to convert values from the model space to RGB space (Appendix 1). Stimulus presentation, including gamma correction, was implemented in Unity, coded in C#.

All analyses were conducted in Python 3.11 using a variety of open-source packages. Model fitting was implemented primarily using JAX (***Bradbury et al., 2018***). Behavioral data were separated into AEPsych-driven and fallback trials on the one hand and validation trials on the other. The WPPM was fit exclusively to the AEPsych and fallback trials. To assess variability in model estimates, we performed 120 bootstrap resamplings of the AEPsych-driven trials, preserving the original ratio between Sobol’, adaptively sampled, and fallback trials in each resampled dataset. The WPPM was then re-fit to each of the 120 bootstrapped datasets.

To compute a 95% bootstrap confidence interval on the threshold contours, we first computed the summed Normalized Bures Similarity (NBS) score (***Muzellec and Cuturi, 2018***) between the predictions from the model fit to each bootstrapped dataset and those from the model fit to the original dataset, evaluated on a finely sampled grid of reference stimuli (-0.85 to 0.85 with 103 uniformly spaced steps). Higher scores indicate greater similarity to the predictions from the original dataset. We then sorted the model fits by their summed NBS scores and retained the top 114 (95% of 120) fits. The confidence interval bounds were defined by the union and intersection of the threshold contours, subsequently computed for any reference stimulus using this fixed set of retained model fits.

For the held-out validation trials, we computed the Euclidean distance between each reference and its paired comparison stimulus. For each of the 25 conditions, a Weibull psychometric function was fit to the binary color discrimination responses, with the guess rate fixed at 33.3% correct. Threshold was defined as the comparison stimulus corresponding to 66.7% correct. To estimate variability, we bootstrapped each condition 120 times and computed 95% confidence intervals for the threshold estimates.

To assess the agreement between the thresholds predicted by the WPPM and those estimated from the validation trials, we performed linear regression (constrained to pass through the origin) between the two sets of predictions using the original dataset, as well as for each of the 120 paired bootstrapped datasets. We then sorted the resulting slopes and correlation coefficients and computed 95% confidence intervals separately for each. Additionally, we computed the number of conditions for which the 95% bootstrap confidence intervals of the WPPM-predicted thresholds and the validation thresholds overlapped as an additional measure of agreement.

### Data and code availability

Data (https://osf.io/k27js/overview) and code (https://github.com/fh862/ellipsoids_eLife2025.git) are publicly available. All experiments, data collection, processing activities, and open sourcing were conducted at the University of Pennsylvania.

## Acknowledments

We thank Nicolas P. Cottaris for his assistance with calibration, our colleagues at the UPenn Vision Labs, Laurence Maloney, and Karl Gegenfurtner for helpful feedback.

## Author contribution

F.H.: Conceptualization; Formal analysis; Investigation; Methodology; Software; Validation; Visualization; Writing – original draft; Writing – review & editing.

R.B.: Investigation; Project administration.

J.C.: Methodology; Software; Writing – review & editing.

C.S.: Methodology; Software.

M.S.: Methodology; Software; Writing – review & editing.

P.G.: Conceptualization; Methodology; Resources; Software; Supervision; Validation; Writing – review & editing.

A.H.W.: Conceptualization; Methodology; Resources; Software; Supervision; Validation; Writing – original draft; Writing – review & editing.

D.H.B.: Conceptualization; Funding acquisition; Methodology; Project administration; Resources; Software; Supervision; Validation; Writing – original draft; Writing – review & editing.

## Appendix 1 Transformation between the DKL, RGB, and model spaces

This section summarizes the colorimetric transformations between the RGB space of our monitor, the 2D model space, and standard color spaces.

The DKL space provides a representation of the isoluminant plane with the adapting point at the origin (***Derrington et al., 1984; Brainard, 1996***). We defined our DKL space with respect to the CIE physiologically-relevant 2-degree cone fundamentals and corresponding photopic luminosity function. We began in DKL space, with adapting point defined by the cone excitations elicited by the displayed background so that the space’s isoluminant plane included the background. In this plane, we then densely sampled chromatic directions spanning 360° around the origin. For each direction we marched outward from the origin to find the edge of the monitor’s gamut in that direction (details explained in Appendix 1.1). Repeating across directions, we obtained a set of gamut boundary points that define a quadrilateral in the isoluminant plane. We then identified the four vertices and recorded their coordinates in both DKL and RGB spaces (***Table S1***). Using these vertices, we derived a projective transformation matrix (homography) that maps coordinates from the DKL space to the model space (see Appendix 1.2 and ***Table S2***). While the homography provides a general solution applicable to any quadrilateral, our derived matrix revealed that an affine transformation provided an accurate approximation. We used this affine transformation and its inverse to convert back and forth between linear RGB values and the model space (see Appendix 1.3 and ***Table S2***).

### Appendix 1.1: Search for the boundary points within the monitor’s gamut

We selected 1,000 angles *θ* that span the isoluminant plane uniformly in the DKL representation. For each angle, we defined a chromatic direction vector in DKL space as **d**_DKL_ = [cos(*θ*), sin(*θ*), 0]^⊤^, where the first two elements correspond to the L–M and S axes of the DKL space, and the third element is set to zero to constrain the direction to the isoluminant plane (i.e., no change in luminance). For each direction, we then determined the farthest point along that vector that remained within the monitor’s gamut in linear RGB space. Specifically, we first converted **d**_DKL_ to LMS cone excitations, denoted **e**_LMS_, as follows:

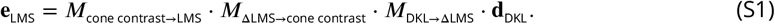

Notably, because the actual LMS cone excitations include contributions from ambient light, denoted as **e**_LMS, ambient_, we subtracted it to isolate the portion due to the RGB stimulus, denoted as **e**_LMS, stimulus_, that is,

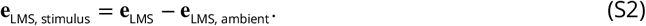

Next, we converted this isolated stimulus response into RGB space:

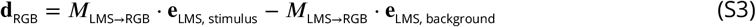

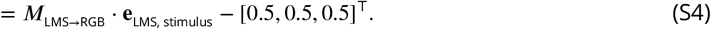

Finally, we marched outward along the direction **d**_RGB_ until the RGB values reached the edge of the RGB cube. The values at this boundary were recorded as a limiting point along that chromatic direction. Repeating this procedure across all sampled directions yielded the full boundary of the isoluminant plane constrained by the monitor’s gamut. From this boundary set, we then identified the four corner vertices (***Table S1***).

**Table S1.** Corner vertices in the DKL, LMS, RGB, and model spaces.

| Corner | DKL<sub>L-M</sub> | DKL<sub>S</sub> | DKL<sub>Lum</sub> | L | M | S | R | G | B | W<sub>dim1</sub> | W<sub>dim2</sub> |
| --- | --- | --- | --- | --- | --- | --- | --- | --- | --- | --- | --- |
| 1 | -0.123 | -0.812 | 0 | 0.145 | 0.147 | 0.016 | 0.000 | 0.733 | 0.000 | -1 | -1 |
| 2 | 0.175 | -0.830 | 0 | 0.164 | 0.111 | 0.014 | 1.000 | 0.407 | 0.000 | 1 | -1 |
| 3 | -0.175 | 0.830 | 0 | 0.142 | 0.154 | 0.152 | 0.000 | 0.593 | 1.000 | -1 | 1 |
| 4 | 0.123 | 0.812 | 0 | 0.160 | 0.117 | 0.150 | 1.000 | 0.267 | 1.000 | 1 | 1 |

### Appendix 1.2: An affine transformation matrix that maps DKL to model space

These vertices, denoted as **v**, were then used to derive a projective transformation matrix *M*_DKL→W_ such that for each vertex pair, we have:

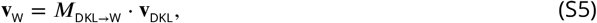

where **v**_W_ = [*v*_W, dim1_, *v*_W, dim2_, 1]^⊤^ is the homogeneous coordinate of a vertex in model space, and **v**_DKL_ = [*v*_DKL, *L*-*M*_, *v*_DKL, *S*_, 0]^⊤^ is the corresponding homogeneous coordinate in DKL space. By plugging in the vertices, we solved the matrix *M*_DKL→W_ as the following,

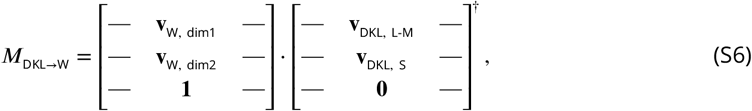

where † denotes the pseudoinverse. Note that the last row of *M*_DKL→W_ is [0, 0, 1], indicating that the transformation is affine (***Table S2***). Although this affine formulation would be sufficient, we initially computed the full projective transformation matrix for generality, since it was uncertain that the DKL vertices would form a parallelogram. In this particular case, both methods yielded equivalent results.

**Table S2.** Transformation matrices between DKL, RGB and model spaces.

| $M_{\text{DKL} \rightarrow \text{W}}$ | $M_{\text{RGB} \rightarrow \text{W}}$ |
| --- | --- |
| $\begin{bmatrix} 6.724 & 0.213 & 0.000 \\ 0.076 & 1.221 & 0.000 \\ 0.000 & 0.000 & 1.000 \end{bmatrix}$ | $\begin{bmatrix} 1.556 & -1.364 & -0.192 \\ -0.444 & -1.364 & 1.808 \\ 0.444 & 1.364 & 0.192 \end{bmatrix}$ |

#### Appendix 1.3: An affine transformation matrix that maps RGB to model space

Given that the transformations from DKL to LMS, LMS to RGB, and DKL to model space are all affine, by the composition property of affine transformations, it follows that the transformation from RGB to model space must also be affine.

To compute the affine transformation matrix, we used corresponding corner vertices in RGB and model spaces. Specifically, we solved for the matrix *M*_RGB→W_ as:

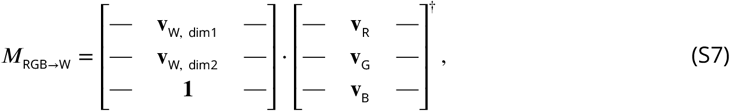

where † denotes the pseudoinverse and we appended a row of ones to the Wishart coordinates to express them in homogeneous form.

### Appendix 1.4: Affine invariance of Mahalanobis distance

We performed trial placement, model fitting, and data presentation using the model space (bounded between –1 and 1). An important feature of the WPPM is that it is equivariant with respect to affine transformations of the color space used to represent the stimuli. That is, if we transform reference and comparison stimuli to a new color space using an affine transformation, and transform the covariance field by the same affine transformation, then the observer model yields a prediction of performance that is unchanged by the transformation. This is because the Mahalanobis distance is itself unchanged by the transformation, as we show below. This is an attractive property because it avoids assigning special status to the particular color space used to represent the stimuli and covariance field.

Let Σ be the covariance matrix. The squared Mahalanobis distance between two points **x**_0_ and **x**_1_ is defined as:

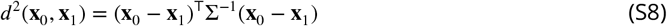

Now consider a linear transformation 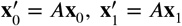. The corresponding transformation of the covariance matrix is Σ^′^ = *A*Σ*A*^⊤^. Then the squared Mahalanobis distance in the transformed space becomes:

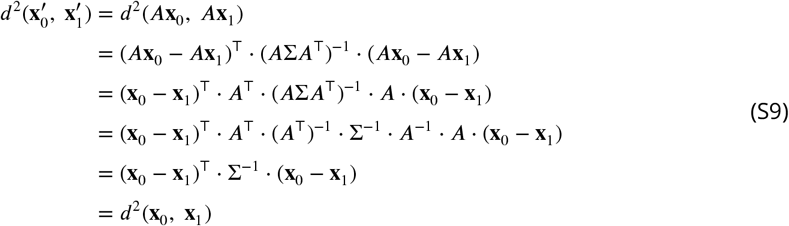

Thus, the Mahalanobis distance is invariant under linear transformations of the data when the covariance matrix is transformed accordingly. Since distance is also invariant to translations (i.e., independent of the choice of origin), this further implies that the Mahalanobis distance is invariant under general affine transformations.

## Appendix 2 Adaptive sampling and WPPM estimates

### Appendix 2.1: AEPsych-driven trials and WPPM-predicted thresholds

**Figure S1.**
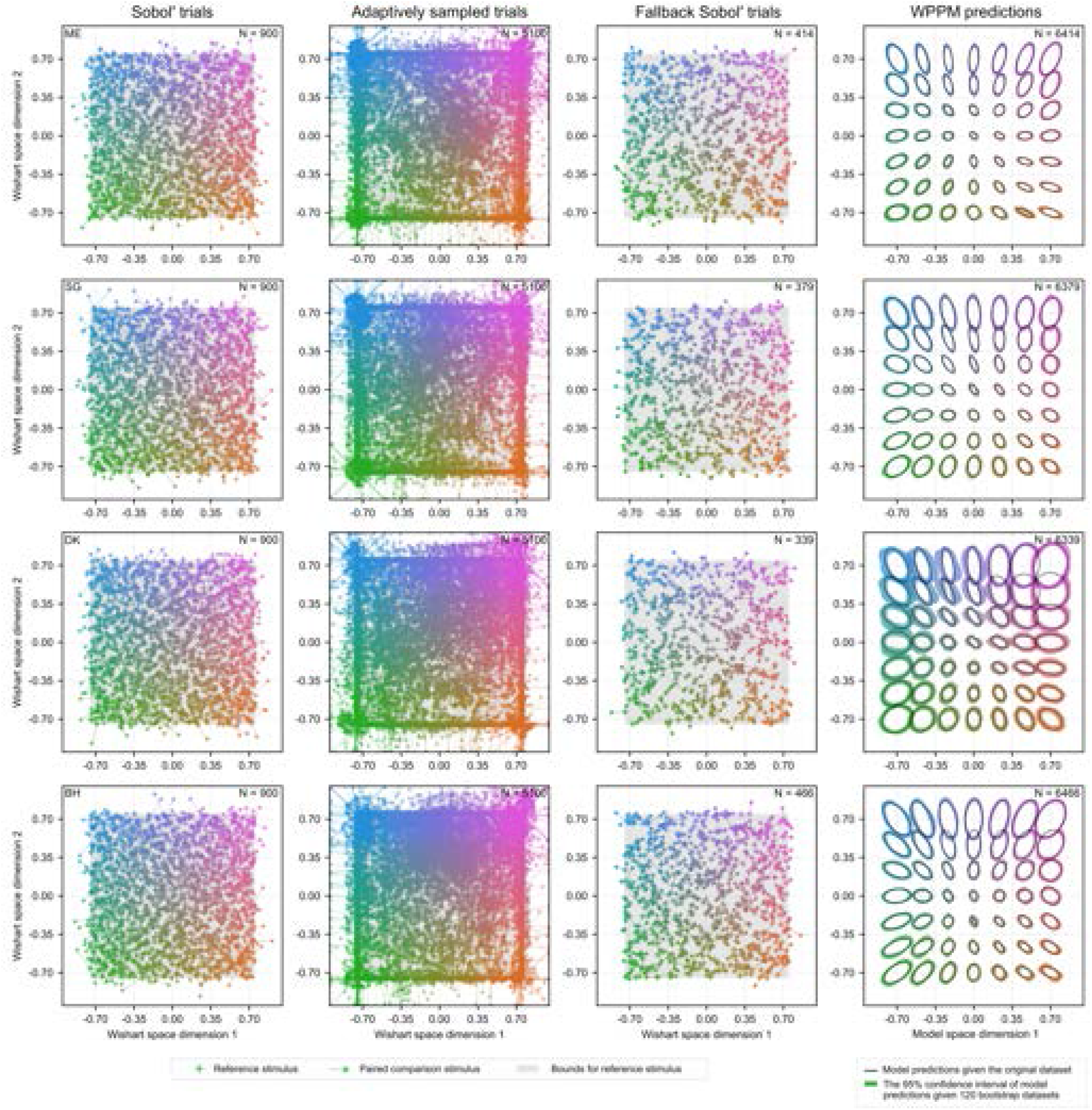
AEPsych-driven trials (900 Sobol’-sampled and 5,100 adaptively sampled), fallback trials, and WPPM predictions for all participants. Each row represents data from one participant.

**Figure S1.**
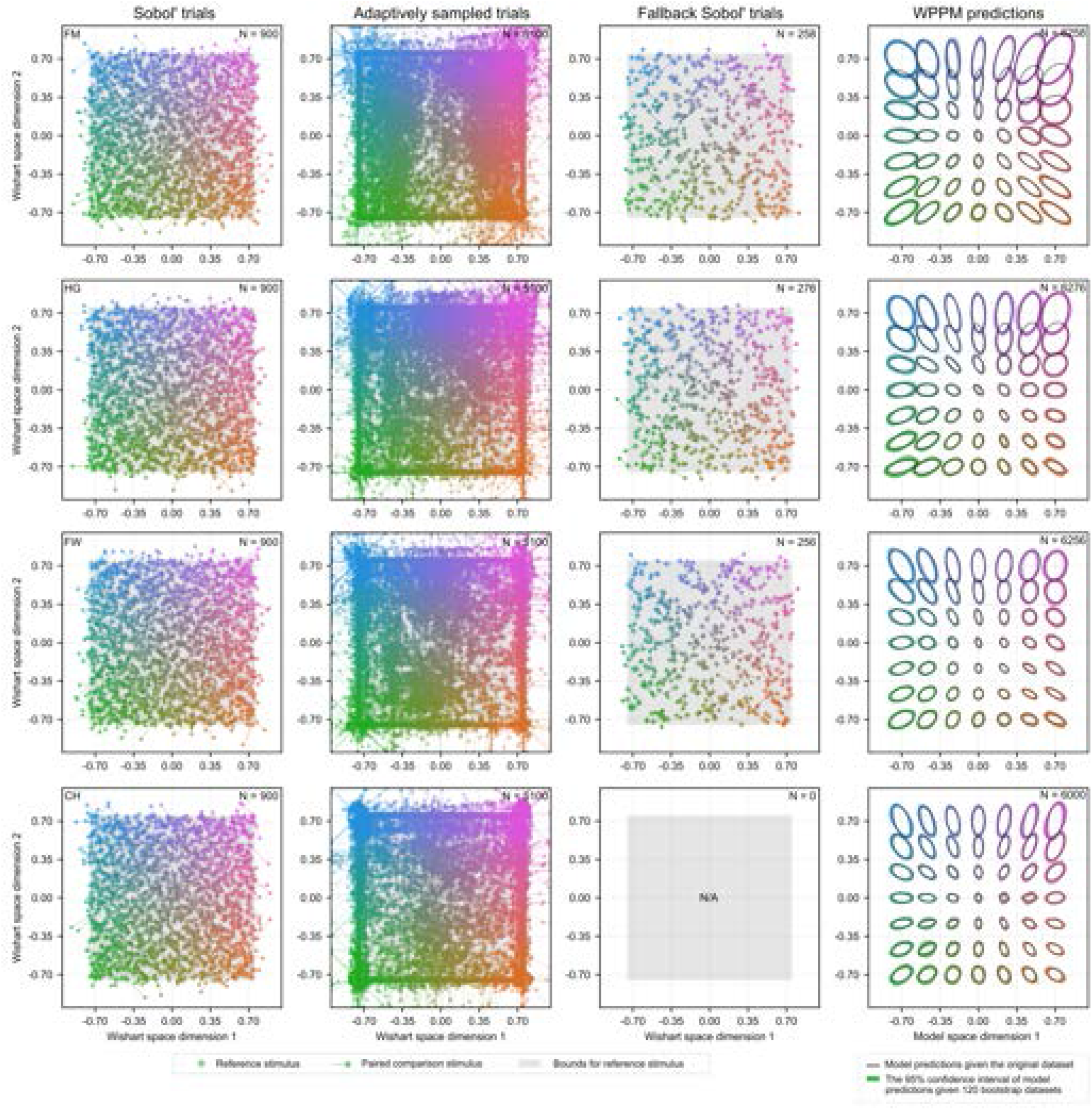
AEPsych-driven trials (900 Sobol’-sampled and 5,100 adaptively sampled), fallback trials, and WPPM predictions for all participants (continued). Each row represents data from one participant. Note that for participant CH, no pre-generated Sobol’ trials were used, as the fallback strategy was implemented later in the study to maintain experimental continuity and reduce delays between trials.

**Figure S2.**
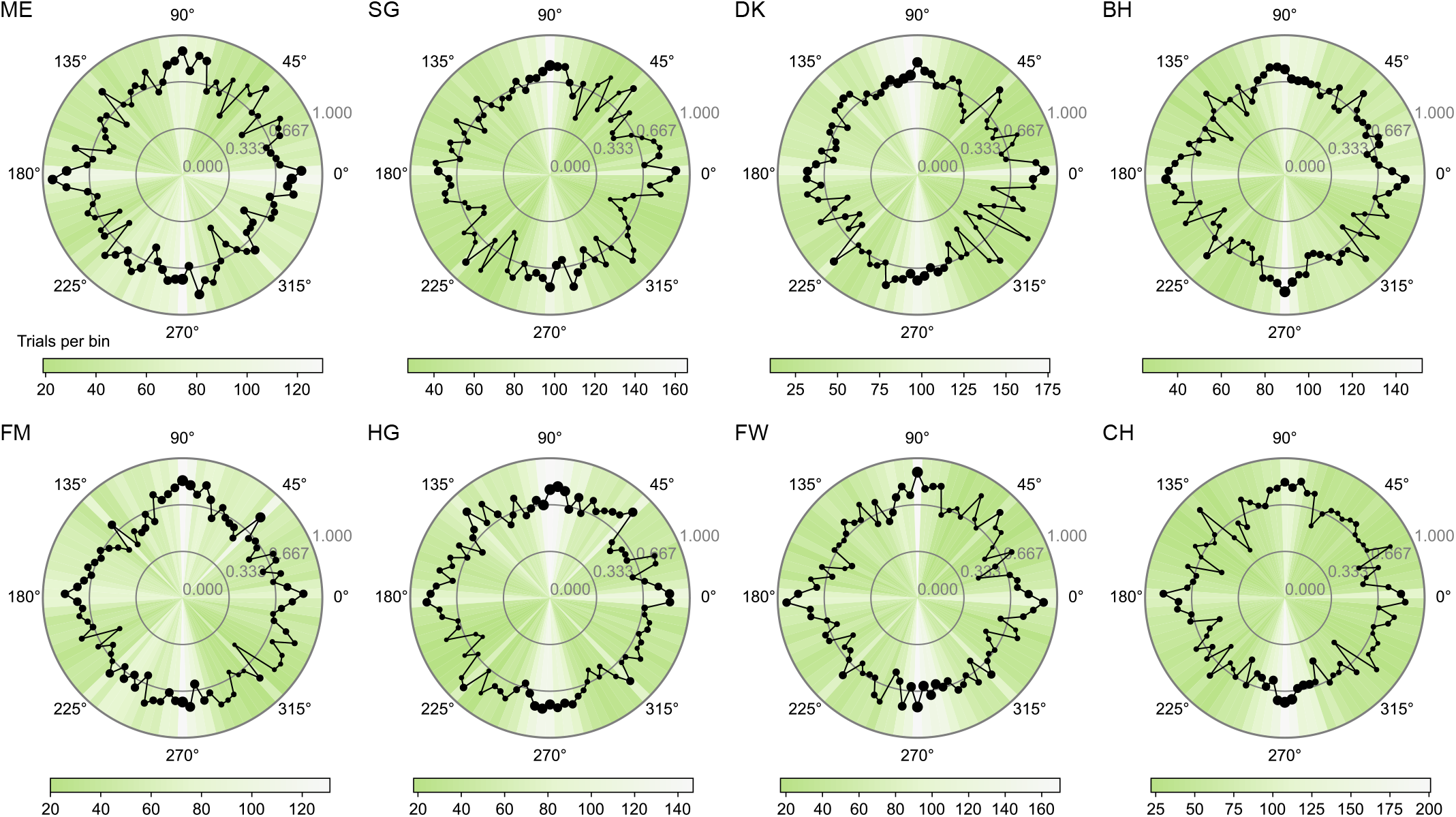
Percent correct as a function of the angular difference between reference and comparison stimuli for all participants. The number of trials within each bin (bin width = 4°) is encoded by both marker size and color, with larger markers and brighter colors indicating a greater number of trials.

### Appendix 2.2: Efficacy of adaptive sampling

After the initial 900 Sobol’-sampled trials, the subsequent 5,100 trials were adaptively placed using AEPsych (***Owen et al., 2021; Letham et al., 2022***) to target the 66.7% performance threshold. To evaluate the efficiency of this adaptive sampling procedure, we binned these adaptive trials based on the angular difference between the reference and comparison stimuli and computed the proportion of correct responses within each bin. If adaptive sampling is effective, we expect performance to cluster around 66.7% correct. Consistent with this, the observed percent correct remained close to 66.7% correct across bins (***Figure S2***), suggesting that AEPsych provided informative trial placement.

**Figure S3.**
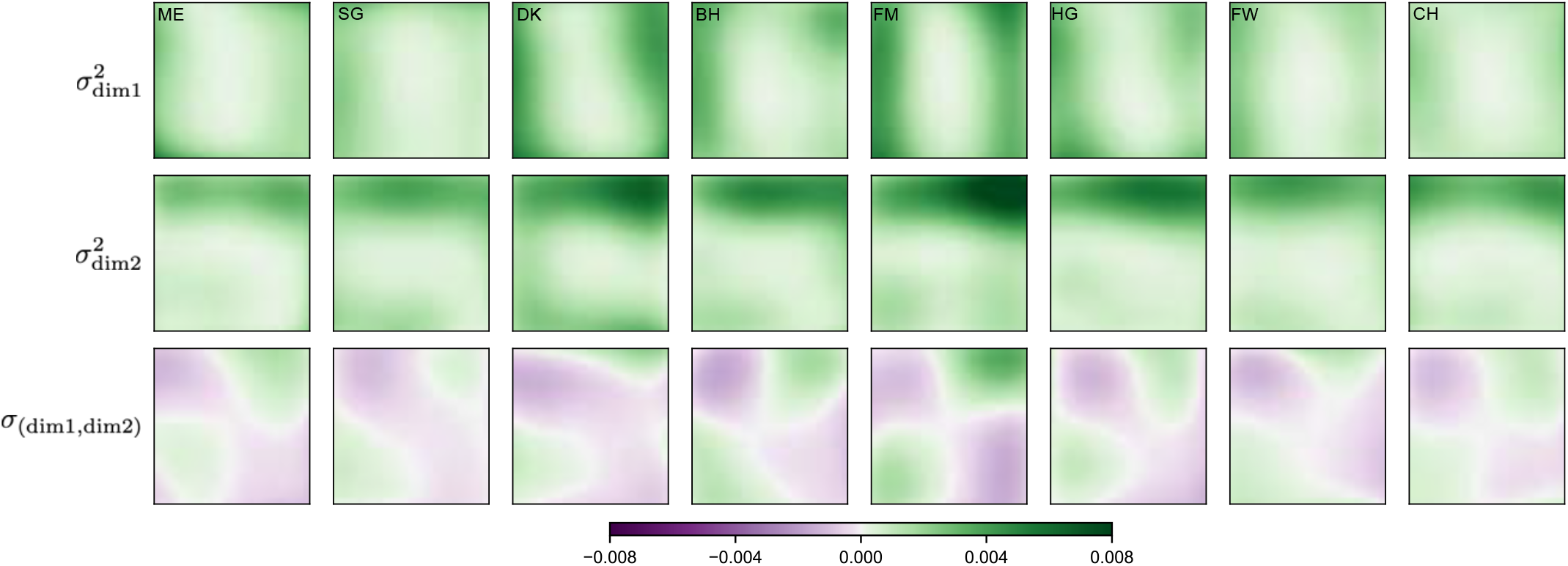
Covariance matrix fields obtained from the best-fitting WPPM for all participants. Rows show (top to bottom): noise variance along the first model dimension, noise variance along the second model dimension, and the covariance between the two dimensions.

### Appendix 2.3: The covariance matrix field

The WPPM specifies a covariance matrix at each reference stimulus across the isoluminant plane. Each matrix characterizes internal noise in terms of the variance along the two model dimensions 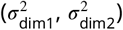 and their covariance (*σ*_dim1,dim2_). For each participant, the best-fitting model produces a covariance matrix field that exhibits qualitatively similar structure across individuals (***Figure S3***).

### Appendix 2.4: The best-fitting weight matrices

The covariance matrix field is determined by the weights applied to the Chebyshev basis functions. Across participants, the absolute magnitude of the best-fitting weights decreases with increasing basis order, indicating that higher-order components contribute less to the covariance field. At the highest order we included, the weights are close to zero. This indicates that given the prior hyperparameters, adding more basis functions would be unlikely to change the estimates we obtained.

**Figure S4.**
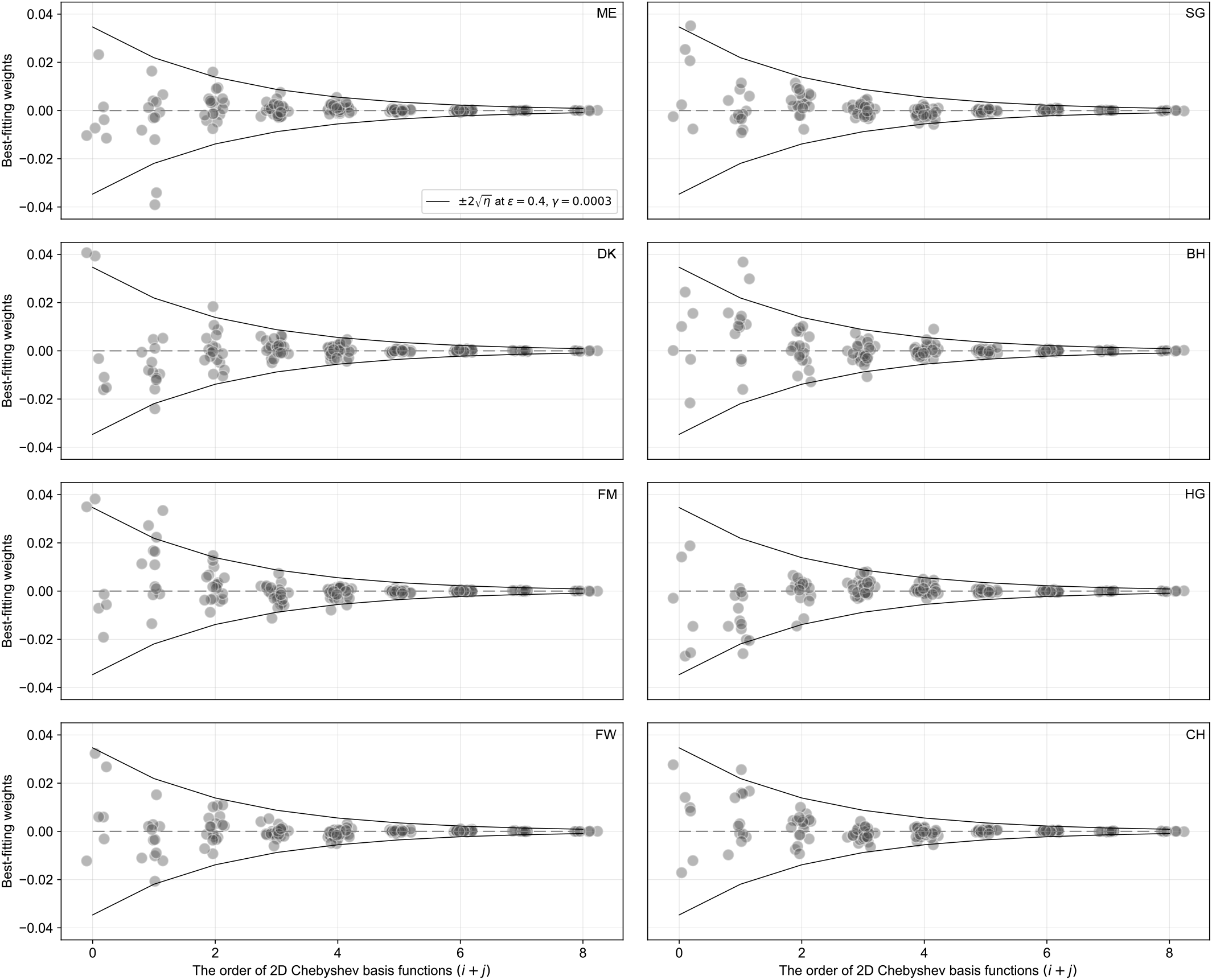
The best-fitting weights for all participants. The symmetric solid curves represent the prior imposed on the model. The prior is implemented by specifying the variance of the weights, *η*, as a function of the polynomial order of the Chebyshev basis functions and two hyperparameters *ϵ* and *γ* (***Equation 16***). The solid curves indicate 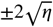 for our hyperparameter choice, corresponding to two standard deviations of the prior distribution.

## Appendix 3 Real-time trial scheduling via dual-computer coordination

**Figure S5.**
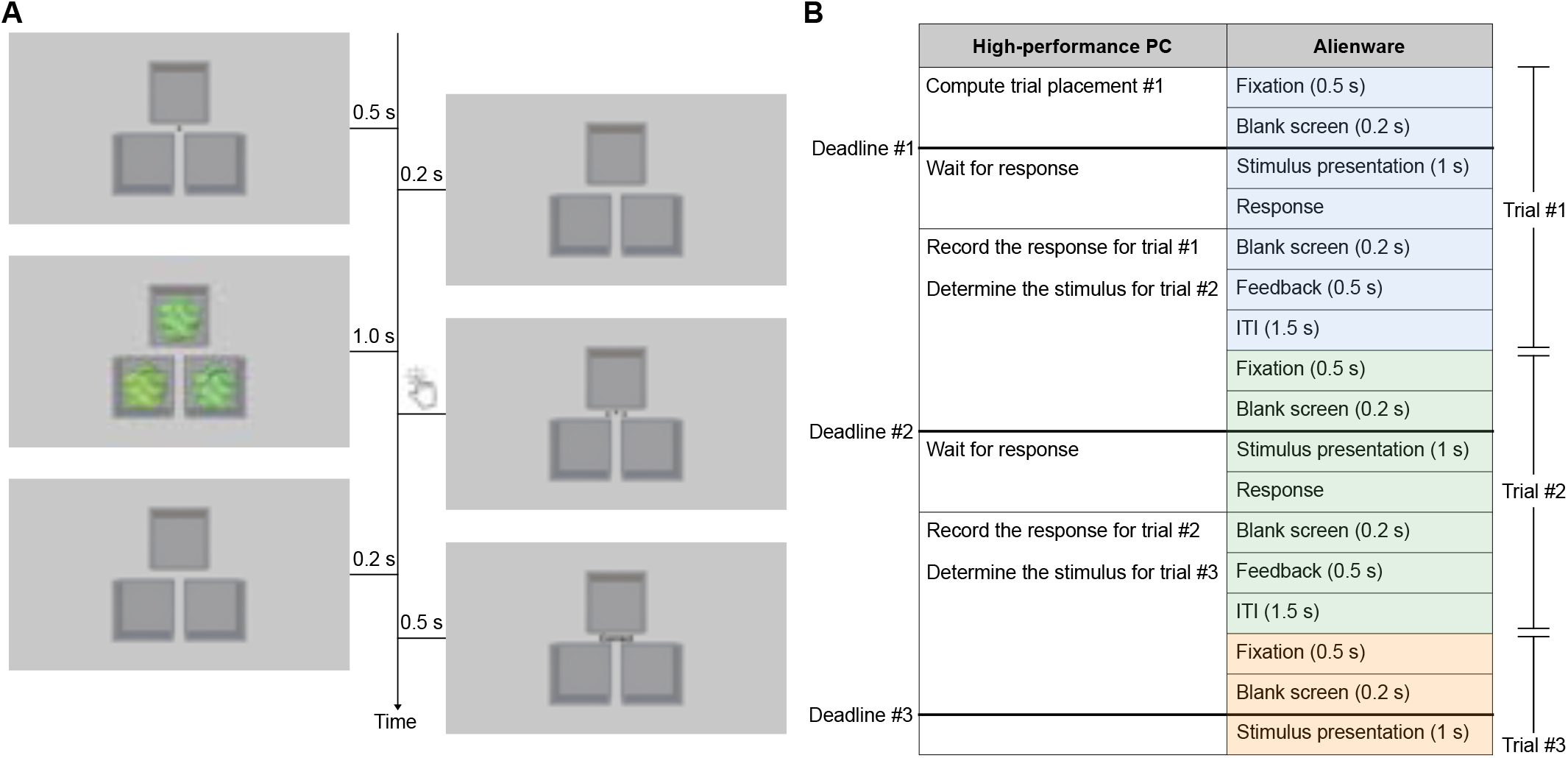
Task timing and real-time trial scheduling. (A) Trial sequence: a 0.5 s fixation cross was followed by a 0.2 s blank interval, then a 1 s presentation of three blobby stimuli. Participants responded at their own pace to identify the odd-one-out, after which a 0.2 s blank screen and a 0.5 s feedback were shown. The inter-trial interval (ITI) was 1.5 s. (B) A schematic representation of the trial timing and computational responsibilities of the two computers.

For each participant, we ran 6,000 AEPsych-driven trials interleaved with an additional 6,000 validation trials. Although our initial plan was to present these 12,000 trials in a predetermined randomized sequence, we quickly realized this approach was impractical. Under such a design, AEPsych would have only the inter-trial interval (ITI) to compute the next trial placement—a window that is difficult to optimize. A long ITI risks participant fatigue or loss of attention, while a short ITI does not provide AEPsych with enough time to complete its computations. To achieve both a smooth experimental flow and adequate computation time for AEPsych, we implemented a fallback trial strategy using a dual-computer setup.

In this setup, stimulus presentation was handled by an Alienware computer, while adaptive trial placement using AEPsych ran on a separate high-performance PC. The two systems communicated via a shared network disk using a custom protocol based on text files that both computers could read and write. This decoupled design provided modular separation between code specialized for stimulus presentation and the sequence of events on each trial and code specialized for trial placement, and should allow our trial placement code to be more easily ported to different stimulus display systems.

With the dual-computer design, AEPsych had at least 2.9 s to compute the next trial after the participant’s response (***Figure S5***). This window spanned both the post-stimulus period of the current trial (0.2 s blank, 0.5 s feedback, 1.5 s ITI) and the pre-stimulus period of the upcoming trial (0.5 s fixation and 0.2 s blank). Importantly, this computation window began only after the participant responded—since AEPsych requires the participant’s response to update its model—and ended just before the stimulus presentation of the next trial, when AEPsych must deliver the RGB values for the upcoming stimuli.

The fallback trial strategy ensured continuous stimulus presentation. If AEPsych failed to return a new trial within the 2.9 s window, we defaulted to presenting the next available MOCS trial from the pre-determined randomized sequence. In such cases, AEPsych’s computation continued in a different thread and attempted to meet the following decision deadline, which is approximately 7 s after the previous one, including an additional 1 s stimulus presentation and an estimated 0.2 s response time. If AEPsych again missed this deadline, the next opportunity came at around 11.1 s. This staggered scheduling ensured that trials continued smoothly while allowing AEPsych sufficient time to compute adaptive placements when possible.

A potential drawback of the fallback trial strategy is that it could disrupt the intended interleaving of adaptive and validation trials, potentially introducing differential learning effects. To mitigate this, we capped how far MOCS trials could advance relative to AEPsych trials. This cap was set at four trials. If this limit was reached and no AEPsych trial was ready, we inserted pre-generated Sobol’ trials instead. These Sobol’ trials were created in advance using participant- and session-specific random seeds and were separate from those selected by AEPsych.

## Appendix 4 Comparison between WPPM and validation thresholds

### Appendix 4.1: Validation data for all participants

**Figure S6.**
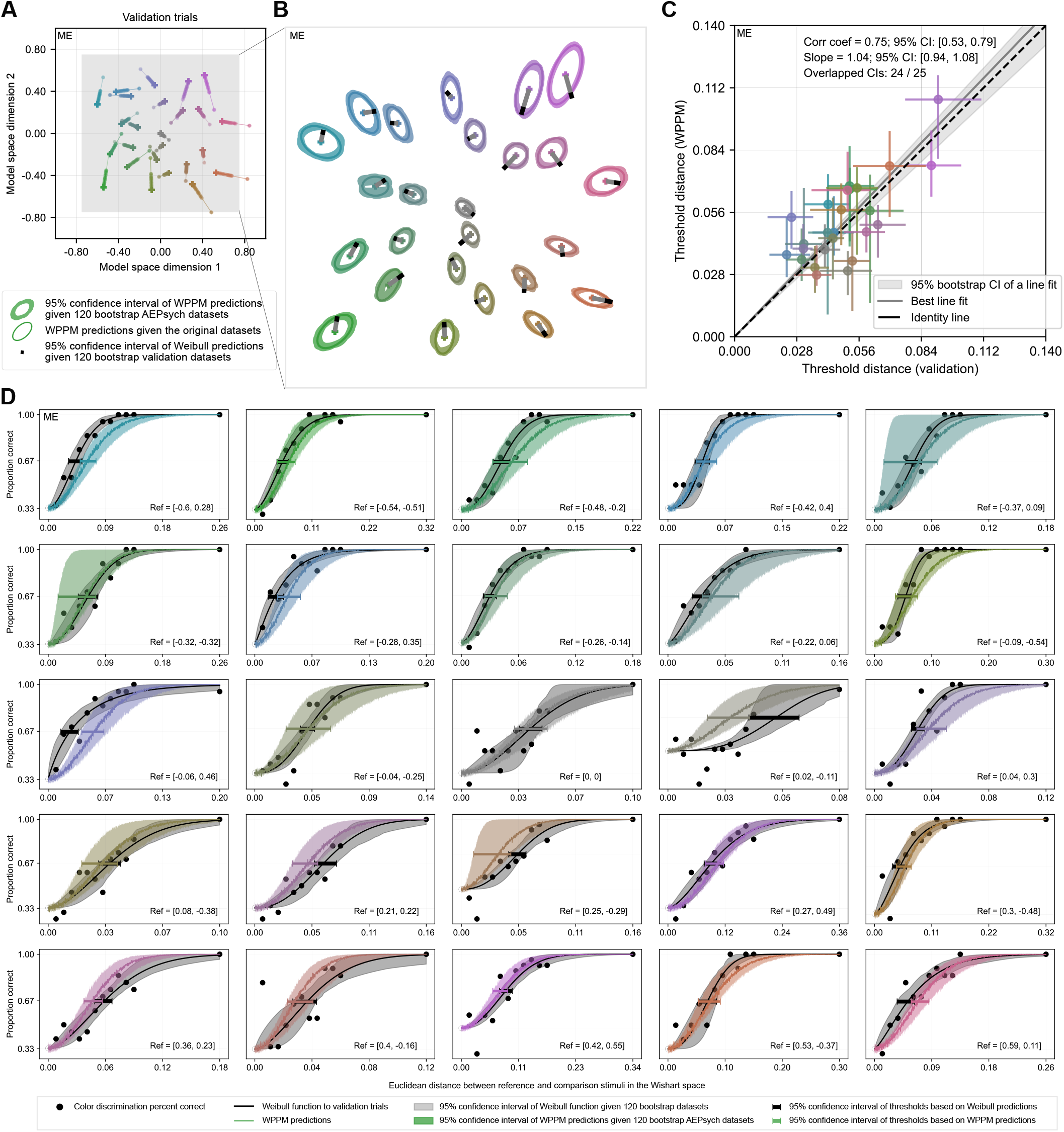
Validation for participant ME. Same format as ***Figure 2***D-G in the main text.

**Figure S7.**
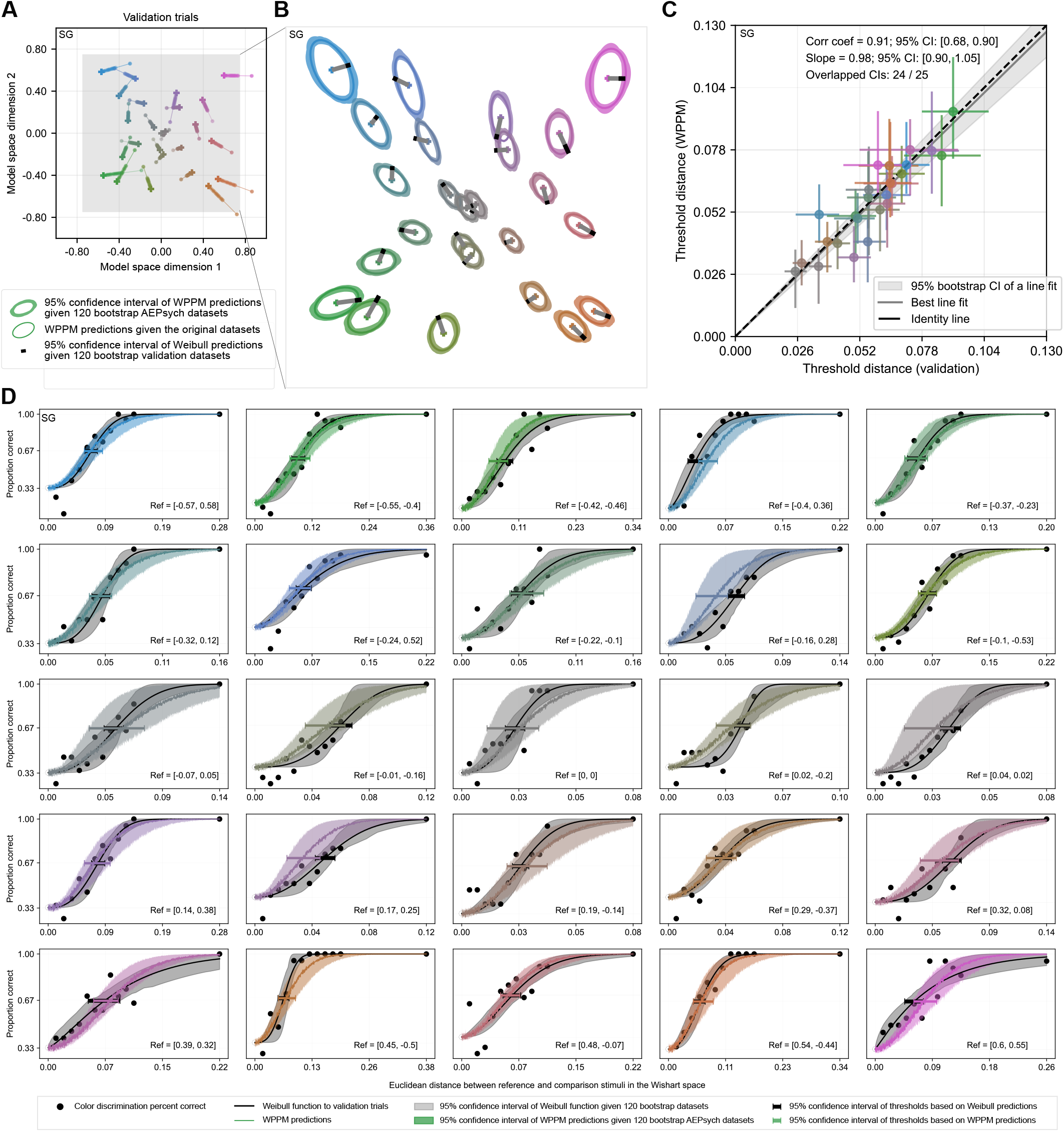
Validation for participant SG.

**Figure S8.**
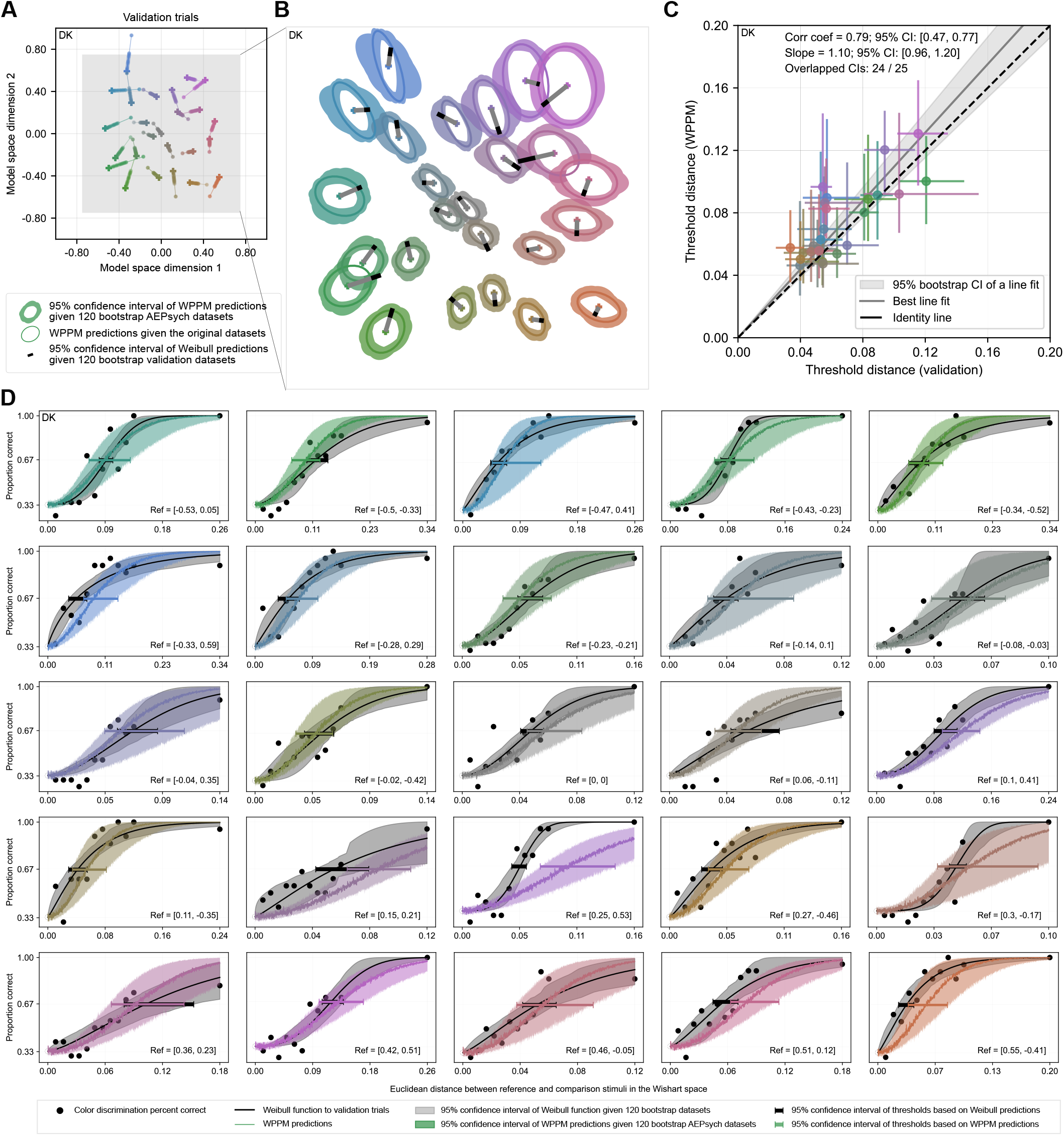
Validation for participant DK.

**Figure S9.**
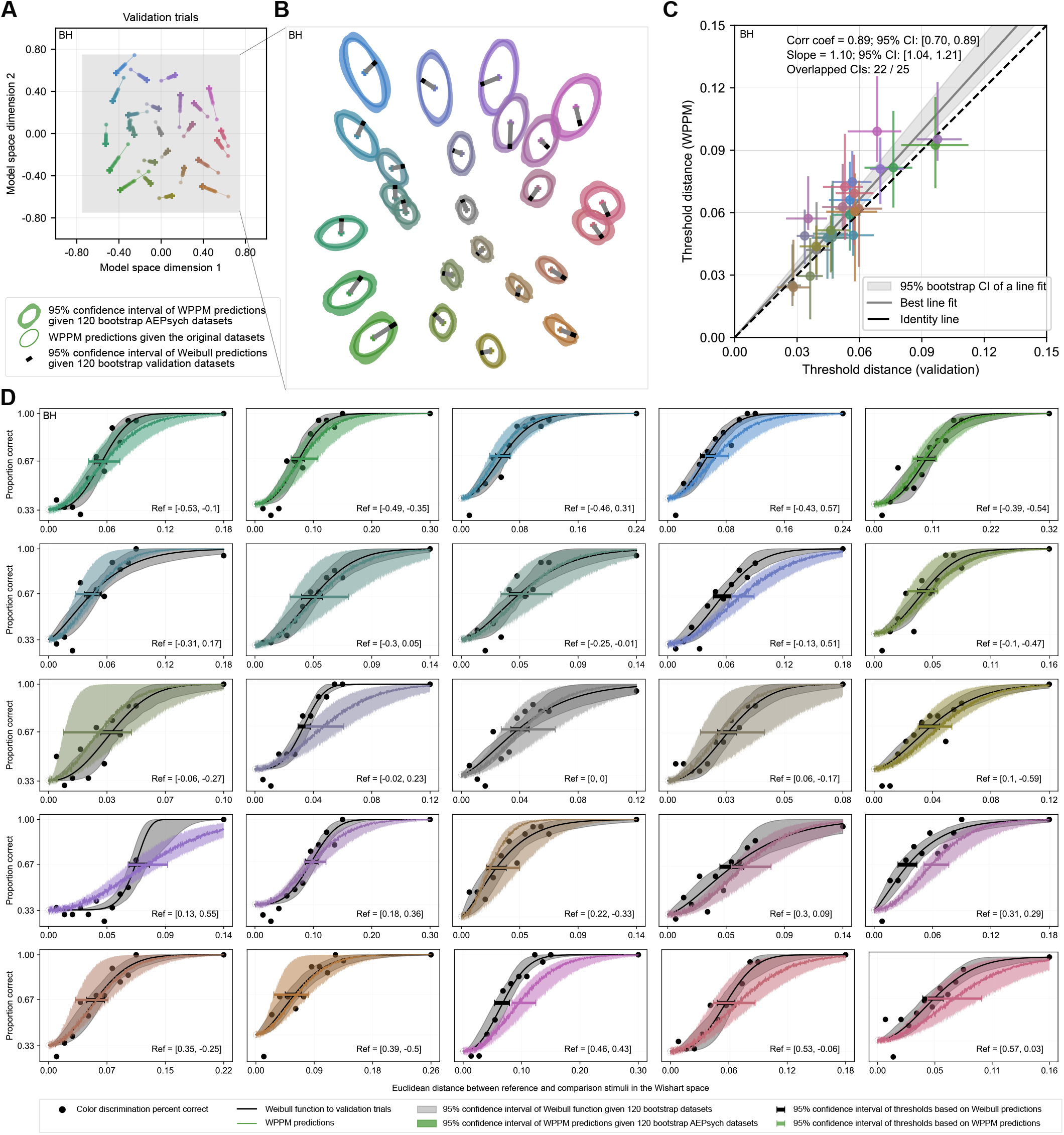
Validation for participant BH.

**Figure S10.**
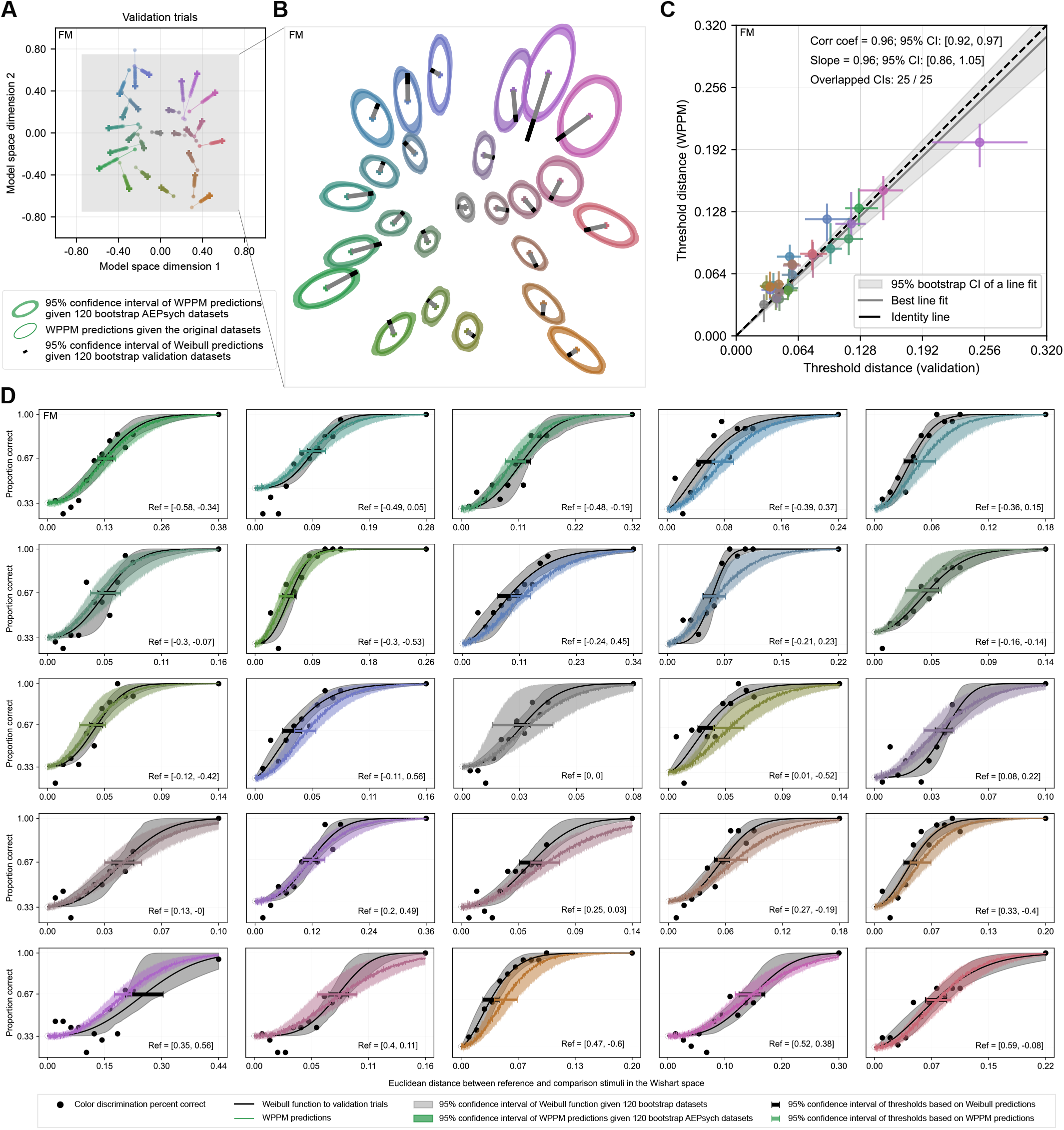
Validation for participant FM.

**Figure S11.**
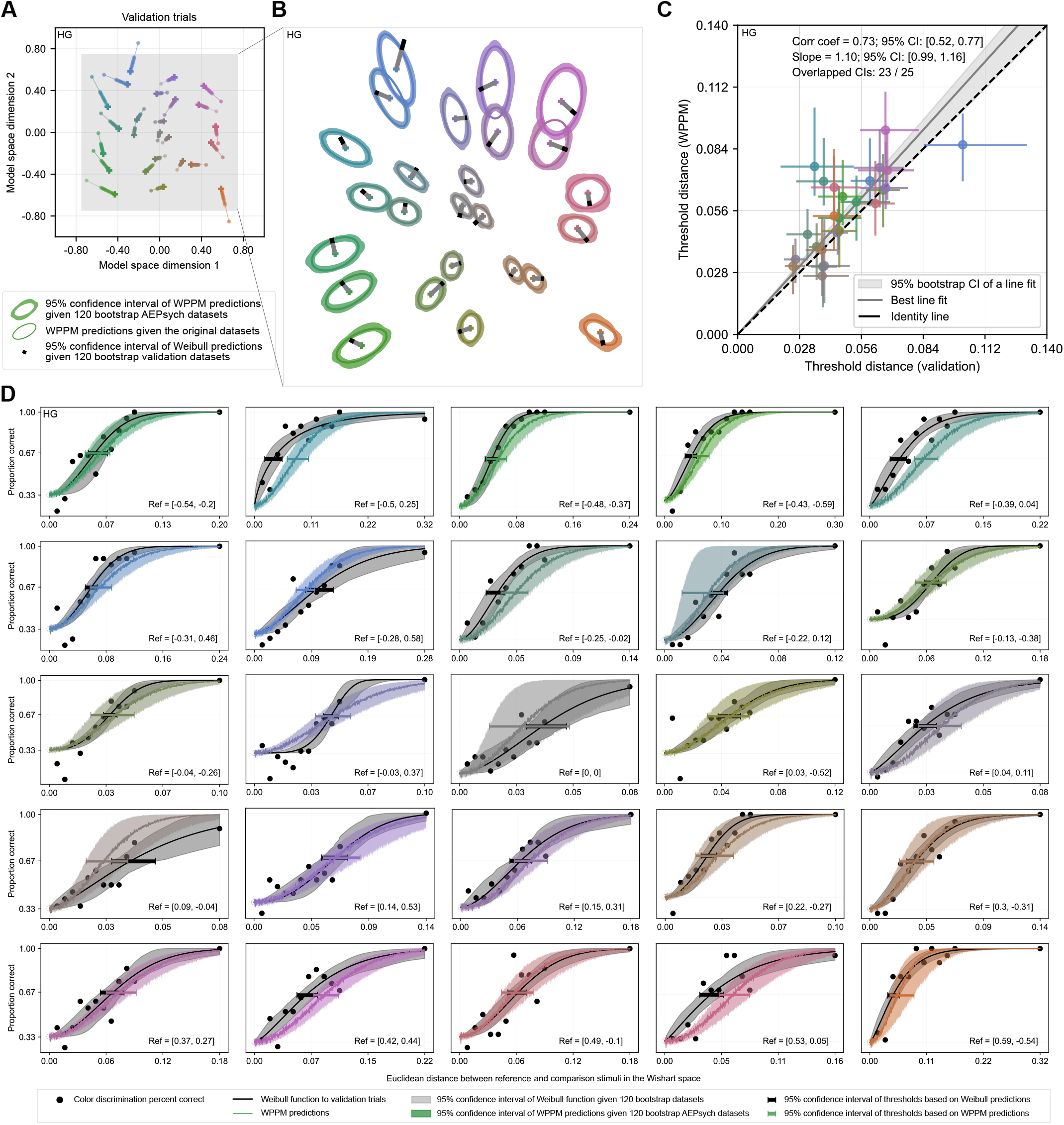
Validation for participant HG.

**Figure S12.**
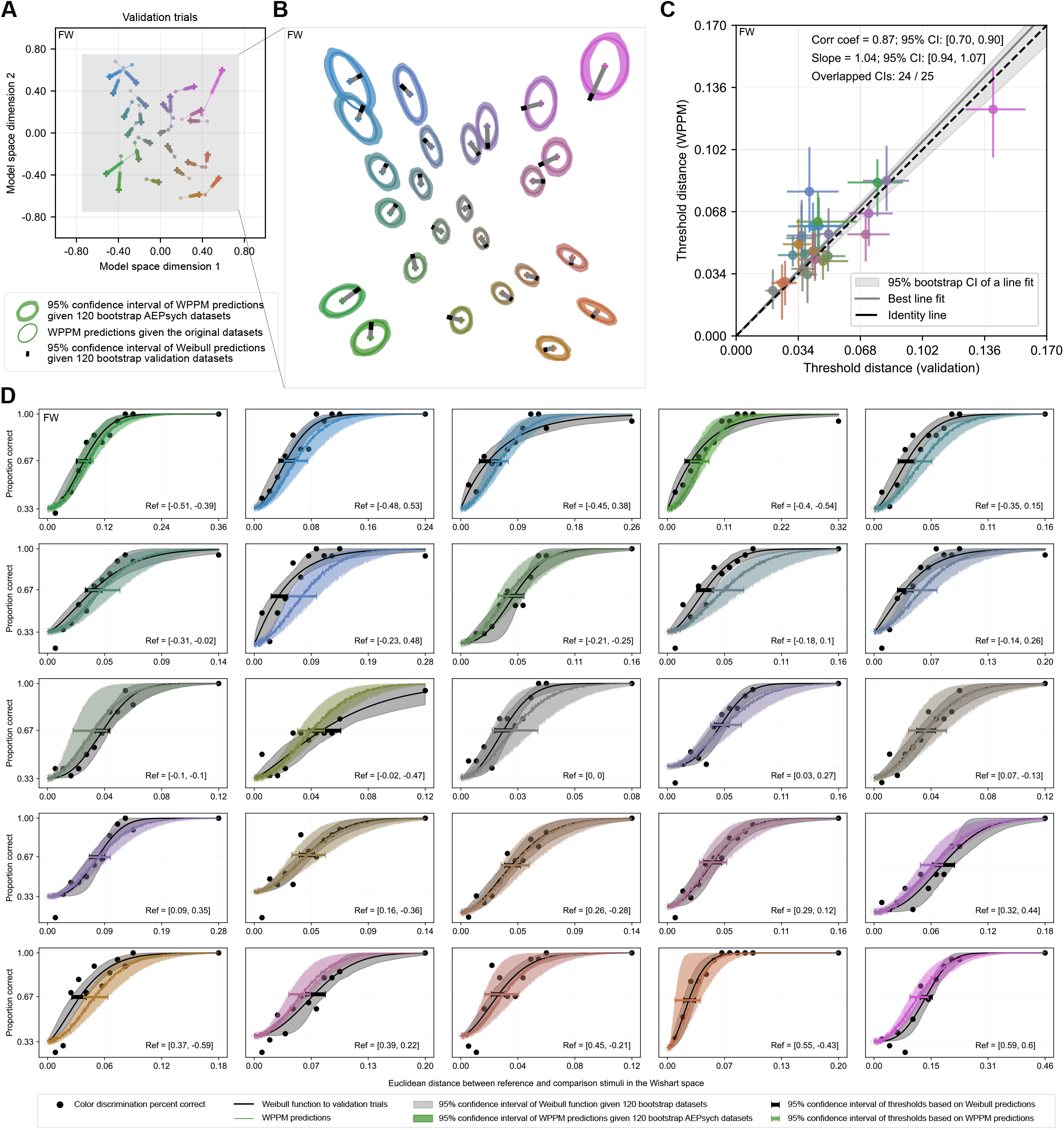
Validation for participant FW.

**Figure S13.**
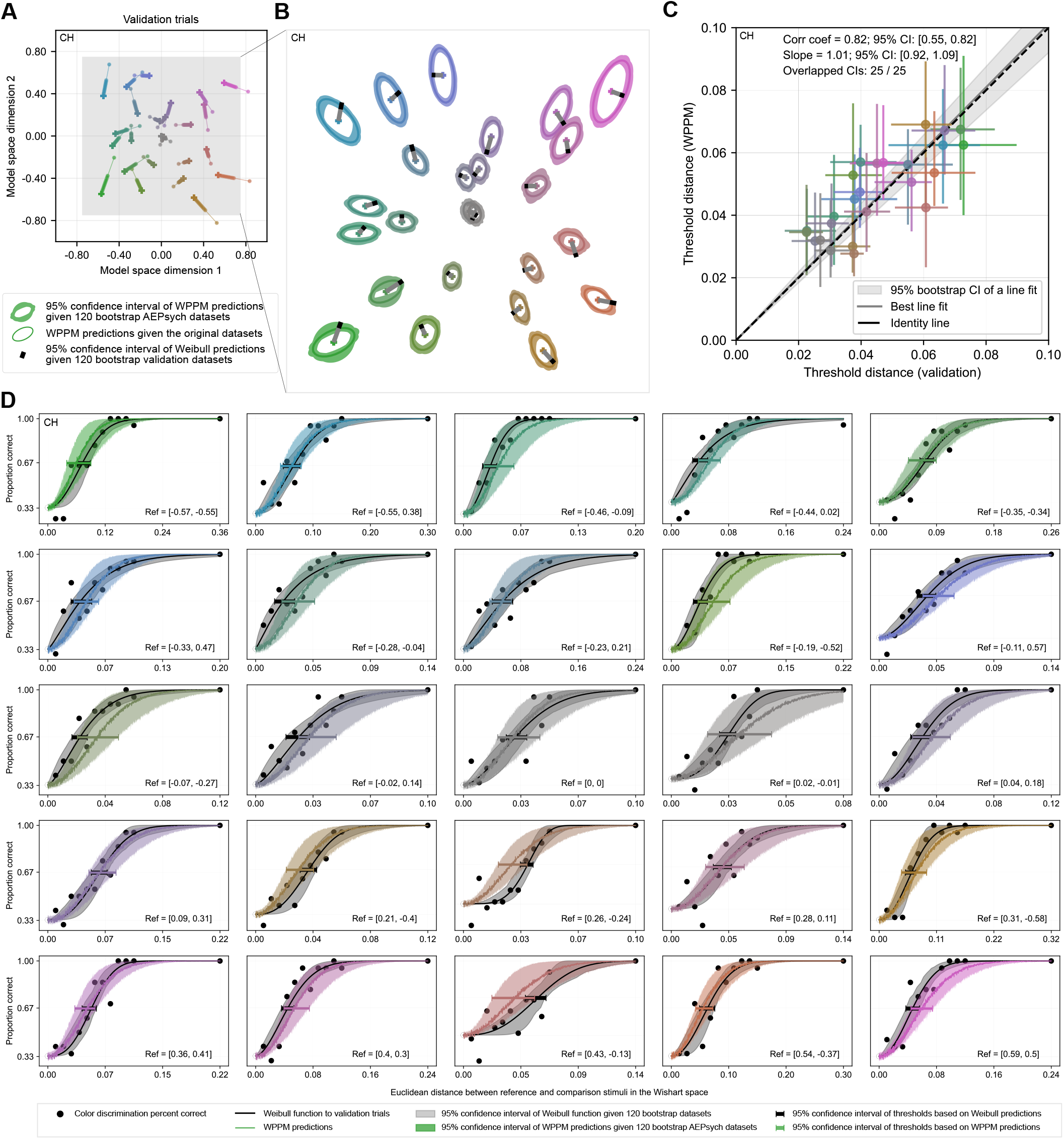
Validation for participant CH.

### Appendix 4.2: Analysis of differences between WPPM and validation thresholds

**Figure S14.**
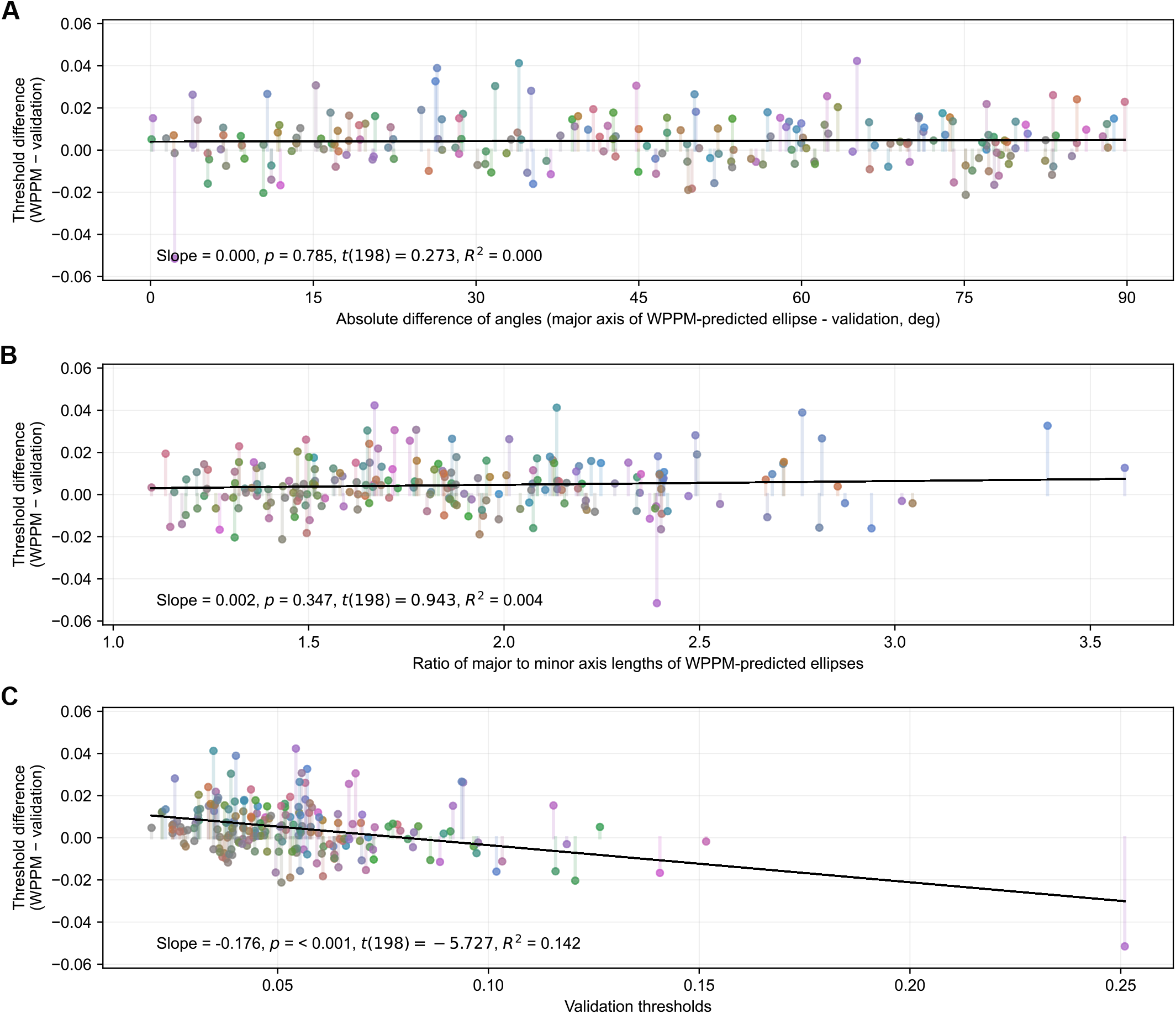
Threshold residuals. Data are pooled across all validation conditions and all participants (*N* = 8). For all panels, color codes for the surface color of the reference stimulus, and the y-axis limits are set to ± the mean of the validation thresholds. (A) Residuals as a function of the absolute angular difference between the major axis of the elliptical threshold contours read out from the WPPM fits and the chromatic direction of the validation condition. (B) Residuals as a function of the aspect ratio (major/minor axis) of the WPPM threshold contours. (C) Residuals as a function of thresholds estimated from validation trials.

We assessed whether the residuals (the differences between the WPPM and validation thresholds) exhibited systematic patterns. We found no significant correlation between the residuals and the absolute angular difference between the chromatic direction of the validation condition and the major axis of the elliptical threshold contours read out from the WPPM fits (***Figure S14***A), nor with the aspect ratio of the contours (***Figure S14***B). Thus, there is no evidence that the residuals vary systematically with the orientation or shape of the contours read out from the WPPM fits (see statistical summary in ***Table S3***).

In contrast, we found a significant negative correlation between the residuals and the magnitude of the validation thresholds (***Figure S14***C; slope = −0.176, *t*(198) = −5.727, *p* < 0.001, *R*^2^ = 0.142), indicating that the WPPM tends to slightly overestimate thresholds when they are small and underestimate them when they are large. However, the magnitude of this bias is small relative to the range of observed validation thresholds.

**Table S3.** Linear regression results assessing the relationship between WPPM–validation threshold residuals and three predictors:(1) the absolute angular difference between the chromatic direction of the validation condition and the major axis of the contours read out from the WPPM fits, (2) the aspect ratio of the contours, and (3) the magnitude of the validation threshold. This analysis was done on human data.

| Predictor | Term | Coef | Std Err | *t* | *p* | [0.025, 0.975] CI | *R*<sup>2</sup> |
| --- | --- | --- | --- | --- | --- | --- | --- |
| Absolute difference of angles | intercept | 0.004 | 0.002 | 2.254 | 0.025 | [0.000, 0.007] | 0.000 |
|  | slope | 0.000 | 0.000 | 0.273 | 0.785 | [-0.000, 0.000] |  |
| Aspect ratio | intercept | 0.001 | 0.004 | 0.319 | 0.750 | [-0.006, 0.008] | 0.004 |
|  | slope | 0.002 | 0.002 | 0.943 | 0.347 | [-0.002, 0.005] |  |
| Validation thresholds | intercept | 0.014 | 0.002 | 7.511 | 0.000 | [0.010, 0.018] | 0.142 |
|  | slope | -0.176 | 0.031 | -5.727 | 0.000 | [-0.237, -0.116] |  |

### Appendix 4.3: Analysis of percent-correct performance for catch trials

For each validation condition, we used the method of constant stimuli to sample 12 comparison levels: 11 were evenly spaced, and one was selected to serve as an easily discriminable catch trial. Participants completed 500 catch trials (1/12 of 6,000 validation trials). These catch trials were included to assess participants’ attentiveness and establish a criterion for potential data exclusion. As shown in ***Table S4***, participants except for DK performed near ceiling on these trials, indicating high task engagement throughout the experiment. Although DK’s performance was somewhat lower, this likely reflects lower overall sensitivity rather than frequent lapses (***Figure S8***), as the “easy” trials may not have been as easily discriminable for this participant.

**Table S4.** Catch trial performance summary across all sessions. The proportion correct reflects the total number of correct responses divided by the total number of catch trials. Lower and upper bounds indicate the participant’s lowest and highest session-level performance, respectively.

| Participant | ME | SG | DK | BH | FM | HG | FW | CH |
| --- | --- | --- | --- | --- | --- | --- | --- | --- |
| Proportion correct | 0.996 | 0.996 | 0.948 | 0.992 | 0.998 | 0.988 | 0.988 | 0.998 |
| Lower bound | 0.977 | 0.974 | 0.868 | 0.975 | 0.975 | 0.954 | 0.957 | 0.980 |
| Upper bound | 1.000 | 1.000 | 1.000 | 1.000 | 1.000 | 1.000 | 1.000 | 1.000 |

## Appendix 5 Simulated observer

To evaluate how well thresholds read out from the WPPM fits aligned with those estimated via Weibull fits, we simulated a dataset with a known ground truth. The following subsections outline the key steps in this process.

### Appendix 5.1: Derivation of the comparison stimuli at threshold on the isoluminant plane

We used CIELab Δ*E*94 as the ground truth metric for deriving color discrimination performance. For any given reference color and any given chromatic direction, both were affine-transformed from the model space to the RGB space. The RGB values were then converted to CIE 1931 XYZ and then to CIELab space, where Δ*E* computations were performed. In the XYZ-to-Lab transformation, we used the monitor gray point (*R* = *G* = *B* = 0.5) as the reference white. We then searched along each chromatic direction in the RGB space to find a comparison stimulus **x**_1_ = (*x*_1,dim1_, *x*_1,dim2_) such that Δ*E* in CIELab was equal to 2.5 (***Figure S15***A). This procedure was repeated across multiple directions. The resulting comparison stimuli were then mapped back into the model space, where we fit an ellipse to define the iso-distance contour (***Figure S15***B).

**Figure S15.**
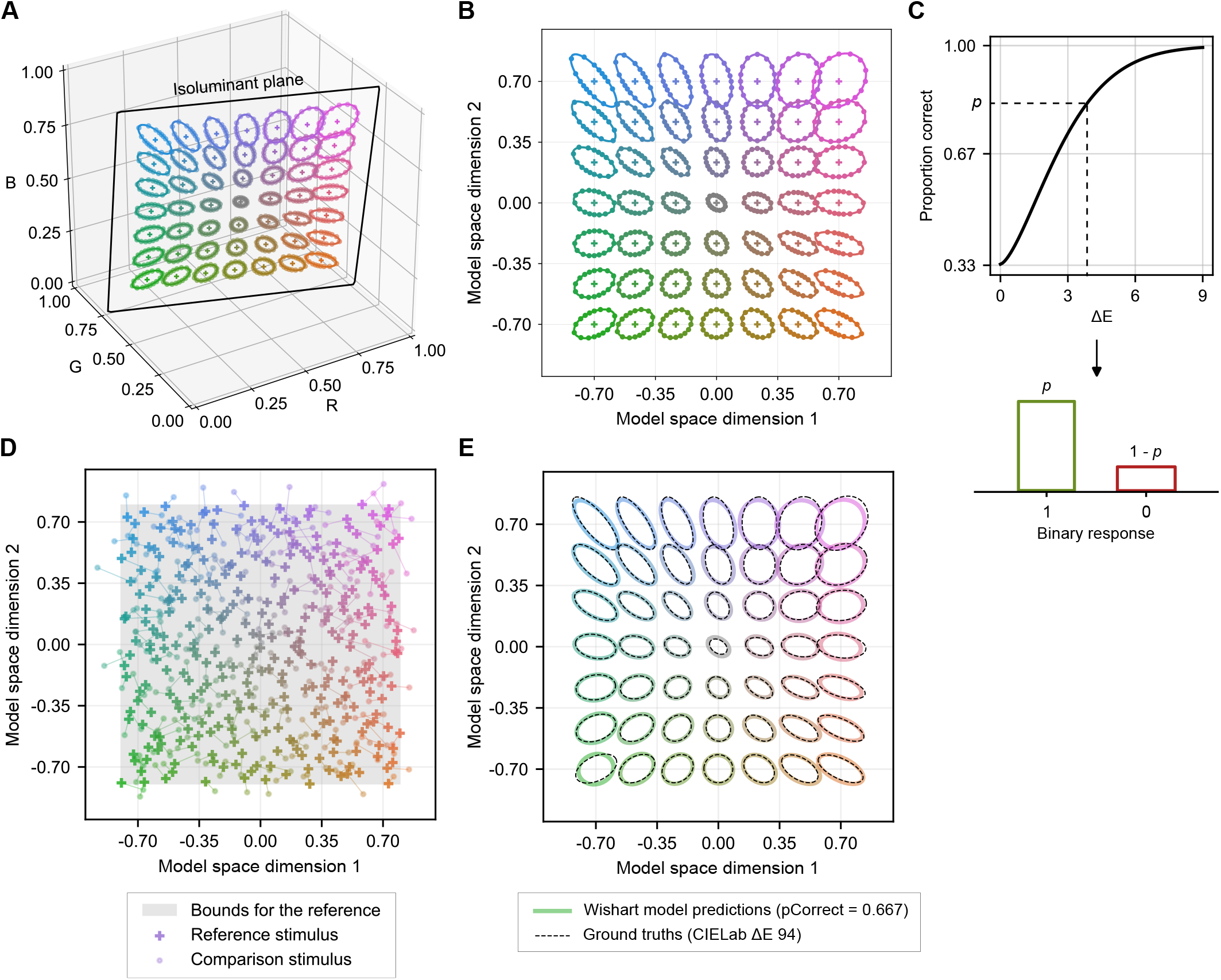
Derivation of the ground-truth Wishart fits based on CIELab Δ*E*94. (A–B) Comparison stimuli at the iso-distance contours in the isoluminant plane, shown in both RGB and model spaces. Note that the reference grid and fixed set of directions shown here are for illustration only; the actual sampling did not use a fixed grid or evenly spaced chromatic directions. (C) The Weibull psychometric function used to simulate binary (correct or incorrect) responses given Δ*E* values. (D) Sampled reference-comparison stimulus pairs. Reference colors and chromatic directions were sampled using Sobol’ sequences, and comparison stimuli were jittered around the iso-distance contour. A total of 18,000 trials were simulated; only the first 200 are shown here for clarity. (E) Comparison between readouts from the WPPM fit and from CIELab Δ*E*94. The WPPM fit was subsequently treated as the ground truth for simulating AEPsych and validation trials.

### Appendix 5.2: Simulation of trials near threshold contours

To introduce some variability, we added bivariate Gaussian noise to each comparison stimulus at the iso-distance contour in the model space. The noise standard deviation was proportional to the Euclidean distance between the reference stimulus **x**_0_ and the comparison stimulus **x**_1_. The jittered comparison stimulus 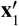 was computed as:

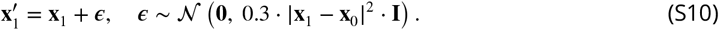

We modeled performance using a Weibull psychometric function, which took the Δ*E* between the reference and jittered comparison stimuli as input and returned the predicted percent correct:

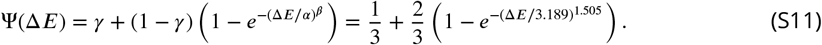

The values of *α* and *β* were selected such that the psychometric curve returns 66.7% correct when Δ*E* = 2.5 (***Figure S15***C). A binary (correct or incorrect) response was sampled from a Bernoulli distribution using this predicted probability:

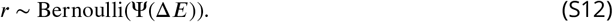

The comparison stimuli were selected to be near threshold, while the reference stimuli and chromatic directions were Sobol’ sampled to ensure uniform coverage of the model space without repeated trials (***Figure S15***D). In total, we simulated 18,000 trials.

### Appendix 5.3: Fit the WPPM and treating the model fits as the ground truth

We fitted the WPPM to the full set of 18,000 trials in the model space, and treated the resulting fit as the ground truth for simulating performance for both AEPsych and validation trials (***Figure S15***E, color lines). We chose to use the WPPM fit as the ground truth—rather than percent-correct performance derived from CIELab Δ*E*94 with a Weibull psychometric function—because our goal here was to evaluate how well the WPPM can recover ground truth that is itself described by the WPPM.

**Figure S16.**
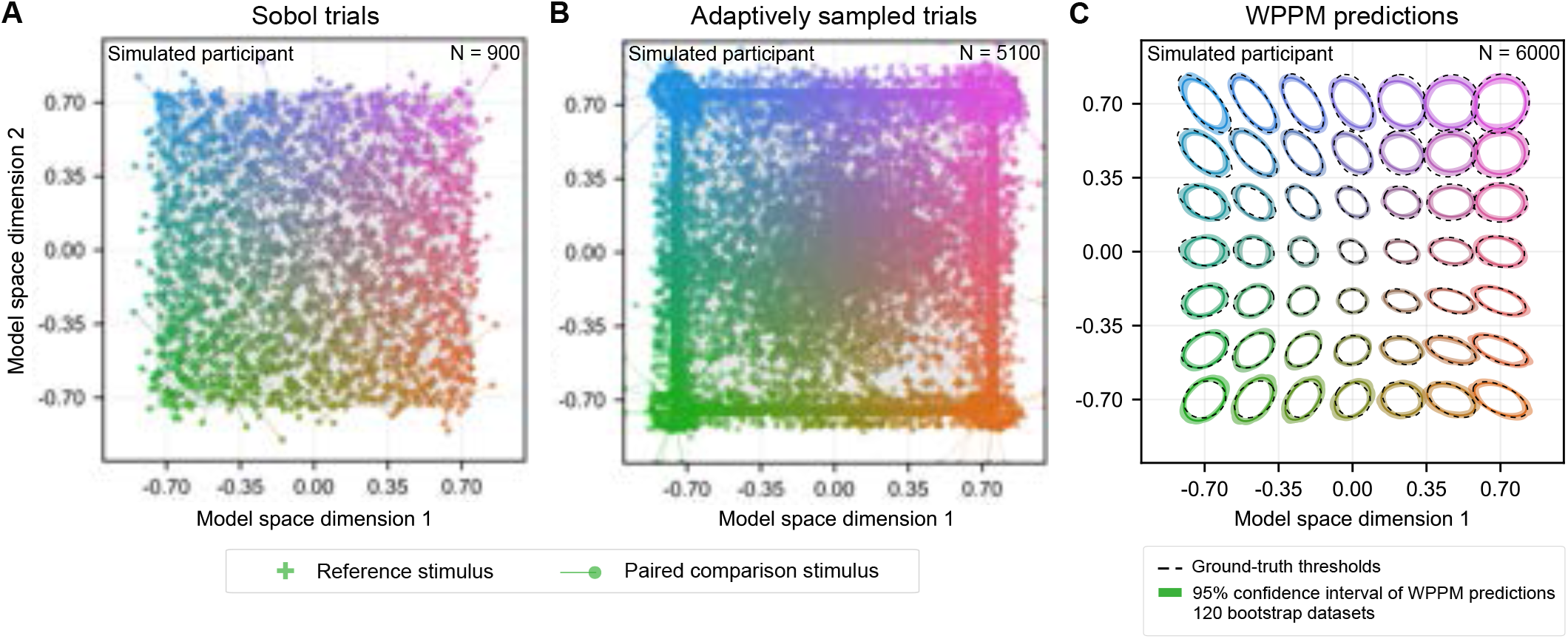
AEPsych-driven trials and WPPM readouts for a simulated observer. Note that the ground-truth thresholds shown in (C) is the same WPPM readouts from ***Figure S15E***.

**Figure S17.**
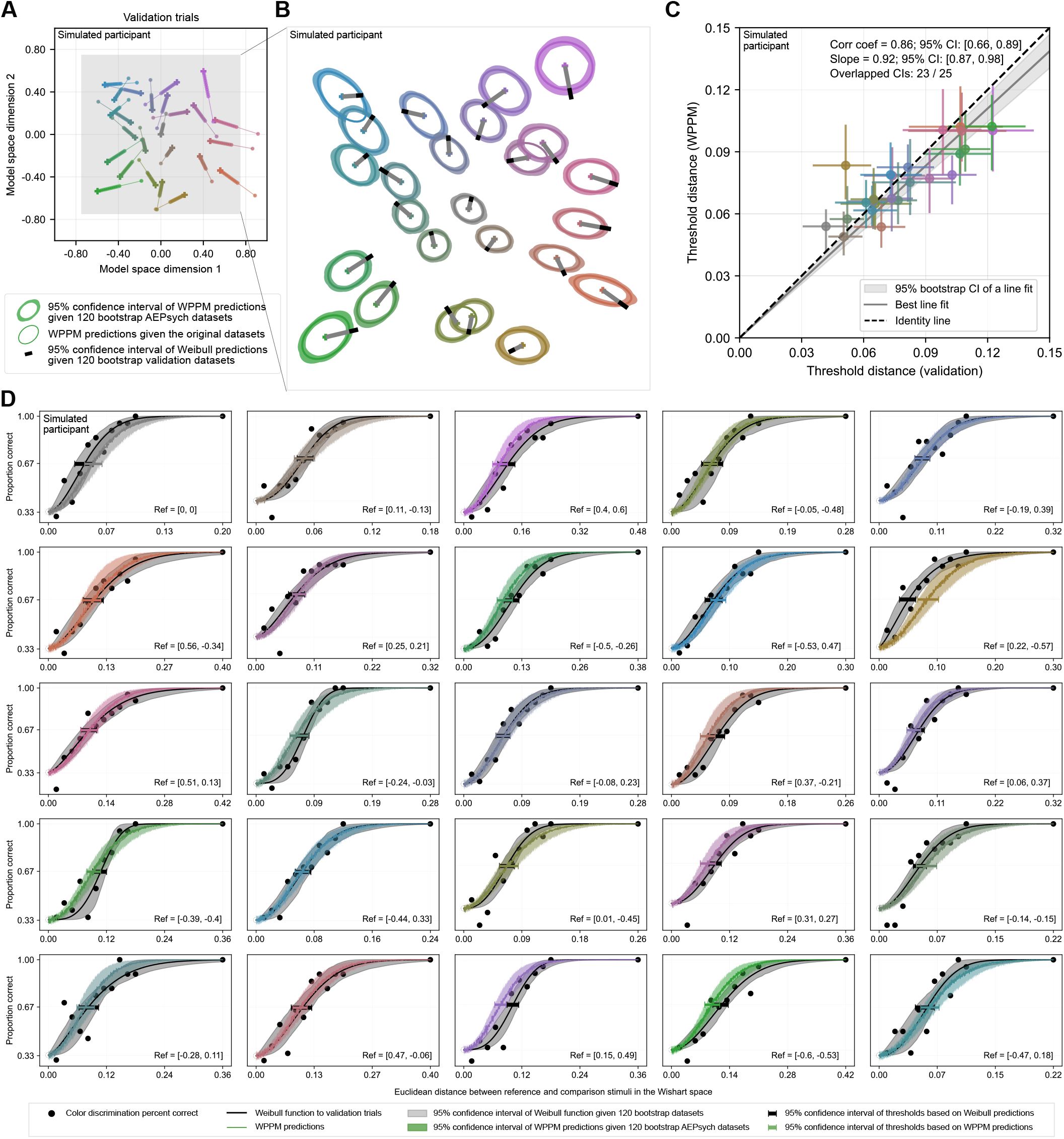
Validation trials and WPPM readouts for a simulated observer.

**Table S5.** Linear regression results for the simulated dataset.

| Predictor | Term | Coef | Std Err | *t* | *p* | [0.025, 0.975] CI | *R*<sup>2</sup> |
| --- | --- | --- | --- | --- | --- | --- | --- |
| Absolute difference of angles | intercept | -0.007 | 0.005 | -1.532 | 0.139 | [-0.017, 0.003] | 0.028 |
|  | slope | 0.000 | 0.000 | 0.812 | 0.425 | [-0.000, 0.000] |  |
| Aspect ratio | intercept | -0.007 | 0.012 | -0.579 | 0.568 | [-0.033, 0.019] | 0.003 |
|  | slope | 0.002 | 0.008 | 0.263 | 0.795 | [-0.014, 0.018] |  |
| Validation thresholds | intercept | 0.028 | 0.006 | 4.429 | 0.000 | [0.015, 0.041] | 0.547 |
|  | slope | -0.393 | 0.075 | -5.273 | 0.000 | [-0.547, -0.239] |  |

### Appendix 5.4: Fit the WPPM to simulated AEPsych trials

Based on the ground-truth WPPM fit, we simulated 900 Sobol’ trials (***Figure S16***A), followed by 5,100 adaptively sampled trials using AEPsych (***Figure S16***B), just like the design for the actual experiment. For each pair of reference and comparison stimuli, we approximated percent-correct performance using Monte Carlo simulation (N = 2,000), and generated binary responses by drawing from a Bernoulli distribution. We then fit the WPPM to this simulated dataset. To approximate the variability of the WPPM readouts, we bootstrapped the data 120 times, maintaining the same Sobol’-to-adaptive trial ratio within each bootstrapped dataset. As shown in ***Figure S16***C, the WPPM was able to reliably recover the ground-truth model, with only minor deviations. This good agreement provides provides context for the analyses in the following subsections, which are then compared with the corresponding analyses of the human data.

### Appendix 5.5: Validation trials and Weibull predictions

In addition to the 6,000 AEPsych trials, we also simulated 6,000 validation trials, mirroring the design of the actual experiment. Unlike the experimental design, these validation trials were simulated separately rather than interleaved, since sequential effects or perceptual learning are not factors in simulation. The WPPM thresholds confidence intervals agreed with 23 of 25 validation threshold confidence intervals (***Figure S17***). A linear regression fit to the validation thresholds (x-axis) and WPPM thresholds (y-axis) yielded a slope of 0.94 and a correlation coefficient of 0.83. These values fall within the range observed for human data (Appendix 4.1).

### Appendix 5.6: Statistical analysis of residuals between WPPM readouts and validation thresholds

We applied the same statistical analysis to the simulated data as we did for the human data (***subsection***). Consistent with the human results (***Figure S14***), we found no strong evidence that residuals systematically varied with the orientation or shape of the elliptical threshold contours read out from the WPPM fits. However, we did observe a significant negative correlation between the residuals and the magnitude of the validation thresholds (slope = −0.393, *t*(23) = −5.273, *p* < 0.001, *R*^2^ = 0.547; ***Figure S18***; ***Table S5***). As noted earlier, the size of this bias is small compared to the overall range of validation thresholds.

### Appendix 5.7: Comparison between WPPM estimates and simulation ground truth

To evaluate whether the WPPM readouts systematically deviated from the ground truth, we sampled thresholds over a fine grid of reference locations (15 × 15 points evenly spaced between –0.7 and 0.7 in model space) and compared them with the corresponding ground-truth thresholds. As a comparison metric, we used the Bures-Wasserstein (BW) distance (***Bhatia et al., 2019***), which quantifies the dissimilarity between two positive semi-definite covariance matrices, Σ_1_ and Σ_2_. Intuitively, it captures the “effort” required to morph one ellipse into another. Mathematically, the BW distance is defined as

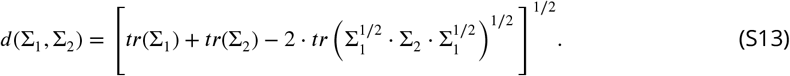

**Figure S18.**
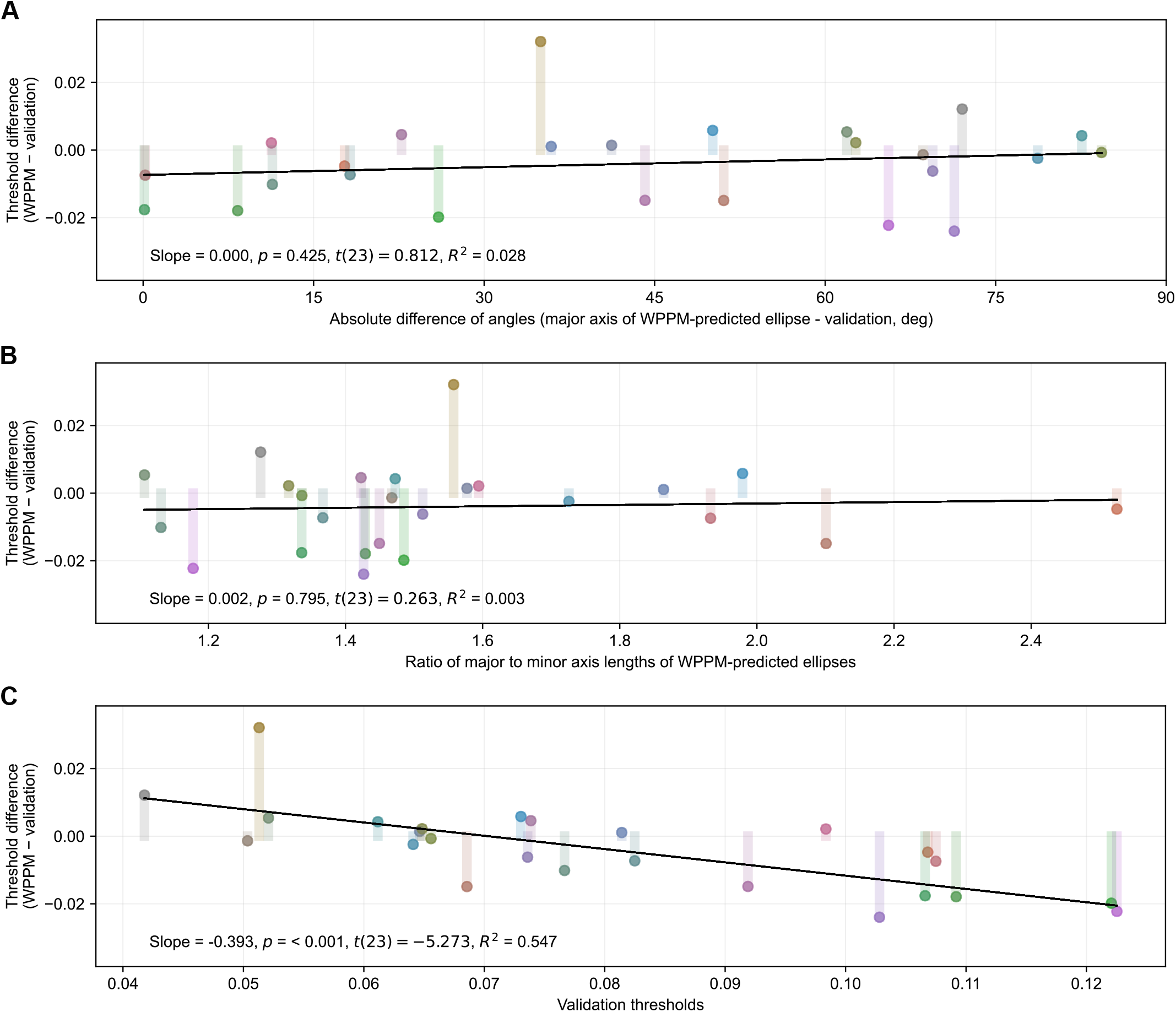
Threshold residuals for a simulated dataset. For all panels, color codes for the surface color of the reference stimulus, and the y-axis limits are set to ± the mean of the validation thresholds. (A) Residuals as a function of the absolute angular difference between the major axis of the elliptical threshold contours read out from the WPPM fits and the chromatic direction of the validation condition. (B) Residuals as a function of the aspect ratio (major/minor axis) of the WPPM threshold contours. (C) Residuals as a function of thresholds estimated from validation trials.

The BW distance is non-negative and equals zero only when the two matrices being compared are identical. Smaller distances indicate greater similarity between the threshold ellipses.

The results showed that BW distance generally increased as the reference color moved farther from the achromatic point (***Figure S19***A), suggesting that the WPPM has more difficulty accurately capturing large threshold contours in regions with higher internal noise. To provide a benchmark for what constitutes a substantial mismatch, we computed the BW distance between each ground-truth ellipse and a circle with radius being the largest major axis length among all ground-truth ellipses. The maximum of these values served as a reference point (shown as the upper limit of the color bar in ***Figure S19***A). Overall, the mismatches observed in our simulations were modest— well below the level expected if the model were fundamentally mischaracterizing the threshold shapes.

**Figure S19.**
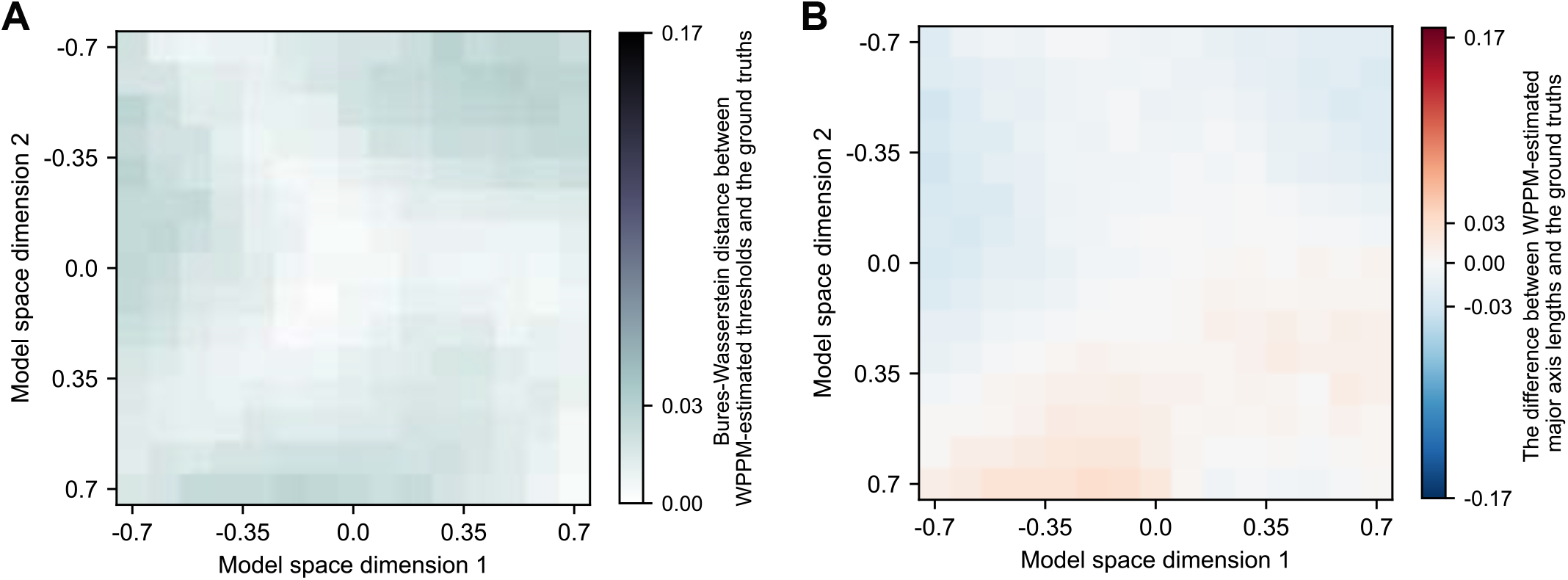
Deviation of WPPM estimates from the ground truth. (A) BW distance between WPPM-estimated thresholds and the ground-truth ellipses. The upper limit of the color map (0.17) corresponds to the maximum BW distance between each ground-truth ellipse and a reference circle whose radius equals the largest major axis length among all ground-truth ellipses. The maximum BW distance between WPPM estimates and the ground truth (0.03) is substantially lower than this reference value. (B) Difference in major axis length between WPPM-readouts and ground-truth ellipses. The colormap limits (±0.17) reflect the ± maximum ground-truth major axis length. Again, the maximum deviation observed (0.03) is small relative to this range.

We also examined differences in the estimated major axis lengths. The WPPM showed slight underestimation in the upper region and overestimation in the lower region of the space (***Figure S19***B; also apparent but small in ***Figure S16***C). Similar to the BW analysis, these deviations were relatively small compared to the overall range of ground-truth values. Together, these results indicate that the WPPM provides a close and robust approximation of the true threshold contours, with only minor local deviations.

## Appendix 6 Comparison with MacAdam ellipses (1942)

**Figure S20.**
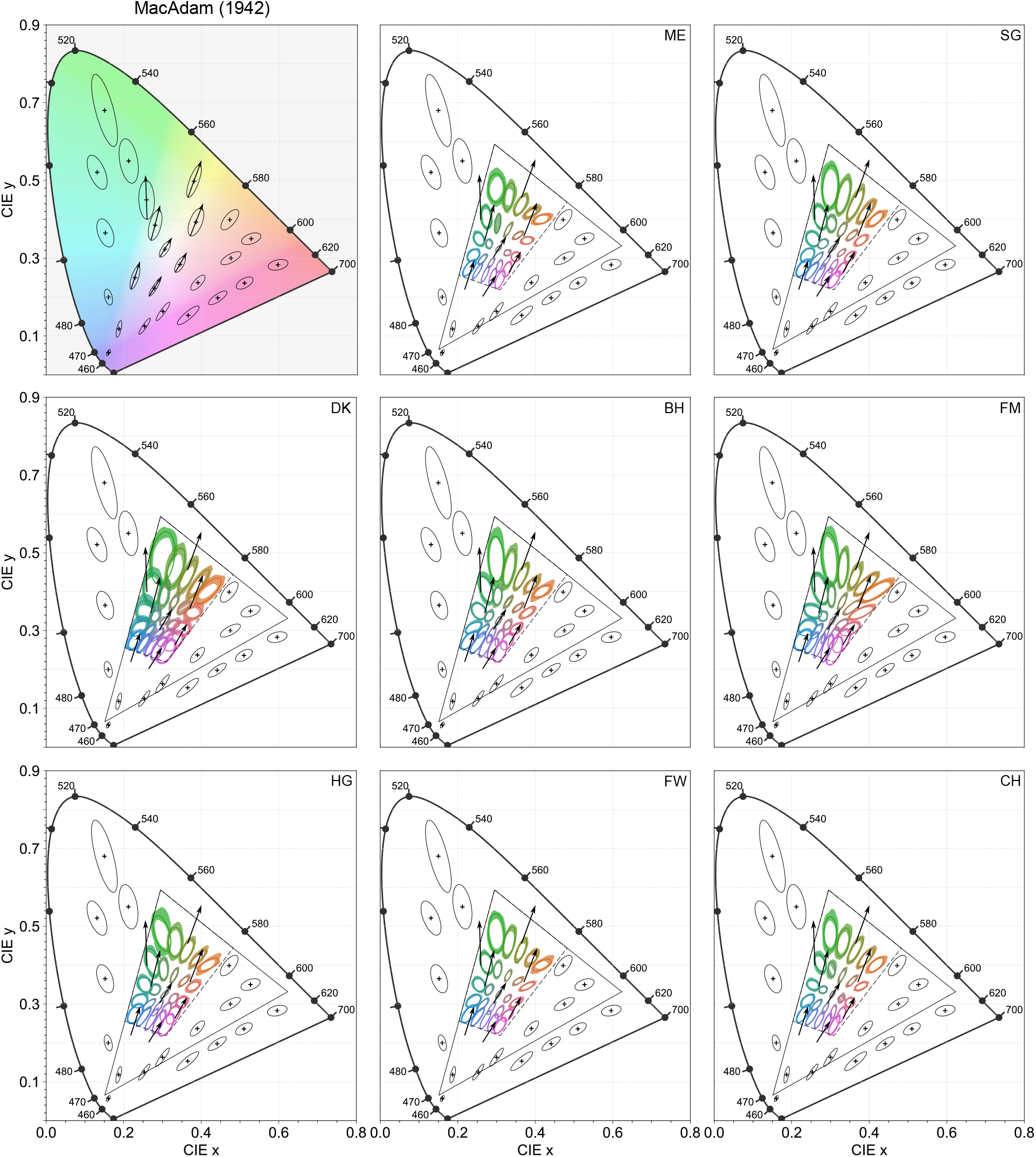
Comparison with ***MacAdam 1942***. First panel: MacAdam’s original threshold ellipses, magnified 10× for visualization. Rest panels: 66.7% threshold contours (colored lines) measured from all participants in our study and transformed from the model space into the CIE 1931 chromaticity diagram. Colored shaded regions indicate 95% confidence intervals computed from 120 bootstrapped datasets. Reference stimuli were sampled from a 5 × 5 grid spanning [−0.7, 0.7] along each dimension of the model space. To reduce visual clutter, MacAdam ellipses falling within the gamut of our isoluminant plane (parallelogram) are shown only by arrows indicating their major axes. For visual comparability, our ellipses are magnified 2× to approximately match the scale of MacAdam’s data. The triangle indicates the monitor gamut.

## Appendix 7 Comparison with Danilova & Mollon (2025)

In this section, we compare our measurements with those from ***Danilova and Mollon 2025*** by transforming our results into the chromaticity space used in their study—a scaled version of the MacLeod–Boynton space (***MacLeod and Boynton, 1979***). While a direct transformation path exists from our model space to theirs (model space → RGB → LMS → MacLeod–Boynton → scaled MacLeod–Boynton), it assumes that the adaptation point and isoluminant plane are identical between the two studies, which is not the case. To account for these differences, we instead took a detour through the DKL space (***Derrington et al., 1984***), where cone-opponent mechanisms are explicitly defined and adaptation is more easily controlled. Specifically, we followed the transformation chain: model space → RGB_*us*_ → LMS_*us*_ → ΔLMS_*us*_ → DKL → ΔLMS_*dm*_ → LMS_*dm*_ → MacLeod–Boynton → scaled MacLeod–Boynton. Here, the subscript “*us*” refers to values computed using our study’s cone fundamentals, luminosity function and adaptation point, while “*dm*” denotes those based on ***Danilova and Mollon 2025***. This approach allowed us to approximate how our stimuli would be represented in their perceptual framework, enabling visual comparison of the threshold contours. The comparison reveals a general qualitative agreement between their measurements and ours (***Figure S21***).

**Figure S21.**
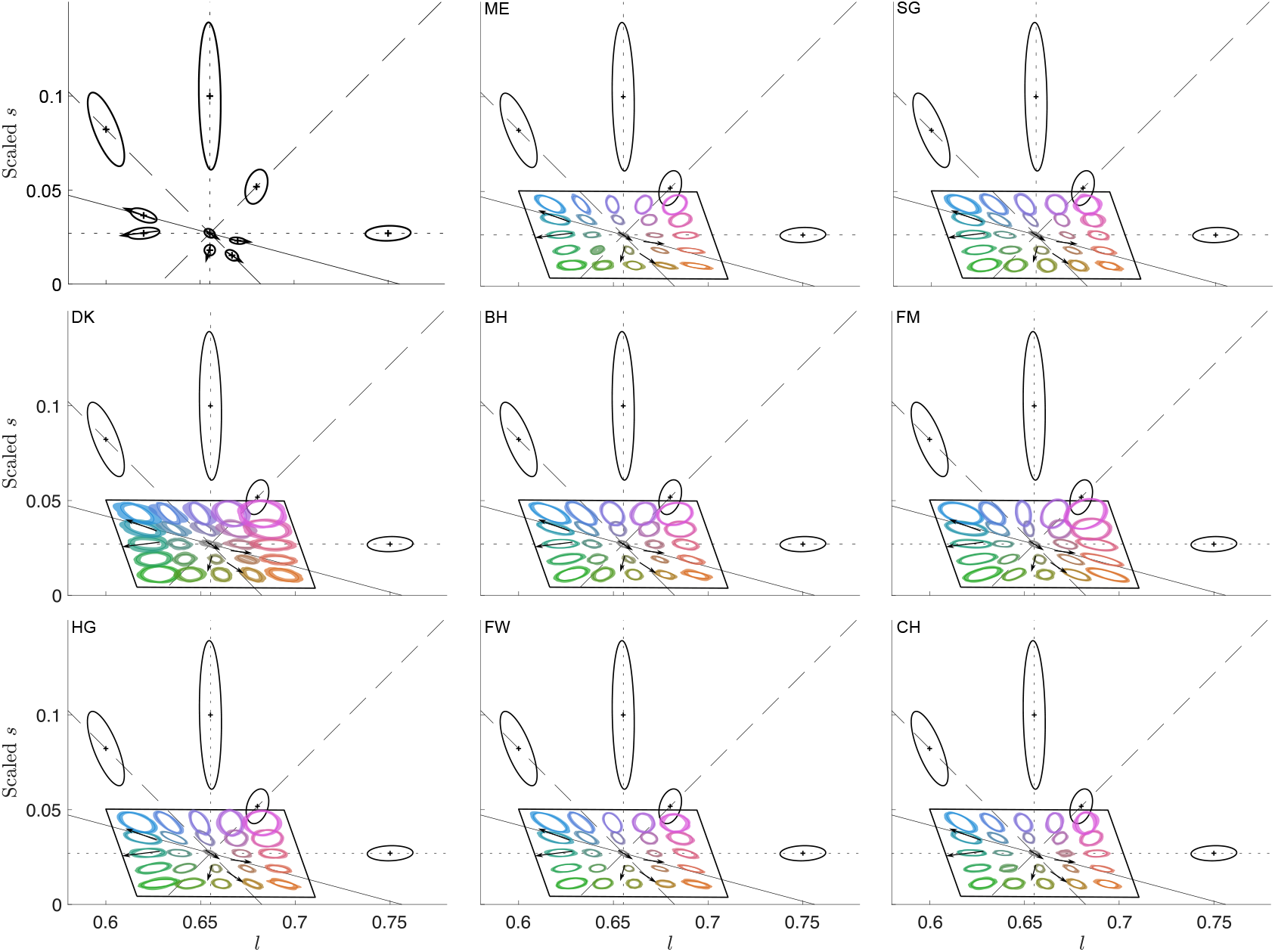
Comparison with ***Danilova and Mollon 2025*** in the scaled MacLeod–Boynton space. Top left: threshold contours from their study (black ellipses), enlarged by 4×. Remaining panels: threshold contours from all participants (colored ellipses). We sampled a grid of reference points evenly spaced from –0.7 to 0.7 (5 steps) in our model space, read out the corresponding threshold contours, and transformed them into the same scaled MacLeod–Boynton space. The parallelogram indicates the gamut of the isoluminant plane. To reduce visual clutter, ellipses from Danilova & Mollon that fall within our gamut are represented by red arrows indicating only their major axes. For visual comparability, our ellipses are enlarged by 1.5× to roughly match the size of those in their study.

## Appendix 8 Comparison with Krauskopf & Gegenfurtner (1992)

We compared our threshold estimates with those reported by ***Krauskopf and Gegenfurtner 1992***. To do so, we transformed our estimates into the color space used in their measurements through a series of steps. We first read out, for each participant, the threshold contour at the achromatic reference color in the model space, which was then transformed to the DKL space (***Derrington et al., 1984***). We then normalized the DKL cardinal axes so that the threshold contour at the achromatic reference had unit length along both axes. This normalized space—referred to here as the stretched DKL space—is the coordinate system in which ***Krauskopf and Gegenfurtner 1992*** conducted their measurements. Finally, we converted the 16 reference stimuli used in their study into our model space, read out the corresponding threshold ellipses, and transformed them into this stretched DKL space to enable direct comparison (***Figure S22***). We observed generally good agreement between their measurements and those of some participants (e.g., CH and FM). Notably, however, individual differences are evident, particularly in the upper-right quadrant of the stretched DKL space (***Figure S23***). In addition, at the adapting chromaticity our ellipses are consistently rotated relative to the DKL axes, as noted in the main text.

**Figure S22.**
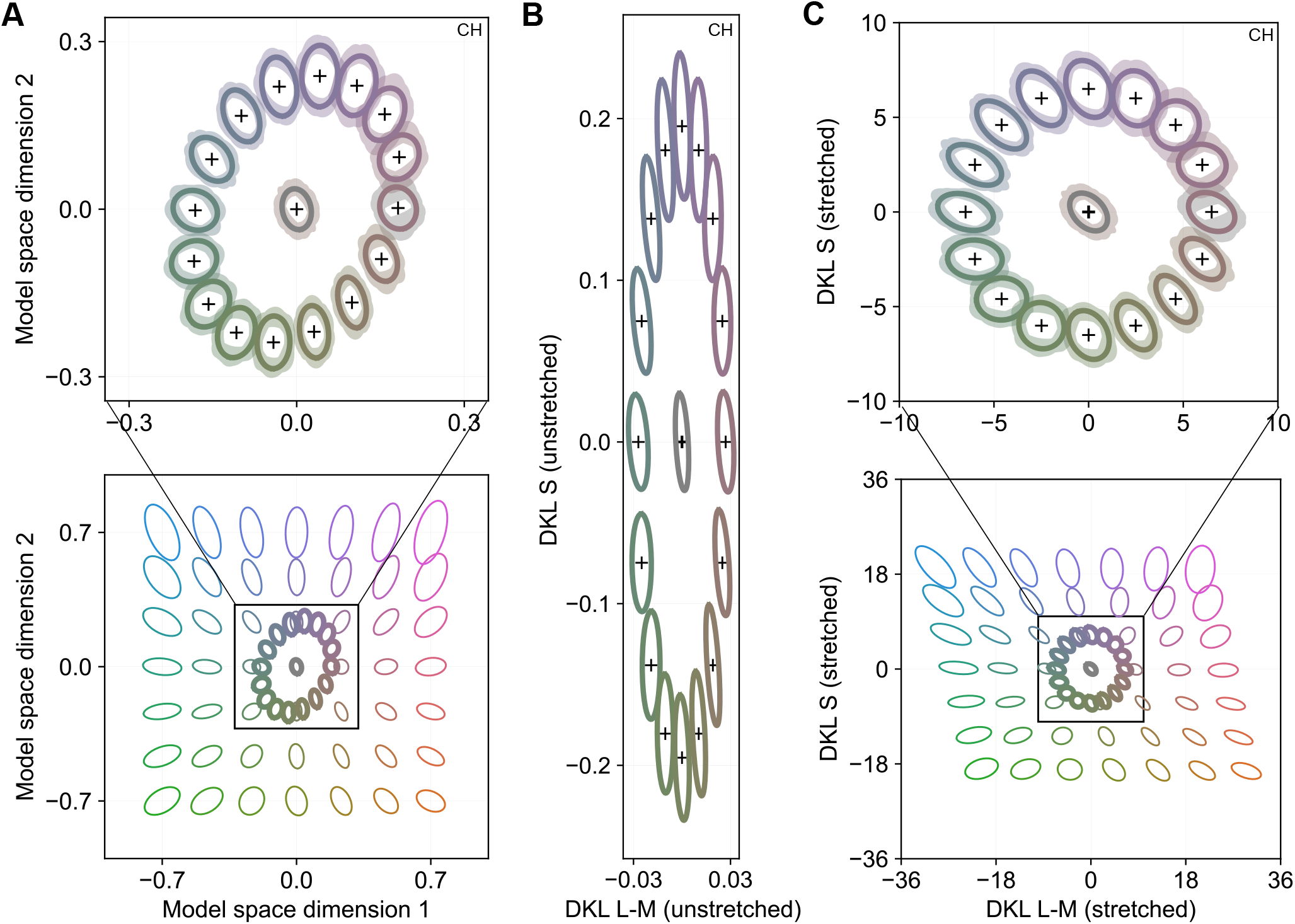
Transformation from the model space to a stretched DKL space used in ***Krauskopf and Gegenfurtner 1992*** for participant CH. (A) Model space. Threshold contours were read out in this space based on each participant’s WPPM fit. Notably, our data were collected on a much larger region of the isoluminant plane than they characterized. (B) The intermediate, unstretched DKL space. Transformations between this space and both the model space and the stretched DKL space are affine. (C) Stretched DKL space, in which the cardinal axes of the original DKL space are rescaled such that the threshold at the achromatic reference point is normalized.

**Figure S23.**
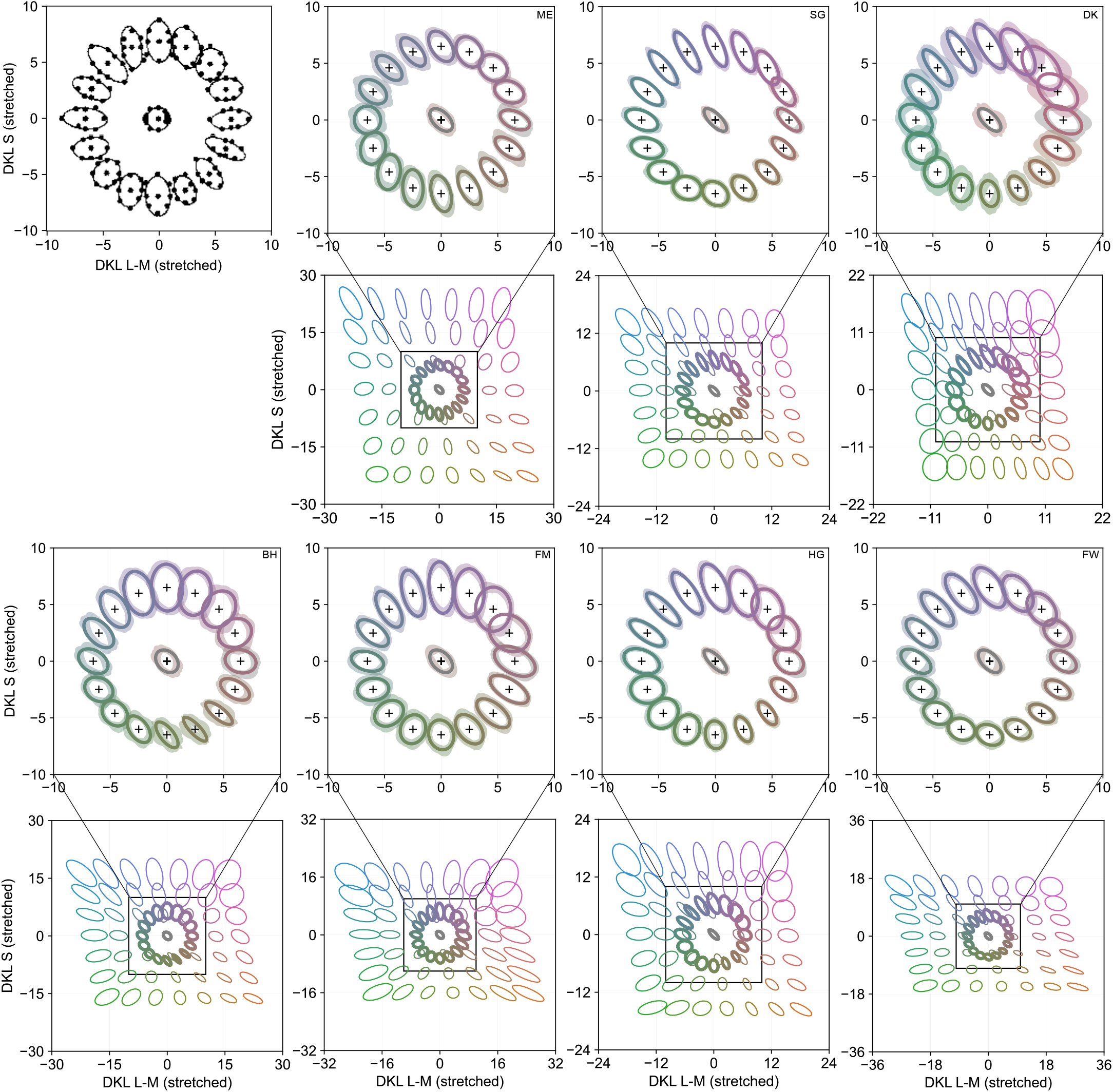
Comparison with ***Krauskopf and Gegenfurtner 1992*** for the remaining seven participants. Top left: original threshold contours reported by ***Krauskopf and Gegenfurtner 1992***, reproduced under Creative Commons CC BY-NC-ND 4.0). Remaining panels: 66.7% threshold contours (colored lines) for the remaining participants, transformed into the stretched DKL space using participant-specific scaling of the cardinal axes. Colored shaded regions indicate 95% confidence intervals computed from 120 bootstrapped datasets. All contours are plotted at their original sizes.

## Appendix 9 Comparison with CIELab Δ*E*76, Δ*E*94, Δ*E*00

**Figure S24.**
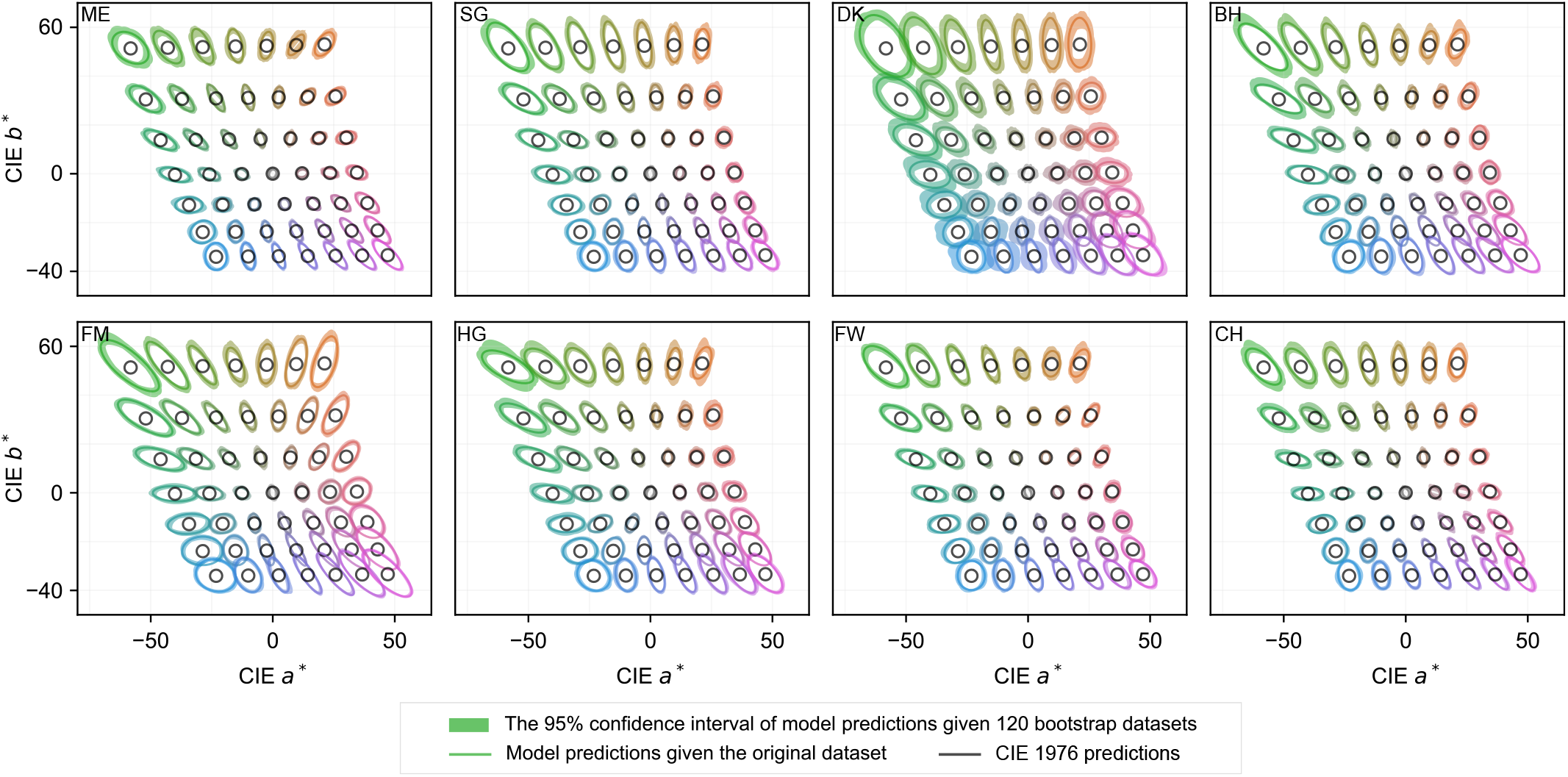
Comparison with CIELab Δ*E*76 color-difference (***Robertson et al., 1977***). Δ*E* values were converted to percent correct using a Weibull psychometric function, and threshold was defined as Δ*E* = 2.5, chosen to approximately match the scale of the measured thresholds in our data. Black contours represent the CIE predictions, whereas colored contours represent the measured thresholds transformed from the model space into CIELab space. Colored shaded regions indicate 95% confidence intervals computed from 120 bootstrapped datasets. Measured thresholds are shown at their original scales.

We obtained the elliptical thresholds on a grid of reference stimuli in the model space from the WPPM fits to participants’ data (***Figure S1***). We then transformed these ellipses from the model space to CIELab space. Specifically, values in the model space were first converted to RGB values via the affine transformation (Appendix 1.3). The resulting RGB values were converted to LMS via a transformation matrix (***Table S6***), computed using the 2^°^ cone fundamentals from ***CIE 2006***, then to XYZ using the color matching functions reported in ***CIE 2015***. Finally, the XYZ values were converted to CIELab using a Python implementation (***Taylor, 2017***). The adapting background (*R* = *G* = *B* = 0.5) was used as the reference white in the XYZ-to-Lab transformation. In addition, we computed threshold contours directly in CIELab space, defining them as iso-distance contours at a fixed perceptual distance of Δ*E* = 2.5.

**Table S6.** Transformation matrix from LMS to XYZ.

| $M_{LMS \rightarrow XYZ}$ | | |
| --- | --- | --- |
| 1.947 | -1.415 | 0.365 |
| 0.690 | 0.248 | 0.000 |
| 0.000 | 0.000 | 1.935 |

Comparisons revealed that the iso-distance contours from Δ*E*94 and Δ*E*00 provided reasonable approximations to our model-predicted thresholds (***Figure S25*** – ***Figure S26***), with only modest deviations. In contrast, the Δ*E*76 contours—despite their continued widespread use—diverged substantially from our measured thresholds (***Figure S24***).

**Figure S25.**
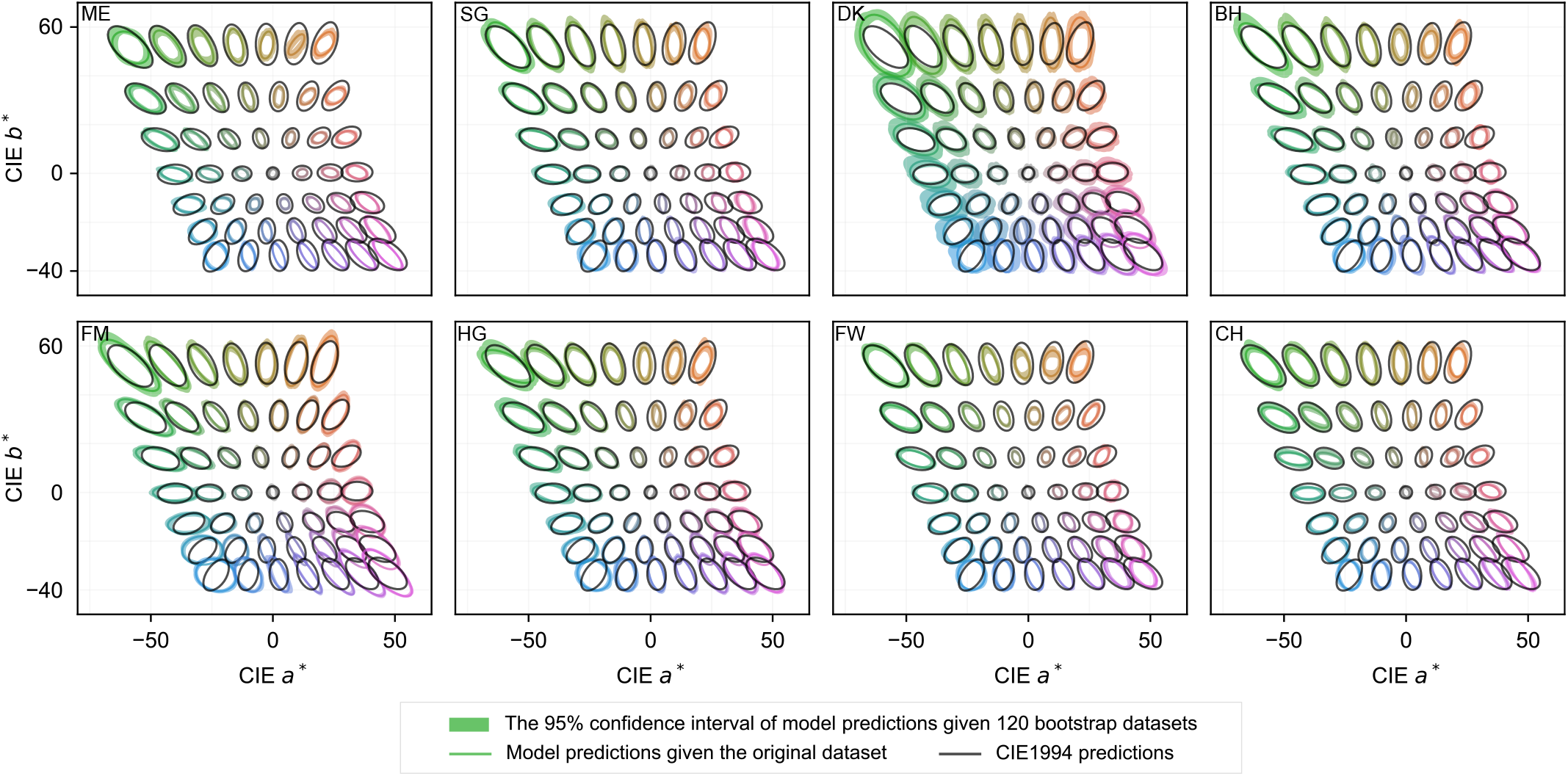
Comparison with predictions based on the CIELab Δ*E*94 color-difference metric (***McDonald and Smith, 1995***).

**Figure S26.**
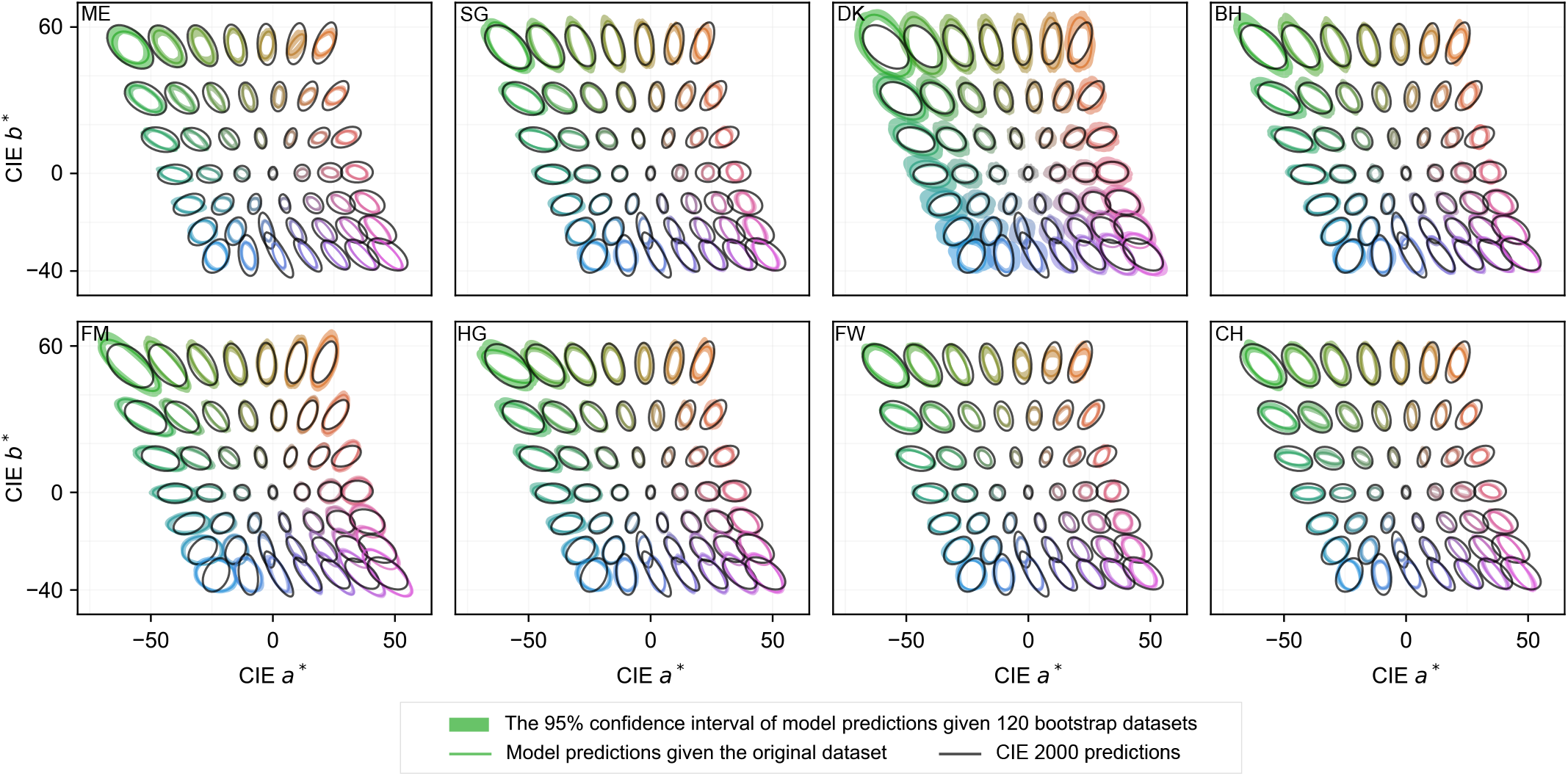
Comparison with CIELab Δ*E*00 color-difference metric (***Sharma et al., 2005***).

## Appendix 10 Hyperparameters that control the smoothness prior

### Appendix 10.1: The effects of *ϵ* and *γ* on WPPM predicted psychometric field

We assume that the internal noise limiting color discrimination varies smoothly across model space. Smoothness is implemented as the variance of the basis weights (***Equation 16***), where *ϵ* determines how rapidly the variance decreases with increasing polynomial order, and *γ* sets the overall variance scale (***Figure S27***).

To evaluate the influence of each hyperparameter, we first varied *ϵ* across a broad range while fixing *γ* = 0.0003. Each partipant’s dataset was divided into five folds for cross-validation. For each value of *ϵ*, we fit the WPPM to a training set consisting of four folds of the data, while leaving the held-out fold as the test set. Model parameters were obtained by minimizing the negative log posterior probability on the training data. To evaluate model performance on both the training and test sets, we computed the negative log likelihood (nLL) without the prior to enable a fair comparison.

After each fold had been treated once as the test set, we computed the mean and the full range of nLL across the five repetitions. As expected, the mean nLL evaluated on the training data decreased monotonically with increasing *ϵ*, reflecting improved fit with greater model flexibility. In contrast, the mean nLL evaluated on the held-out test data initially decreased and then gradually increased (***Figure S28***–***Figure S29***). This pattern reflects a tradeoff governed by the smoothness prior. When smoothness is enforced too strongly, the model produces an overly uniform threshold field that fails to capture the data. As the smoothness constraint is relaxed, predictive performance improves and remains relatively stable over a range of *ϵ* values. Beyond this range, further reductions in smoothness lead to increased variability in the estimated psychometric field across the five cross-validation repetitions.

We performed the same analysis to examine the influence of the hyperparameter *γ* on the estimated covariance field and observed the same qualitative pattern (***Figure S30***–***Figure S31***). Importantly, the hyperparameter values used in the main analyses (*ϵ* = 0.4 and *γ* = 0.0003) lie within the regime that balances oversmoothing against increasing variability.

**Figure S27.**
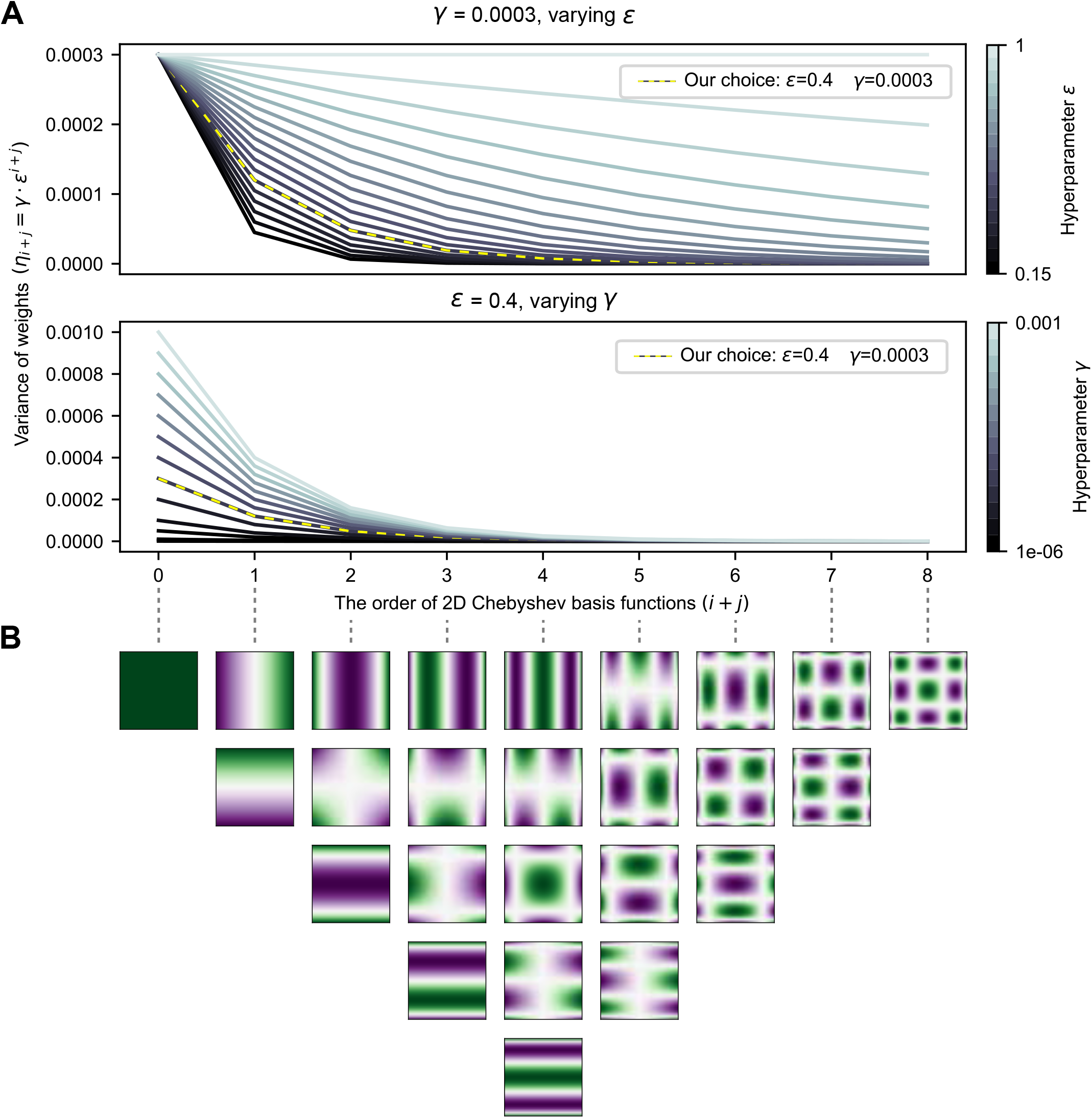
The effects of *ϵ* and *γ* on the variance of model weights. (A) Variance of the Chebyshev basis weights as a function of polynomial order (*i* + *j*). The top panel illustrates the effect of varying *ϵ* while holding *γ* fixed, whereas the bottom panel shows the effect of varying *γ* while holding *ϵ* fixed. The yellow dashed curve indicates the hyperparameter values used in the main analyses (*ϵ* = 0.4, *γ* = 0.0003). (B) Two-dimensional Chebyshev basis functions arranged in order of increasing total polynomial degree (*i* + *j*).

**Figure S28.**
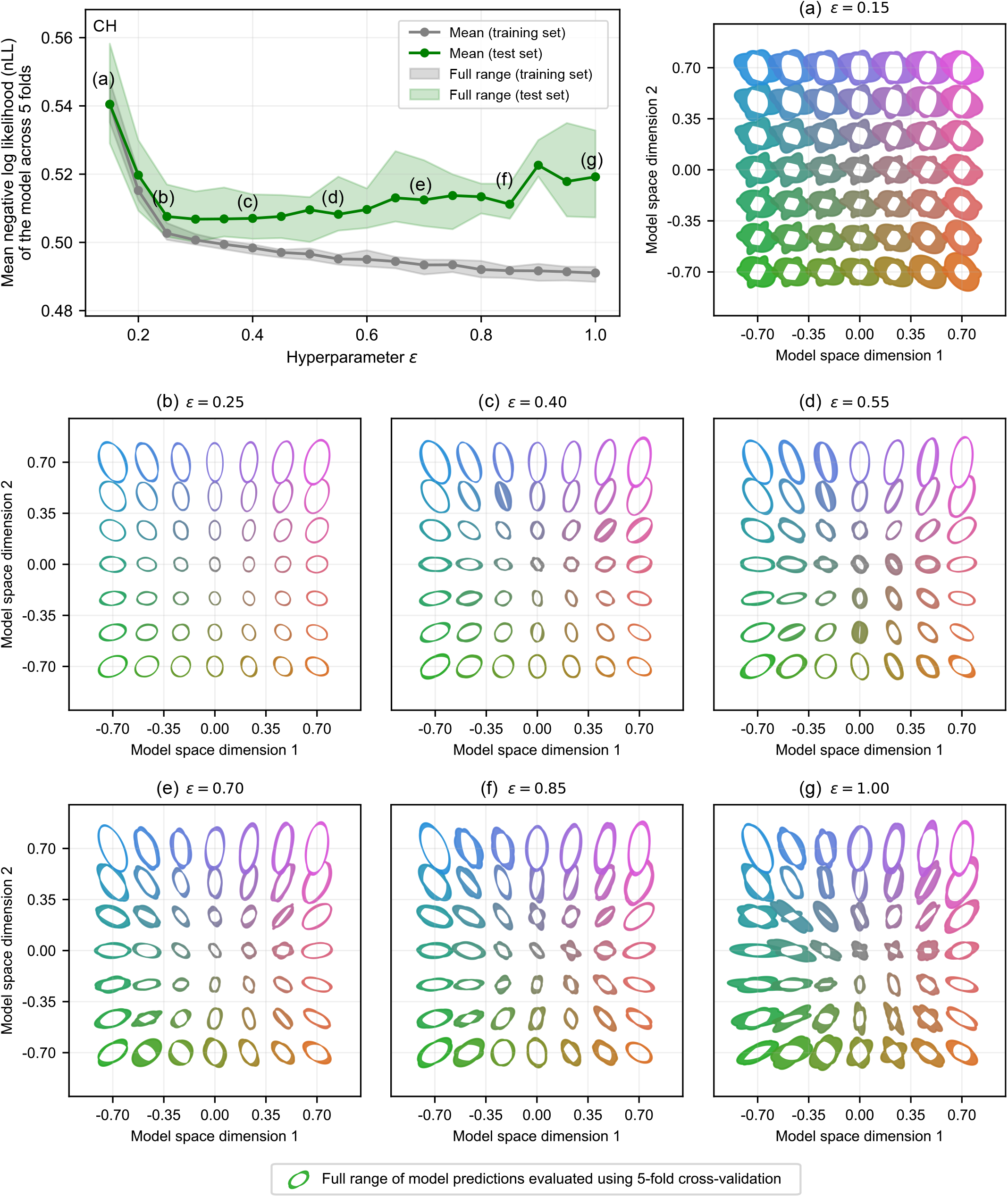
Effects of *ϵ* on WPPM-predicted psychometric field for a representative participant. Gray line and shaded region indicate the mean and full range of negative log likelihood (nLL) on the training set across five repetitions of five-fold cross-validation. Green line and shaded region indicate the mean and full range of nLL on the test set. Panels (a)–(g) show the model-predicted thresholds on a 7 × 7 reference grid for selected values of *ϵ*.

**Figure S29.**
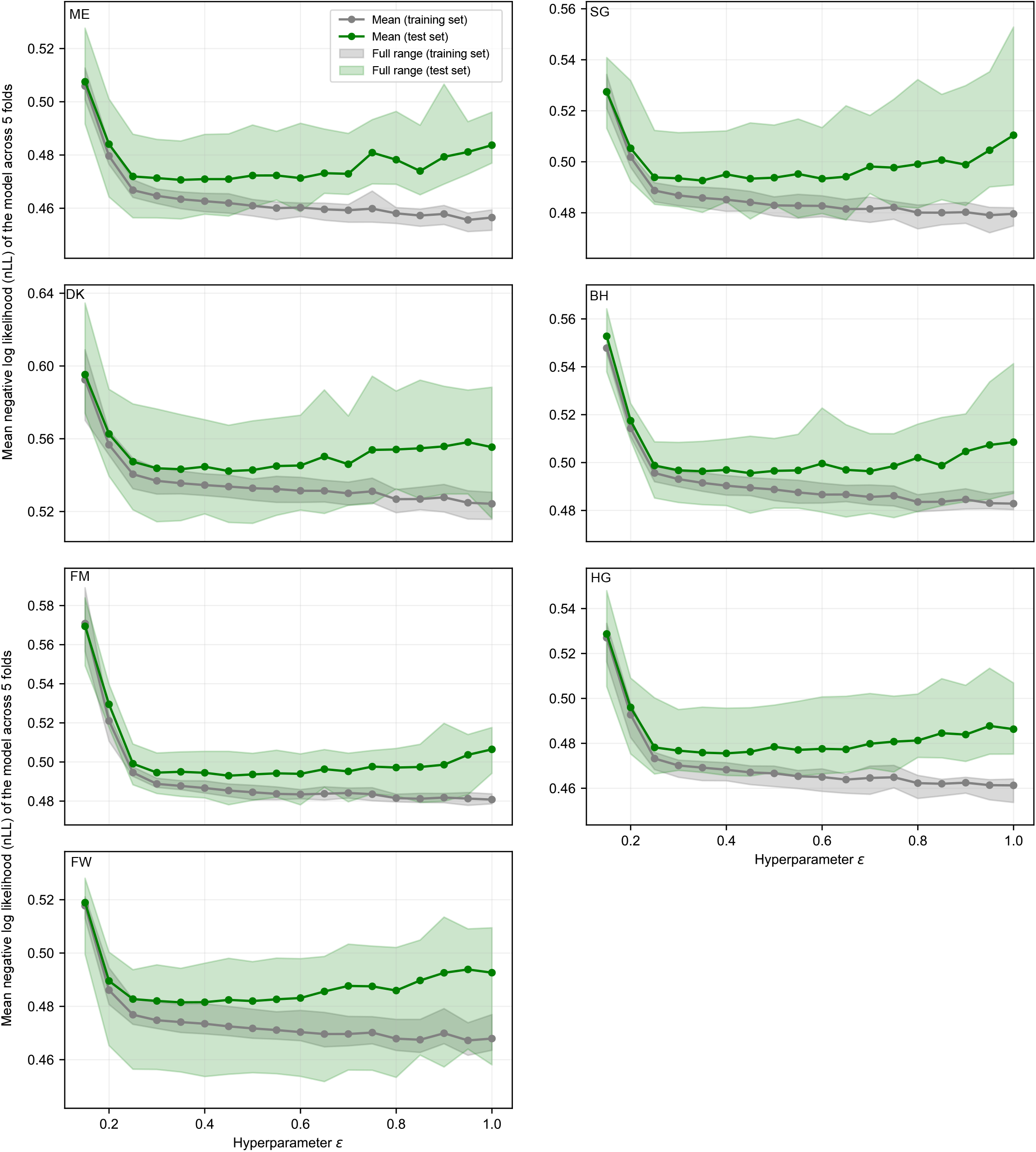
Effects of *ϵ* on model performance for the remaining seven participants.

**Figure S30.**
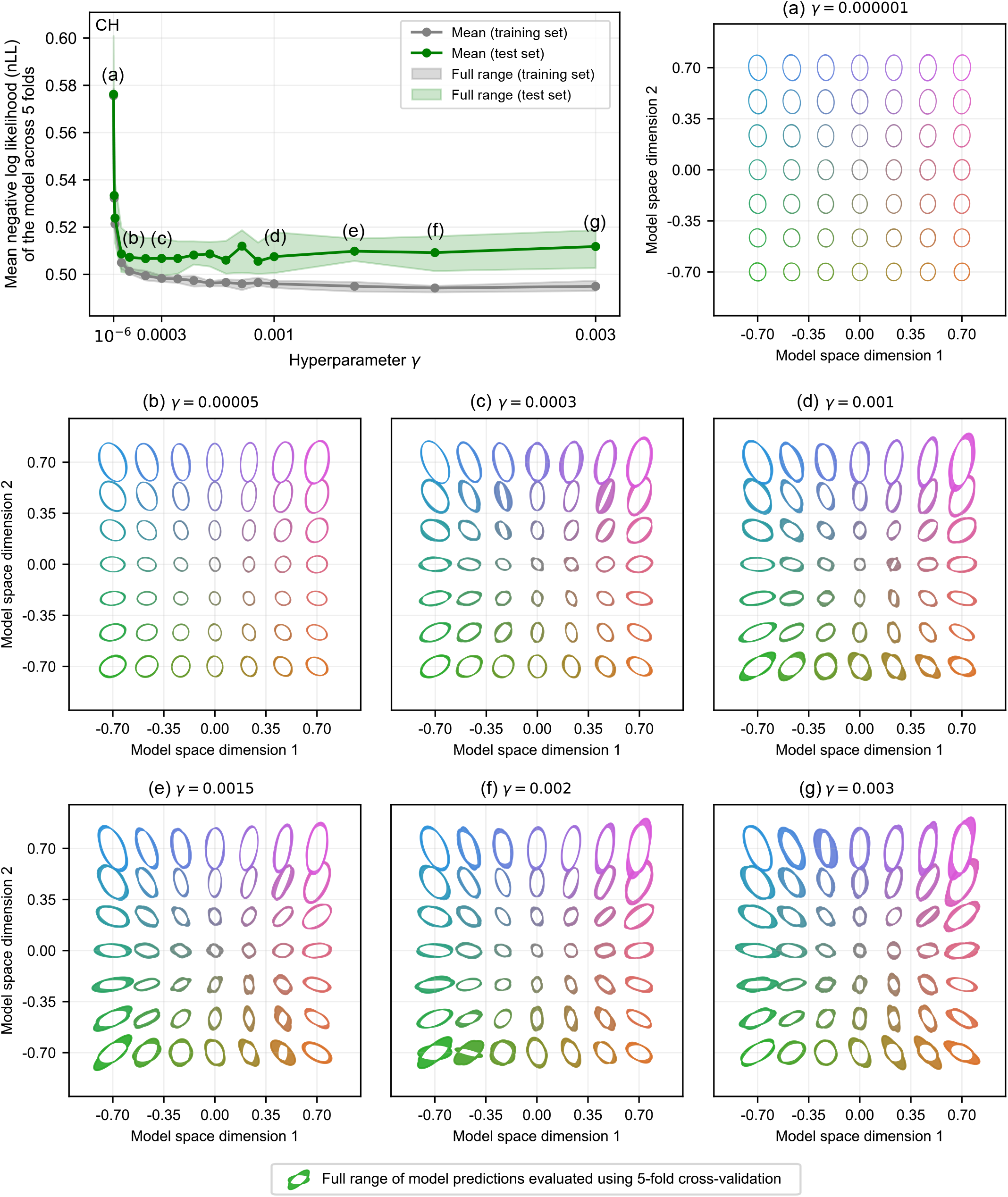
Effects of *γ* on WPPM-predicted psychometric field for a representative participant. Gray line and shaded region indicate the mean and full range of negative log likelihood (nLL) on the training set across five repetitions of five-fold cross-validation. Green line and shaded region indicate the mean and full range of nLL on the test set. Panels (a)–(g) show the model-predicted thresholds on a 7 × 7 reference grid for selected values of *γ*.

**Figure S31.**
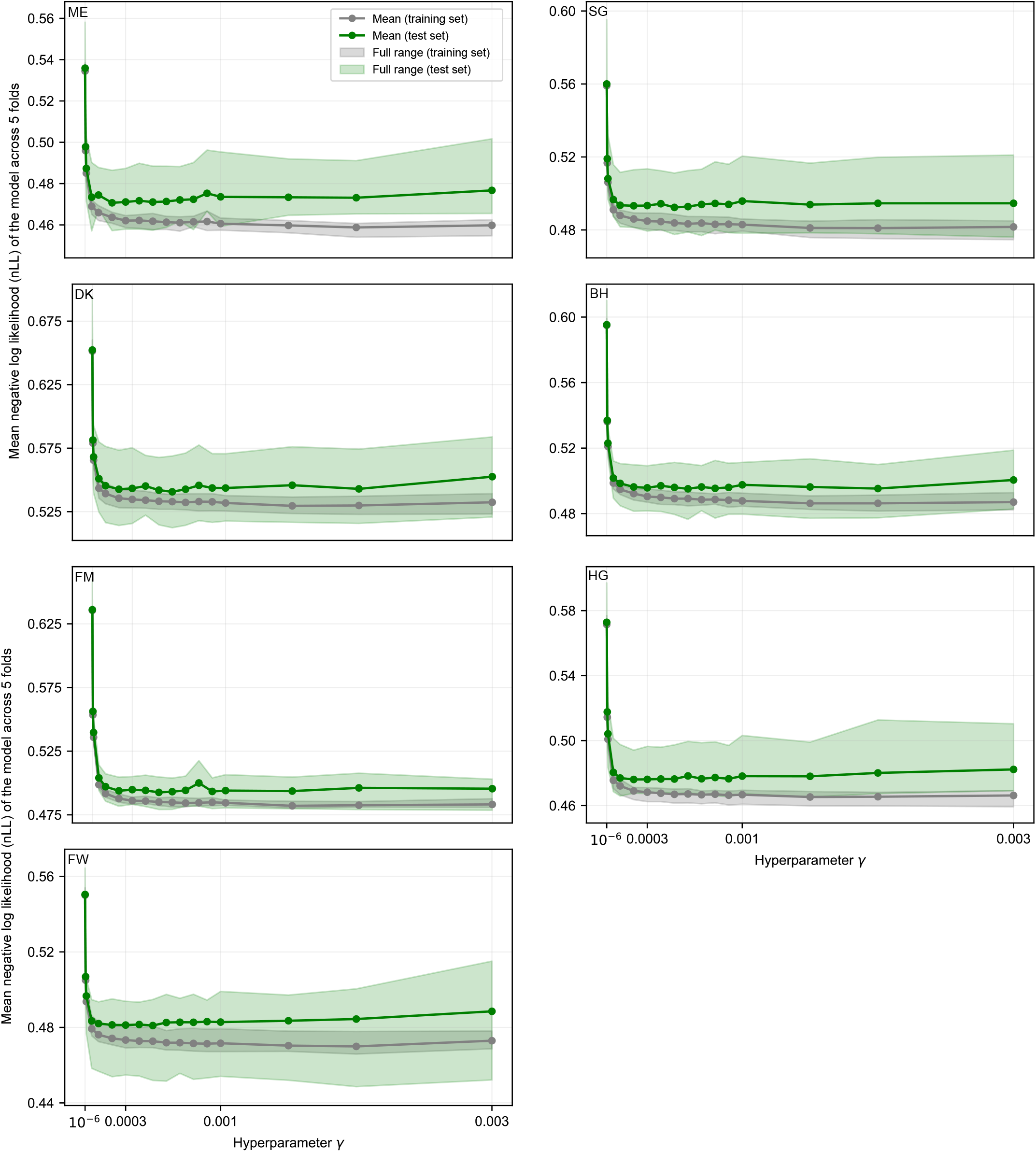
Effects of *γ* on model performance for the remaining seven participants.

### Appendix 10.2: The effects of *ϵ* on the WPPM-validation threshold residuals

We examined how *ϵ* influences the agreement between WPPM-predicted thresholds and validation thresholds while fixing the variance scale *γ* at 0.0003. We held *γ* constant because cross-validation results showed that beyond a certain value the nLL for both training and test sets plateaued, indicating that further changes in *γ* had only a minimal impact on the model predictions (***Figure S31***). Using the model fits reported in Appendix 10.1 for each value of *ϵ*, we computed predicted thresholds at the 25 validation conditions and quantified the residuals between WPPM predictions and validation thresholds, following the procedure described in Appendix 4.2.

To assess systematic patterns in these residuals, we fit a linear regression model with three predictors: (1) the absolute angular difference between the chromatic direction of the validation condition and the major axis of the contours derived from the WPPM fit, (2) the aspect ratio of the contours, and (3) the magnitude of the validation threshold. The regression slopes are summarized in ***Table S7***. Only the regression slopes are reported, as the intercepts were not as relevant.

Of the three predictors, the regression slopes relating threshold residuals to angular difference and to aspect ratio did not vary systematically with *ϵ*, showing no consistent or monotonic trend across different *ϵ*. In contrast, the regression slope relating threshold residuals to validation threshold exhibited a clear and systematic dependence on *ϵ*. As the smoothness prior became weaker (larger *ϵ*), the negative correlation between threshold residuals and validation threshold weakened. This pattern is consistent with the expected role of *ϵ* in regulating the strength of regularization in the psychometric field.

**Table S7.**
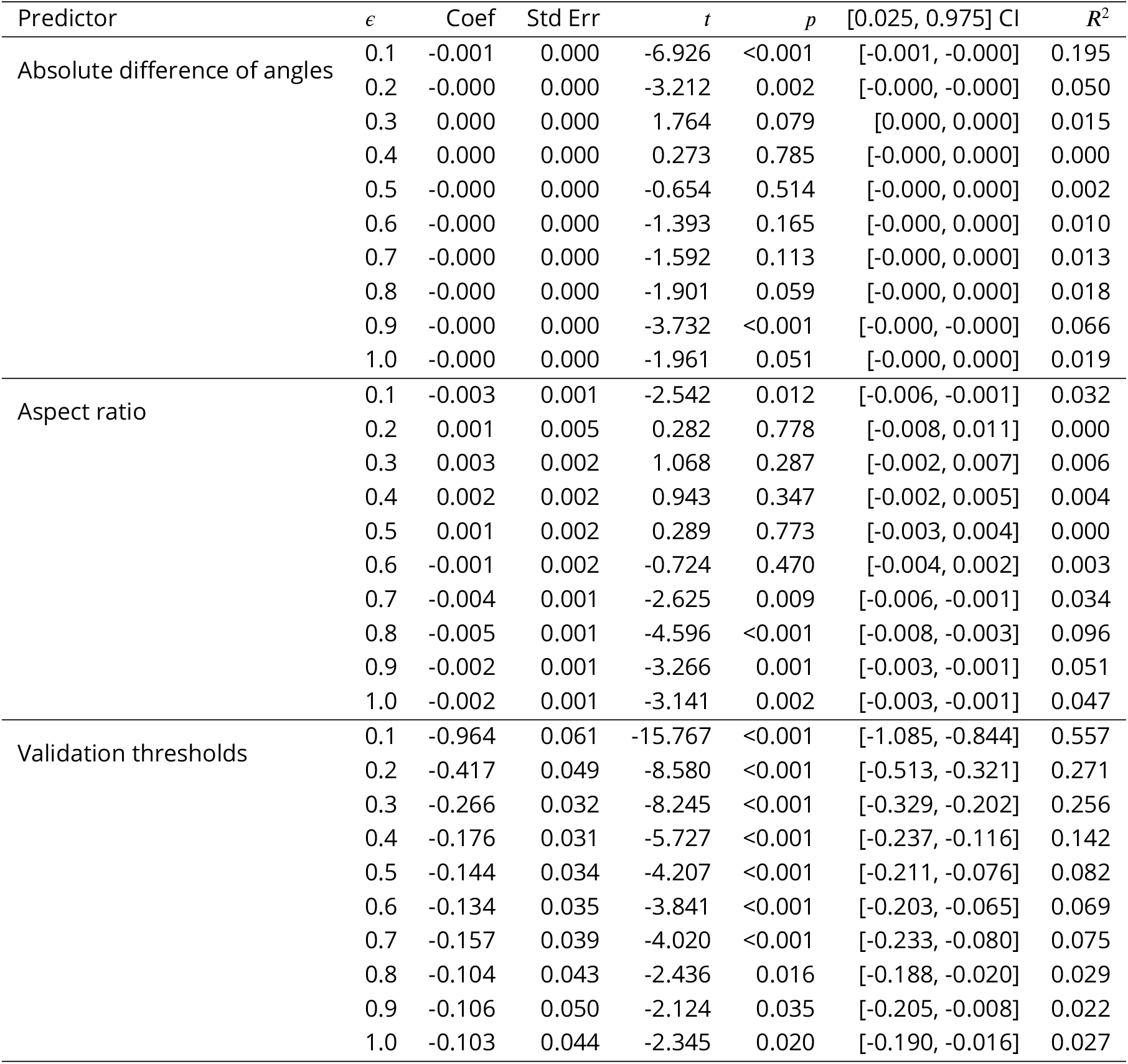
The slope of linear regression assessing the relationship between WPPM–validation threshold residuals and three predictors reported in ***Table S3***. The hyperparameter γ was fixed at 0.0003, while *ϵ* was varied.

| Predictor | $\epsilon$ | Coef | Std Err | $t$ | $p$ | [0.025, 0.975] CI | $R^2$ |
| --- | --- | --- | --- | --- | --- | --- | --- |
| Absolute difference of angles | 0.1 | -0.001 | 0.000 | -6.926 | <0.001 | [-0.001, -0.000] | 0.195 |
|  | 0.2 | -0.000 | 0.000 | -3.212 | 0.002 | [-0.000, -0.000] | 0.050 |
|  | 0.3 | 0.000 | 0.000 | 1.764 | 0.079 | [0.000, 0.000] | 0.015 |
|  | 0.4 | 0.000 | 0.000 | 0.273 | 0.785 | [-0.000, 0.000] | 0.000 |
|  | 0.5 | -0.000 | 0.000 | -0.654 | 0.514 | [-0.000, 0.000] | 0.002 |
|  | 0.6 | -0.000 | 0.000 | -1.393 | 0.165 | [-0.000, 0.000] | 0.010 |
|  | 0.7 | -0.000 | 0.000 | -1.592 | 0.113 | [-0.000, 0.000] | 0.013 |
|  | 0.8 | -0.000 | 0.000 | -1.901 | 0.059 | [-0.000, 0.000] | 0.018 |
|  | 0.9 | -0.000 | 0.000 | -3.732 | <0.001 | [-0.000, -0.000] | 0.066 |
|  | 1.0 | -0.000 | 0.000 | -1.961 | 0.051 | [-0.000, 0.000] | 0.019 |
| Aspect ratio | 0.1 | -0.003 | 0.001 | -2.542 | 0.012 | [-0.006, -0.001] | 0.032 |
|  | 0.2 | 0.001 | 0.005 | 0.282 | 0.778 | [-0.008, 0.011] | 0.000 |
|  | 0.3 | 0.003 | 0.002 | 1.068 | 0.287 | [-0.002, 0.007] | 0.006 |
|  | 0.4 | 0.002 | 0.002 | 0.943 | 0.347 | [-0.002, 0.005] | 0.004 |
|  | 0.5 | 0.001 | 0.002 | 0.289 | 0.773 | [-0.003, 0.004] | 0.000 |
|  | 0.6 | -0.001 | 0.002 | -0.724 | 0.470 | [-0.004, 0.002] | 0.003 |
|  | 0.7 | -0.004 | 0.001 | -2.625 | 0.009 | [-0.006, -0.001] | 0.034 |
|  | 0.8 | -0.005 | 0.001 | -4.596 | <0.001 | [-0.008, -0.003] | 0.096 |
|  | 0.9 | -0.002 | 0.001 | -3.266 | 0.001 | [-0.003, -0.001] | 0.051 |
|  | 1.0 | -0.002 | 0.001 | -3.141 | 0.002 | [-0.003, -0.001] | 0.047 |
| Validation thresholds | 0.1 | -0.964 | 0.061 | -15.767 | <0.001 | [-1.085, -0.844] | 0.557 |
|  | 0.2 | -0.417 | 0.049 | -8.580 | <0.001 | [-0.513, -0.321] | 0.271 |
|  | 0.3 | -0.266 | 0.032 | -8.245 | <0.001 | [-0.329, -0.202] | 0.256 |
|  | 0.4 | -0.176 | 0.031 | -5.727 | <0.001 | [-0.237, -0.116] | 0.142 |
|  | 0.5 | -0.144 | 0.034 | -4.207 | <0.001 | [-0.211, -0.076] | 0.082 |
|  | 0.6 | -0.134 | 0.035 | -3.841 | <0.001 | [-0.203, -0.065] | 0.069 |
|  | 0.7 | -0.157 | 0.039 | -4.020 | <0.001 | [-0.233, -0.080] | 0.075 |
|  | 0.8 | -0.104 | 0.043 | -2.436 | 0.016 | [-0.188, -0.020] | 0.029 |
|  | 0.9 | -0.106 | 0.050 | -2.124 | 0.035 | [-0.205, -0.008] | 0.022 |
|  | 1.0 | -0.103 | 0.044 | -2.345 | 0.020 | [-0.190, -0.016] | 0.027 |

**Figure S32.**
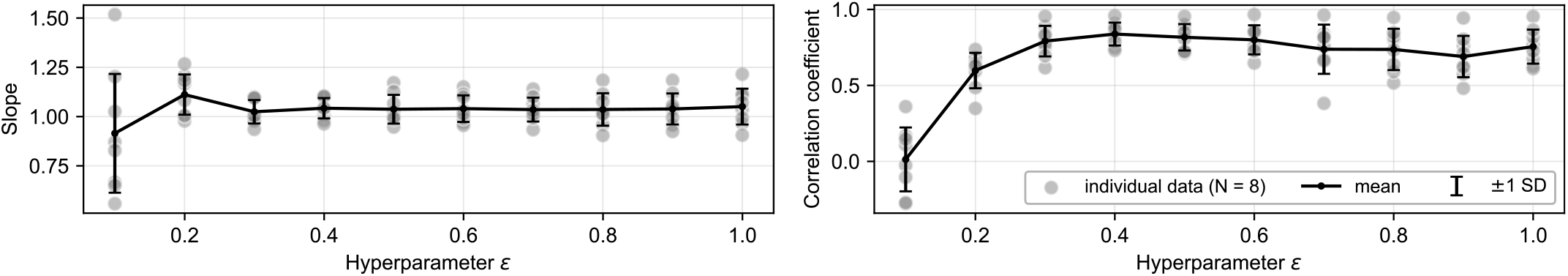
Effects of *ϵ* on the slope and correlation coefficient of the linear regression between WPPM-predicted thresholds and validation thresholds.

### Appendix 10.3: The effects of *ϵ* on linear regression between WPPM and validation thresholds

We also examined how *ϵ* affects the linear regression between WPPM and validation thresholds computed at the 25 validation conditions, as illustrated in ***Figure S6***C–***Figure S13***C. Across participants (N = 8), the mean regression slope did not deviate systematically from zero. However, the standard deviation of the slope across participants was large when *ϵ* was small (strong smoothness prior), indicating substantial bias at the individual level. The standard deviation of the slope across participants reached a minimum at *ϵ* = 0.4 and increased again for larger values of *ϵ*. Additionally, the correlation coefficient between WPPM and validation thresholds was near zero under strong smoothness, peaked at *ϵ* = 0.4, and declined again as *ϵ* increased further (***Figure S32***).

Together, these results reflect a bias–variance tradeoff. Excessive smoothness introduces bias by failing to capture structure in the data, whereas insufficient smoothness increases variance in model predictions through overfitting. These results further support our choice of *ϵ* = 0.4 as lying near the optimal balance between bias and variance.

## Appendix 11 Display characterization

### Appendix 11.1: Calibration of monitor output

**Figure S33.**
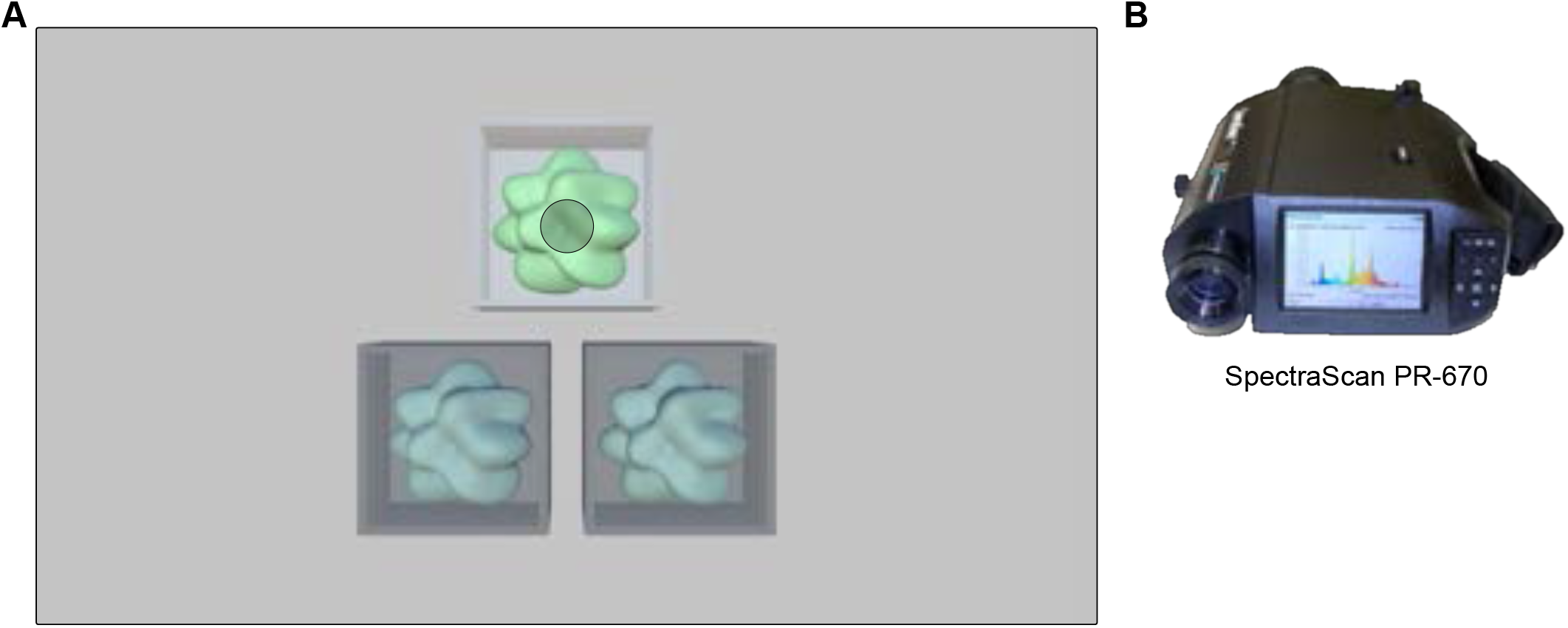
Stimuli and equipment used for calibration. (A) The stimulus setup during calibration was identical to that used in the main experiment. The surface color of both the cubic room and the blobby stimulus (shown here as the top-position stimulus) was varied during the calibration procedure. The shaded gray circular region on the stimulus indicates the area measured by the spectroradiometer’s lens. (B) A SpectraScan PR-670 used for all calibration measurements.

Calibration was carried out with three blobby objects arranged in a triangular configuration inside the cubic room (***Figure S33***A). A SpectraScan PR-670 radiometer (***Figure S33***B), positioned at the same viewing distance as the chin-rest, recorded all measurements (***Brainard et al., 2002***).

We first obtained each primary’s gamma function by measuring the screen output at 61 evenly spaced input levels (from 0 to 1) rendered through Unity (v2022.3.24f1) (***Figure S34***A). The resulting curves lie above the identity line because Unity internally applies its own assumed gamma exponent when texture values are altered. We also measured the spectral power distributions (SPDs) of the red, green, and blue primaries given different intensity levels (***Figure S34***B), and examined the stability of the primaries’ chromaticity in the CIE diagram (***Figure S34***C). There was almost no drift in the chromaticities, indicating the monitor’s color output remained stable across intensity levels (***Figure S34***D). To evaluate linearity and repeatability, we compared nominal (predicted) versus measured luminance and chromaticity for two independent measurement runs (***Figure S34***E). Deviations from linearity were minimal and nearly identical across repeats, confirming reliable reproduction (***Figure S34***F). Finally, we tested whether the cubic room’s background color affected the stimulus SPD; no measurable influence was detected (***Figure S34***G).

To assess consistency across different locations on the screen, we conducted the same set of measurements on each of the three blobby stimuli, and compared their primaries and chromaticities. The results showed consistent color behavior of the monitor (***Figure S35***), and thus we applied a single gamma correction curve to all three stimuli. This correction was derived from measurements of the bottom-right blobby stimulus. Specifically, we interpolated a gamma table for 4,096 RGB input values using a combination of linear and polynomial fits, from which we derived an inverse gamma function (***Figure S36***A). To validate this correction, we repeated the measurements with the gamma correction applied in Unity. The measured output closely aligned with the identity line across all three primaries, indicating accurate correction (***Figure S36***B).

**Figure S34.**
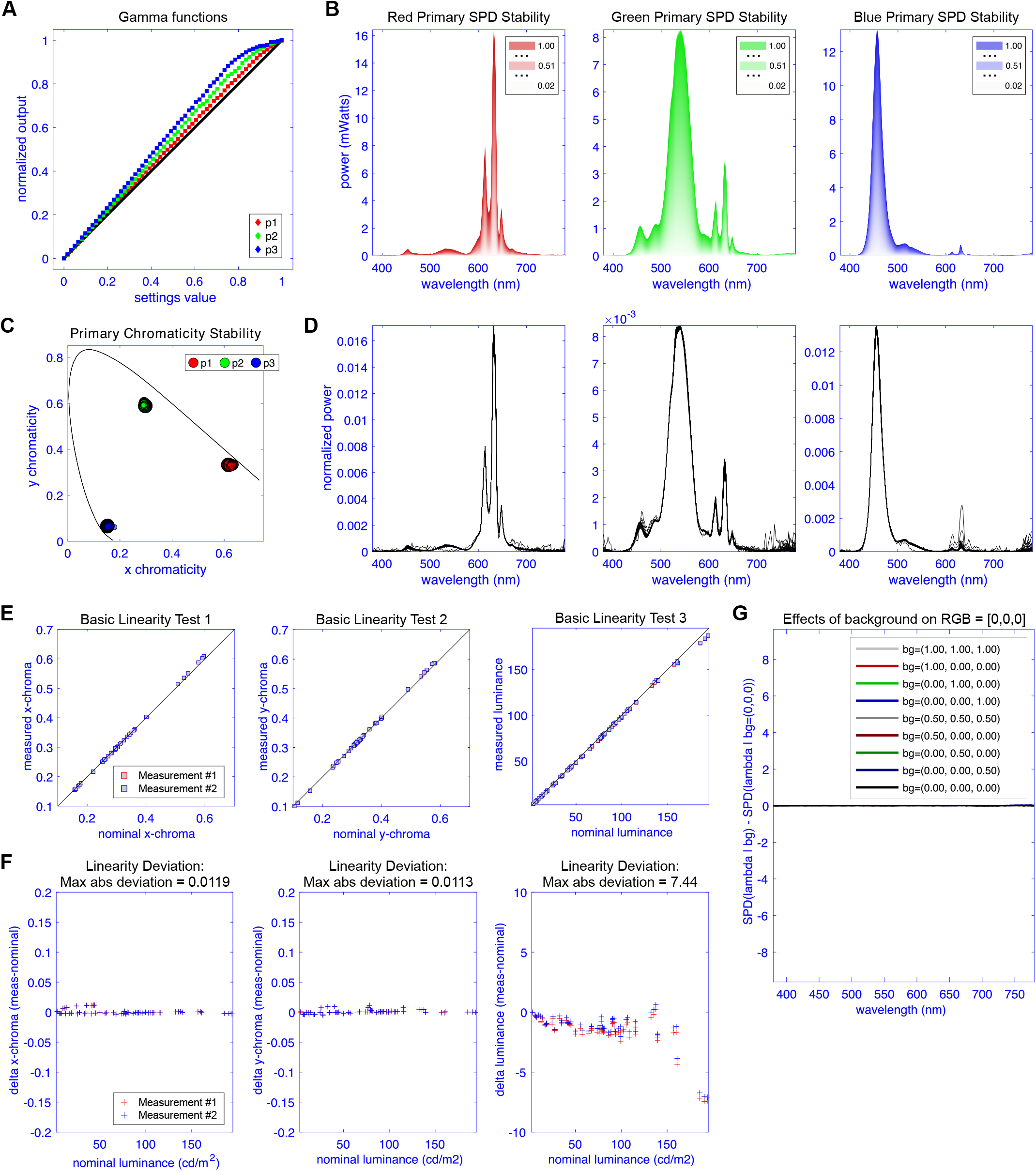
Characterization of display output via Unity’s rendering pipeline. (A) Gamma functions for red, green and blue primaries. Note that Unity’s internal correction places them above the identity line. (B) Spectral power distributions (SPDs) of the three primaries across a range of intensity levels. (C) The chromaticity of each primaries in the CIE chromaticity diagram at different intensity levels. (D) Normalized SPDs for each primary, showing that SPD shape is stable across intensity levels. (E) Linearity tests comparing predicted and measured chromaticity and luminance across two independent measurement runs. (F) Deviations from linearity. (G) Effect of the cubic room’s background color on the SPD of the blobby stimulus, showing no detectable influence.

**Figure S35.**
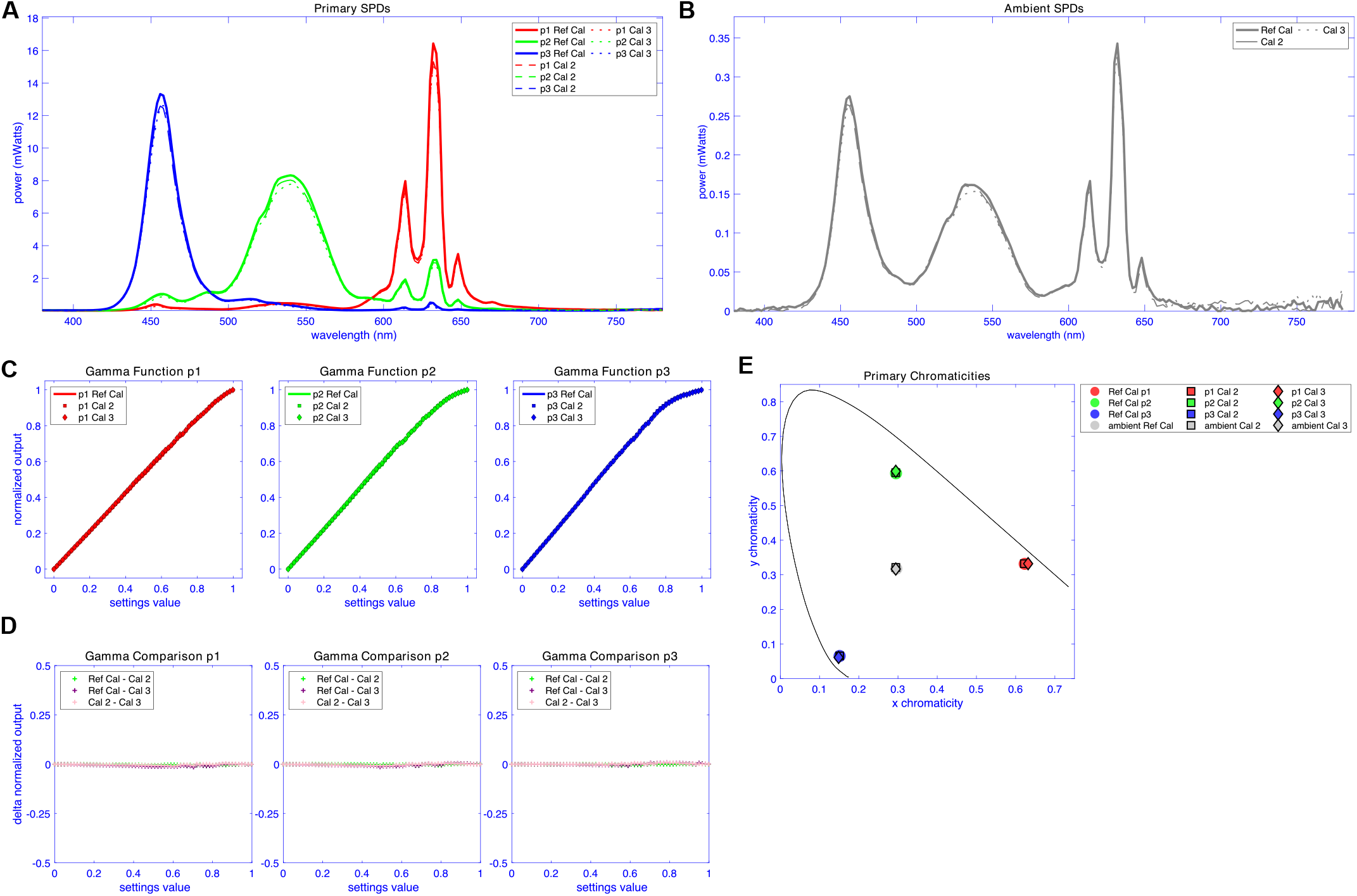
Comparison of display output across stimulus locations. (A) Spectral power distributions (SPDs) for each stimulus location: Ref Cal (bottom right), Cal 2 (bottom left), and Cal 3 (top). (B) Ambient light SPDs measured during calibration. (C) Gamma functions for each primary (red, green, blue) across all three stimulus locations. (D) Differences in normalized output for each pairwise comparison of stimulus locations, plotted separately for each primary. (E) Chromaticity coordinates of each primary in the CIE diagram, shown for all three stimulus locations.

Finally, to ensure the gamma correction remained stable over time, we repeated the measurements of display output via Unity’s rendering pipeline on the bottom-right stimulus with gamma correction applied, approximately one month after data collection began. The results confirmed that the correction remained accurate and consistent (***Figure S37***).

### Appendix 11.2: Assessment of color depth

Color depth measurements were conducted using a single blobby stimulus positioned at the center of the screen (***Figure S38***A). This stimulus was originally the top stimulus in the triangular configuration, and the camera view was adjusted to center it on the screen. Additionally, compared to the scene used in the main experiment, the other two blobby stimuli and all cubic room elements were excluded from rendering and thus were not visible. A Klein K-10A colorimeter (***Figure S38***A), placed directly in front of the monitor without any distance, was used to make the measurements.

Specifically, we tested RGB values ranging from 1928/4095 to 2128/4095, in increments of 1/4095. Each stimulus was displayed for 5 seconds, and the RGB values from the first frame of the frame buffer were saved in EXR format. We then compared the average RGB values across the surface of the blobby object (extracted from the EXR files) to the luminance measured throughout the full stimulus presentation. Although individual pixels exhibited quantization below 12-bit precision, the mean luminance increased with each 1/4095 increment, rather than in a staircase pattern. A similarly smooth progression was observed in the average R, G, and B channel values, with the R channel shown as an example in ***Figure S38***B.

**Figure S36.**
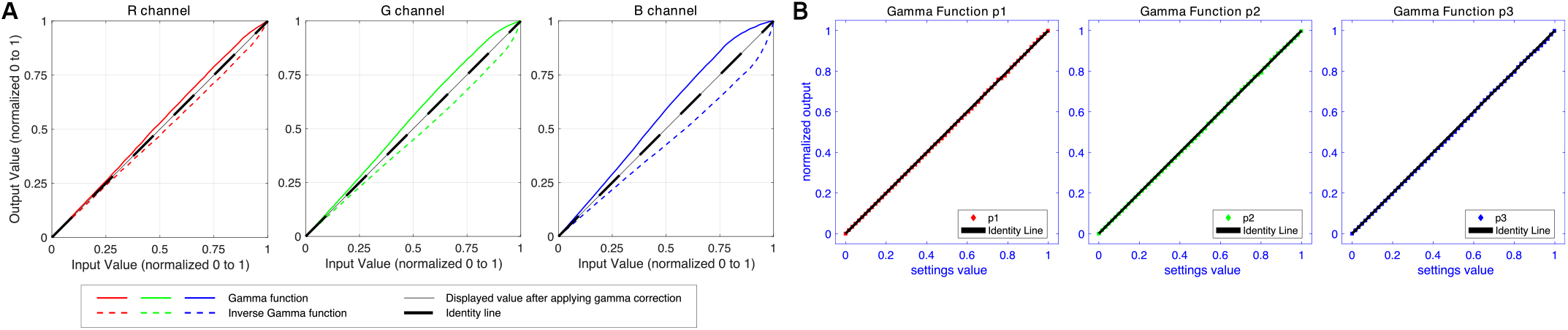
Gamma correction. (A) Measured gamma functions and their corresponding inverse functions for the red, green, and blue primaries, used to construct the gamma correction lookup table. (B) Gamma functions re-measured after applying the correction in Unity, showing close alignment with the identity line for all three primaries.

**Figure S37.**
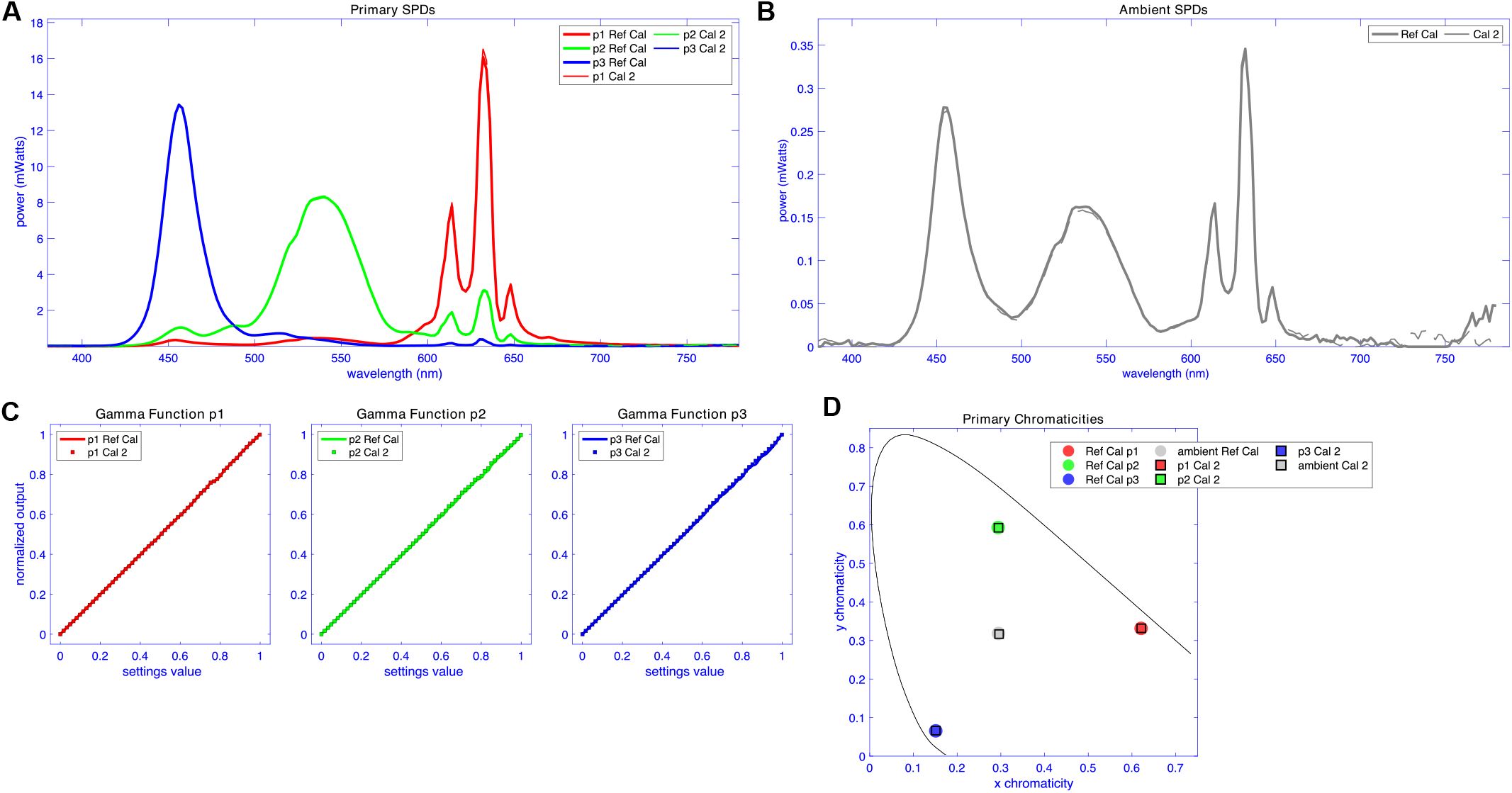
Comparison of display output with gamma correction over time. (A) Spectral power distributions (SPDs) when measured at the bottom-right blobby stimulus location. Ref Cal (initial measurements prior to the experiment) and Cal 2 (follow-up measurements roughly one month after data collection began). (B) Ambient light SPDs measured during each calibration. (C) Gamma functions for the red, green, and blue primaries across both sessions, with gamma correction applied. (D) Chromaticity coordinates of each primary plotted in the CIE diagram for both calibration runs.

To better understand how Unity and our video chain achieved this behavior, we analyzed horizontal slices of pixel values from the EXR files. When extracting a very thin slice—just one pixel in height—the individual pixel values exhibited staircase-like changes, consistent with 8-bit quantization. However, as we increased the height of the horizontal slice, the averaged channel values became progressively smoother. These results suggest that Unity achieves effective 12-bit color depth through internal spatial dithering (***Figure S38***C).

**Figure S38.**
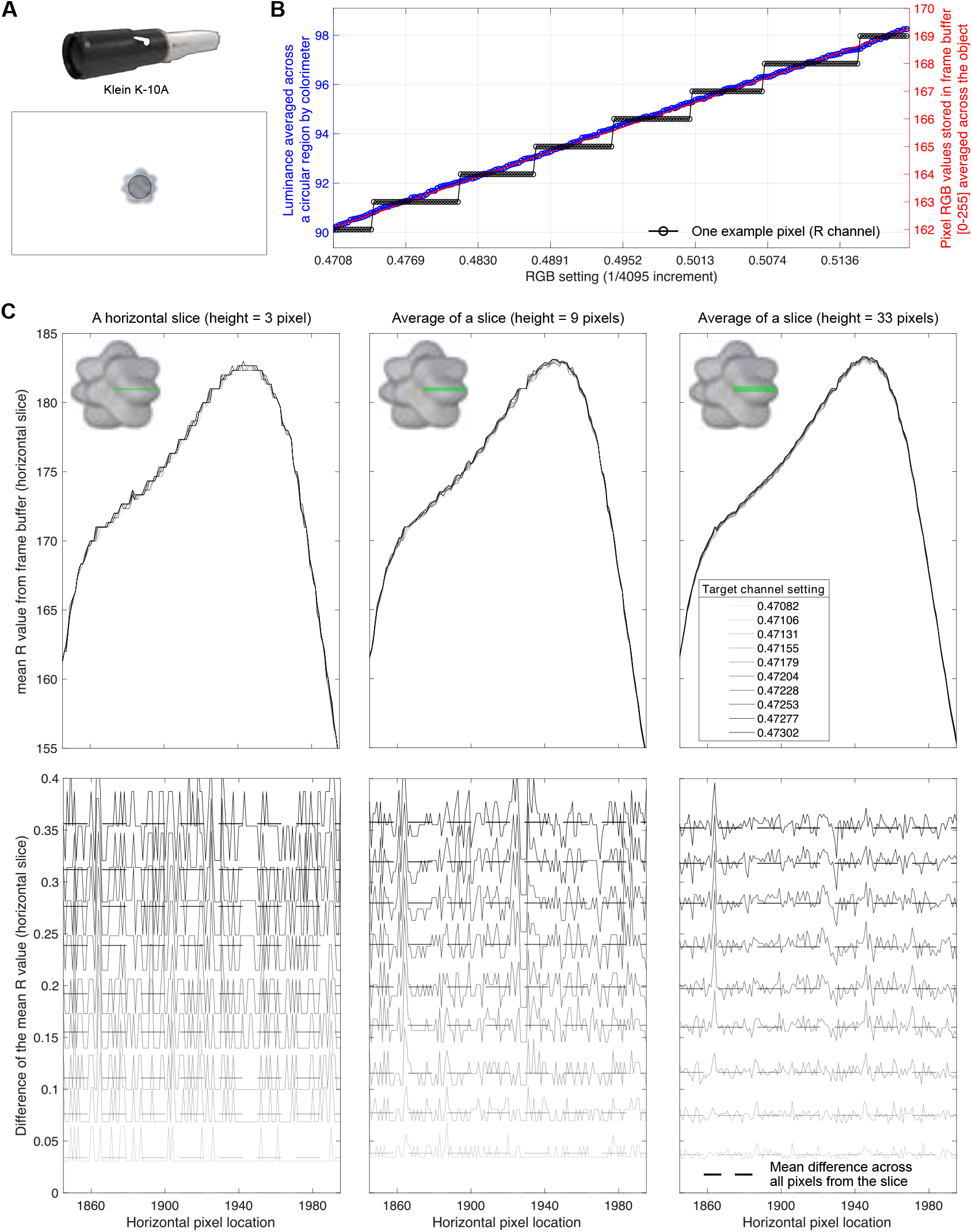
Evidence of spatial dithering by Unity’s rendering pipeline. (A) The stimulus setup during measurements was identical to that used in the main experiment. The surface color of both the cubic room and the blobby stimulus (shown here as the top-position stimulus) was varied across trials during the measurements. The shaded gray circular region on the stimulus indicates the area measured by the colorimeter’s aperture. (B) Spatial dithering by Unity’s standard shader is suggested by comparing the luminance measurements from the Klein K-10A (averaged across a circular region on the blobby object) with the RGB values stored in the frame buffer. The measured luminance shows small incremental changes as the RGB settings increase in steps of 1/4095. These measurements are consistent with what we obtain by averaging over pixels in a saved image of the frame buffer (saved from Unity in .exr format). The averaged pixel values exhibit 12-bit quantization even though individual pixel values exhibit 8-bit quantization. (C) Top row: mean R channel values averaged vertically within a horizontal slice of the blobby object. Bottom row: differences in the R channel values between the minimum target R channel setting and each of the rest settings. Different shades of gray represent different target R settings. For illustration, only a portion of the horizontal slice is shown, and solid lines in the bottom row are scaled by a factor of 0.1. Dashed lines: the mean difference averaged across all pixels within each slice.

## Appendix 12 Differentiable Monte Carlo approach

Recall from the main text that the log likelihood function implied by the WPPM observer model can be written in terms of

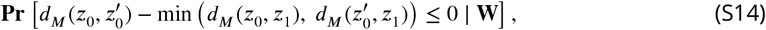

which has no simple closed form solution and must be estimated by Monte Carlo simulation. This quantity of interest takes the form of a cumulative distribution function *g*(*u*) = Pr[*v* ≤ *u*] for some random variable *v* and scalar constant *u*. In this section, we describe how to approximate the log likelihood in a manner that is compatible with automatic differentiation libraries, which enables gradient-based optimization of the log posterior density.

Let *P*_*θ*_ denote some probability distribution parameterized by *θ*. Given *n* independent and identically distributed random variables, *v*_1_, …, *v*_*n*_ ∼ *P*_*θ*_, we would like to form an estimate of the cumulative distribution function, *g*(*u*) = Pr[*v* ≤ *u*] where *v* ∼ *P*_*θ*_. A simple and well-known estimate is empirical cumulative distribution function:

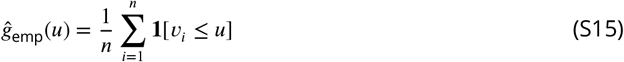

where **1**[⋅] is the indicator function—i.e. **1**[*A*] evaluates to one if the event *A* occurs and evaluates to zero otherwise. In many respects, this is a perfectly fine estimator. For example, the celebrated Dvoretzky–Kiefer–Wolfowitz inequality (***Dvoretzky et al., 1956***) states that this estimate converges exponentially fast to the true cumulative distribution function as *n* → ∞.

In our setting, we would like to not only evaluate *g*(*u*) for any given *u*, but to also evaluate ∂*g*(*u*)/∂*θ*_*j*_ for all parameters *θ*_1_, …, *θ*_*J*_ that define the underlying distribution *P*_*θ*_. ***Equation S15*** provides an estimate of *g*(*u*) but it is unfortunately not differentiable with respect to *v*_1_, …, *v*_*n*_ because **1**[*v*_*i*_ ≤ *u*] is a discontinuous step as a function of *u*. A straightforward and intuitive solution is to replace this step function with a smooth sigmoid function. We formalize this approach below, showing that it can be motivated by forming a smoothed estimate of the underlying density function.

Specifically, let *K*(*v*) denote a smooth, nonnegative function that integrates to one and satisfies *K*(*v*) = *K*(−*v*). Suppose that *P*_*θ*_ has a density function *f* (*v*). Then, given *v*_1_, …, *v*_*n*_ ∼ *P*_*θ*_ we can estimate the density function as:

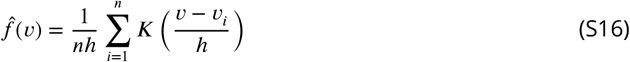

where *h* > 0 is a user-specified hyperparameter called the bandwidth. ***Equation S16*** is known as a *kernel density estimate* (***Wasserman, 2006***). Asymptotically, 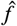 approaches the true density function

*f* as *n* → ∞ and *h* → 0. Intuitively, larger values of *h* lead to smoother density estimates, which is preferable in sample-limited (i.e. small *n*) regimes.

Now define:

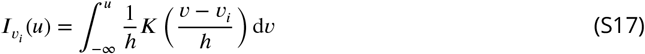

which is a smooth sigmoid function centered at *v*_*i*_, and consider the following estimate of the cumulative distribution function:

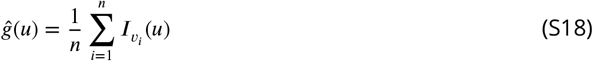

Notice that in the limit of *h* → 0, we recover the empirical cumulative distribution estimator *ĝ* = *ĝ*_emp_ because, in this limit, we have that 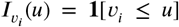. We can further justify ***Equation S18*** as a reasonable estimator of *g* by recognizing it as the integral of the density estimate in ***Equation S16***. That is,

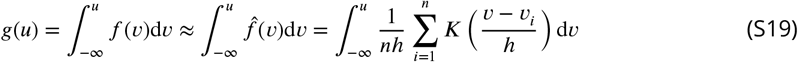

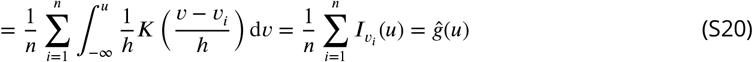

Our refined estimator ĝ is clearly differentiable whenever we choose *K*(*v*) to be a smoothly differentiable function. In our model fitting routine, we chose *K*(*v*) to be the density of a standard logistic distribution:

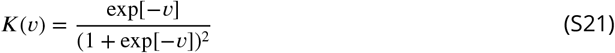

This smoothing kernel has heavy tails, which we reasoned would enable numerically stable autodifferentiation routines even when *h* is chosen to be small. Another feature is that the integrated density is the well-known *logistic function*:

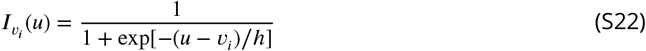

which is familiar and easy to compute.

